# 10-Nitro-oleic acid improves survival following gamma irradiation and modulates hematopoietic and intestinal responses

**DOI:** 10.64898/2026.09.25.754470

**Authors:** Jana Pereckova, Rachel Vichova, Jiri Toufar, Daniel Chrenko, Jiří Pospíšil, Tomas Perecko

## Abstract

Acute radiation syndrome remains a major consequence of high-dose ionizing radiation exposure, and effective pharmacological countermeasures are limited. Nitro-fatty acids possess anti-inflammatory and cytoprotective properties, suggesting potential effectiveness in radiation injury. Here, we investigated the radioprotective effects of 10-nitro-oleic acid (10-NO₂OA) in mouse models of total body irradiation (TBI).

C57BL/6 mice were exposed to either sublethal (4 Gy) or LD_50_-level (7.5 Gy) gamma irradiation and treated with 10-NO₂OA according to pre-irradiation, post-irradiation, or combined treatment regimens. Survival, hematological responses, bone marrow cells (BMCs)/colony-forming units (CFU), plasma granulocyte colony-stimulating factor (G-CSF), NRF2-responsive signaling, and intestinal responses were evaluated.

Administration of 10-NO₂OA before and after irradiation significantly improved 30-day survival following 7.5 Gy TBI (57% vs. 20% in vehicle-treated controls). In contrast, post-irradiation treatment alone was ineffective, whereas pre-irradiation treatment alone produced a non-significant trend toward improved survival. The protective effect was absent in BALB/c mice, indicating strain dependence. Although 10-NO₂OA did not prevent long-term radiation-induced cytopenia, it increased neutrophil and platelet counts, elevated plasma G-CSF levels shortly after irradiation, and transiently enhanced BMC counts. In irradiated BMCs, 10-NO₂OA activated NRF2-responsive gene expression *in vitro* and *in vivo*. Exploratory intestinal analyses revealed improved macroscopic appearance and partial restoration of irradiation-suppressed cytokine and growth factor production.

In conclusion, 10-NO₂OA improves survival following LD_50_-level gamma irradiation and is associated with modulation of hematopoietic, cytoprotective, and intestinal responses. These findings identify 10-NO₂OA as a promising candidate radioprotective agent and support a multifactorial mechanism of action.

## Introduction

Acute radiation syndrome (ARS) remains a major consequence of high-dose ionizing radiation exposure, whether in accidental settings, radiological incidents, or intensive therapeutic regimens involving total body irradiation (TBI) [1]. Its clinical course is determined by dose, timing, and the vulnerability of individual tissues, with the hematopoietic and gastrointestinal systems representing two of the most sensitive targets [2]. The human hematopoietic system is highly sensitive to gamma irradiation, showing minor acute damage at 0.5–1 Gy and early bone marrow injury at 1–2 Gy [3]. In humans, hematopoietic ARS (H-ARS) can develop at doses as low as approximately 2.5 Gy, with severity increasing at higher doses; the estimated LD_50_ is about 3.5–4 Gy without supportive care and 4.5–7 Gy with intensive medical management. Gastrointestinal ARS generally emerges at around 6 Gy and is typically lethal at doses ≥10 Gy [2]. Damage to these compartments disrupts immune defense, tissue renewal, and systemic homeostasis, making them central determinants of survival [1].

Current clinical management of ARS is centered largely on supportive care and, for hematopoietic injury, on cytokine-based therapies that accelerate marrow recovery [1, 4]. In H-ARS, granulocyte colony-stimulating factor (filgrastim and pegfilgrastim) and granulocyte–macrophage colony-stimulating factor (sargramostim) are used to stimulate proliferation and differentiation of myeloid progenitors and to shorten neutropenia, whereas romiplostim, a thrombopoietin receptor agonist, promotes megakaryopoiesis and platelet recovery [1, 4]. Importantly, these interventions mainly address H-ARS. Approved therapies that specifically target gastrointestinal radiation injury remain lacking [2]. Although supportive care and selected hematopoietic growth factors can improve outcome in some settings, effective pharmacological agents that provide broader protection against radiation injury remain limited [1].

In this context, endogenous lipid mediators with anti-inflammatory and cytoprotective activity have emerged as attractive candidates for radiation countermeasure development. Nitro-fatty acids are electrophilic lipid derivatives generated during inflammatory and nitrative reactions and are increasingly recognized as modulators of cellular stress responses [5-7]. Among them, 10-nitro-oleic acid (10-NO_2_OA) is of particular interest because it can influence redox-sensitive and inflammatory signaling pathways implicated in radiation injury, including nuclear factor erythroid 2-related factor 2 (NRF2)-associated cytoprotective responses and suppression of pro-inflammatory cascades [8-10]. These pleiotropic properties suggest that 10-NO_2_OA may not simply reduce oxidative damage but could more broadly shape the tissue response to irradiation [6, 10].

We previously showed that a mixed 9/10-NO_2_OA isomer improves early hematopoietic recovery after sublethal TBI, increasing bone marrow cellularity and supporting recovery of circulating leukocytes [11]. Further, we found that 10-NO_2_OA effectively activates the NRF2 signaling pathway in primary bone marrow cells under physiologically relevant oxygen conditions [8]. These findings raised the possibility that 10-NO_2_OA may act as a radioprotective agent; however, several key questions remained unresolved. It is unclear whether these early hematopoietic effects would translate into improved survival after a more severe, LD_50_-level gamma irradiation, whether protection would extend beyond the hematopoietic compartment, and whether the underlying mechanism would be adequately explained by canonical NRF2-responsive signaling alone.

In the present study, we examined the radioprotective potential of 10-NO_2_OA in mouse models of TBI encompassing both the hematopoietic and gastrointestinal components of ARS. We assessed the influence of treatment timing on survival and evaluated the effects of 10-NO_2_OA on peripheral blood and bone marrow responses, early molecular alterations in irradiated bone marrow cells, including activation of the NRF2 pathway, and intestinal injury following LD_50_ irradiation. By integrating these endpoints, we sought to determine whether the beneficial effects of 10-NO_2_OA are mediated primarily through preservation of hematopoiesis, protection of the gastrointestinal tract, or a combination of both.

## Materials and methods

### Nitro-fatty acids

Nitro-fatty acids (NO₂FAs), namely (*E*)-10-nitro-oleic acid (10-NO₂OA), (*E*)-9-nitro-oleic acid (9-NO₂OA), (*E*)-10-nitro-linoleic acid (10-NO₂LA), (*E*)-9-nitro-conjugated linoleic acid (9-NO₂cLA), and 10-nitro-stearic acid (10-NO₂SA) (Fig. 6E), were prepared using our previously developed regio- and stereoselective synthetic methods [8, 12]. All NO₂FAs were obtained with >98% regioisomeric and stereochemical purity, and their identity was confirmed by spectroscopic and analytical methods. Detailed synthetic procedures, purification methods, and characterization data are provided in the Supplementary file 2_Synthesis.

### Animals

Male C57/BL6 and BALB/c mice aged 10–12 weeks and weighing 25–30 g were purchased from AnLab, Czech Republic. Mice were housed in groups of 5 animals per cage under standard conditions at a room temperature of 22±2 °C, with food and water provided *ad libitum*. Mice were maintained on a twelve-hour light-dark schedule (500 lux). Experiments were initiated after 7 days of quarantine. All animal experiments were conducted in accordance with EU Directive 2010/63/EU and Czech Act No. 246/1992 Coll., on the Protection of Animals Against Cruelty and Decree No. 419/2012 Coll., on the Protection of Experimental Animals. Animal experiments were approved by the Resort Professional Commission of the Czech Academy of Sciences for Approval of Projects of Experiments on Animals (AVCR 1823/2023 SOV II), supervised by the institutional Animal Welfare Body of the Institute of Biophysics of the Czech Academy of Sciences, and performed by certified personnel. Experiments with 3–6 mice per group were repeated in at least two independent trials. The total numbers of animals used in each experimental setting is provided in the corresponding figure legends.

BALB/c mice were included in a complementary survival study to assess the strain dependence of the radioprotective effects of 10-NO_2_OA.

### 10-NO2OA administration

The mice were randomized into three groups for mechanistic and hematological studies: the control (vehicle; no irradiation), TBI (total body irradiation) + vehicle, and TBI + 10-NO_2_OA. Survival studies were conducted only in irradiated animals and included the TBI + vehicle and TBI + 10-NO_2_OA groups. Mice received intraperitoneal injections of either vehicle (PBS + 10% ethanol) or 10-NO_2_OA (in PBS + 10% ethanol) at a dose of 10 mg/kg body weight in a volume of 0.1 ml. For survival studies, mice received either four doses (24 and 1 h before irradiation and 4 and 48 h after irradiation), two pre-irradiation doses (24 and 1 h before irradiation), or two post-irradiation doses (4 and 48 h after irradiation). For hematological studies, mice received either four doses (24 and 1 h before irradiation and 4 and 48 h after irradiation), or two pre-irradiation doses (24 and 1 h before irradiation). In selected experiments, mice receiving 10-NO_2_OA without irradiation were also analyzed. In mechanistic studies, additional dosing regimens and sample collection time points were used as appropriate for individual experimental endpoints; these are described in the corresponding Methods sections and summarized in Scheme 1.

**Scheme 1.**
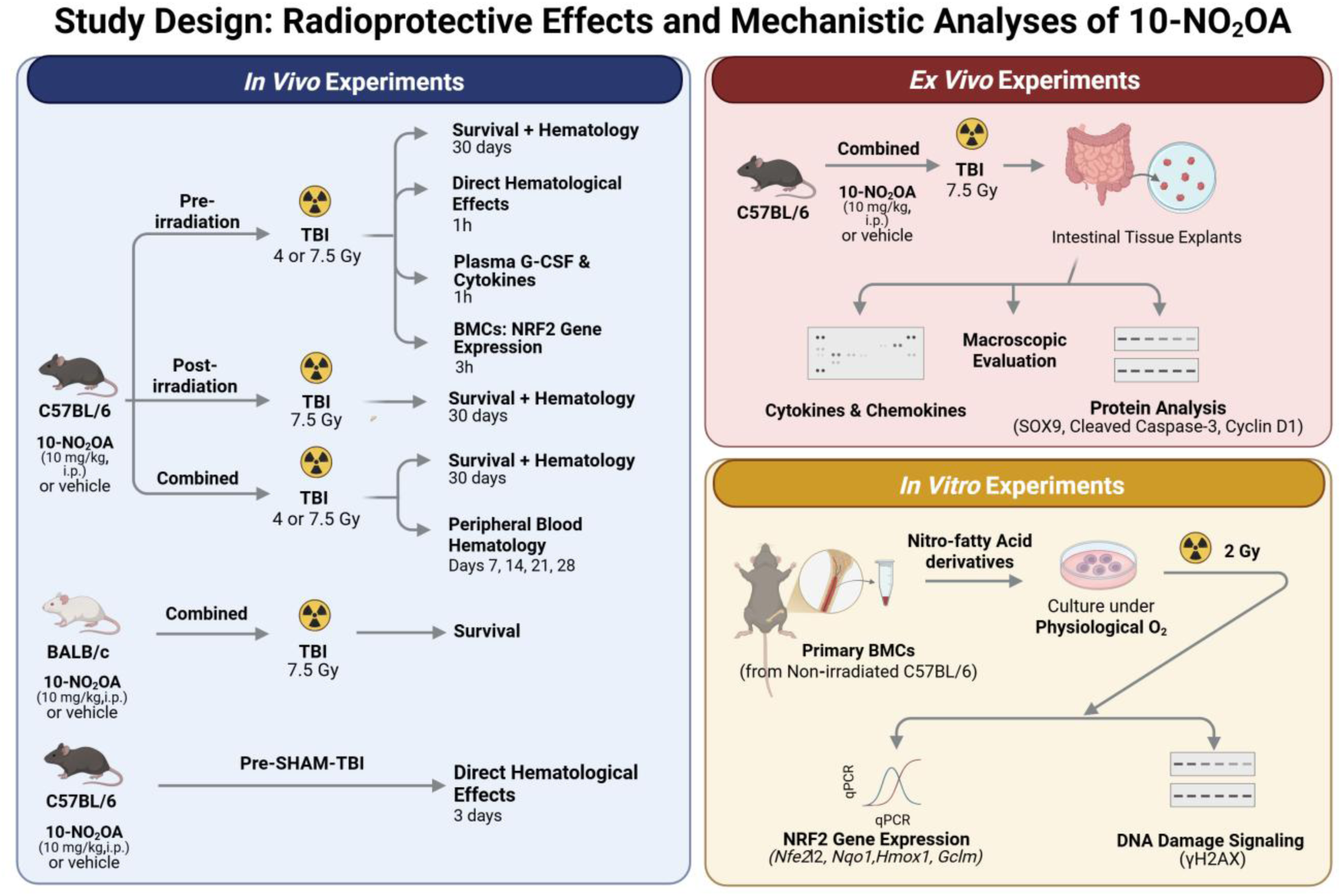
Overview of the experimental design and 10-NO_2_OA treatment regimens. The study comprised three complementary experimental approaches: *in vivo*, *ex vivo*, and *in vitro*. C57BL/6 mice were exposed to total body irradiation (TBI) with either 4 Gy (sublethal dose) or 7.5 Gy (LD_50/30_-level dose) and received intraperitoneal injections of 10-NO_2_OA (10 mg/kg body weight) or vehicle. The combined regimen comprised four injections administered 24 and 1 h before TBI and 4 and 48 h after TBI; schedule-dependence studies used either two pre-irradiation injections (24 and 1 h before TBI) or two post-irradiation injections (4 and 48 h after TBI). Survival and body weight were monitored for 30 days after 7.5 Gy TBI. Peripheral blood, plasma, bone marrow cells, and intestinal tissues were collected at the indicated time points for hematological, G-CSF, cytokine, gene-expression, protein, and macroscopic analyses. Selected survival experiments were also performed in BALB/c mice. Intraperitoneal (i.p.); total body irradiation (TBI); 10-nitro-oleic acid (10-NO_2_OA); bone marrow cells (BMC); granulocyte colony-stimulating factor (G-CSF); nuclear factor erythroid 2-related factor 2 (NRF2). Created in BioRender. Pereckova, J. (2026) https://BioRender.com/hvl3xom.

### Irradiation and survival experiments

Total body irradiation (TBI) of mice was performed at a dose rate of 1 Gy/min using a gamma-ray source (⁶⁰Co, Chirana, Czech Republic). Conscious mice were placed in a plastic container and exposed to a single sublethal dose of 4 Gy or an experimentally determined LD₅₀/₃₀ dose of 7.5 Gy. Control mice (vehicle-treated, non-irradiated) spent the same amount of time in the radiation facility without exposure to irradiation. For hematological studies, mice were euthanized 7, 14, 21, or 28 days after TBI, whereas survival studies were terminated 30 days after irradiation. Additional dosing schedules and sampling time points were used in selected mechanistic experiments. A schematic overview of the experimental design is provided in Scheme 1.

Survival experiments were conducted only in irradiated mice and comprised of two experimental groups: vehicle-treated and 10-NO_2_OA-treated animals. Animals in survival experiments were monitored twice daily by trained laboratory personnel for morbidity and mortality, including assessment of body weight, behavior, eye closure, food intake, and posture. These observation criteria were predefined and approved by the institutional ethics committee, and all procedures complied with the approved animal protocol. Animals in a moribund state or with a body weight loss exceeding 20% were humanely euthanized by CO_2_ inhalation followed by cervical dislocation. Survival studies were performed in C57BL/6 mice. An additional survival experiment was conducted in BALB/c mice to evaluate whether the effects of 10-NO_2_OA were strain dependent.

### Hematologic analysis

Peripheral blood was collected on days 7, 14, 21, 28, or 30 after TBI from anesthetized mice by cardiac puncture into EDTA-coated microtubes (Sarstedt, Germany). Mice were euthanized by overdose of ketamine/xylazine anesthesia (Gedeon Richter, Hungary; BioVeta, Czech Republic). In mechanistic experiments, blood was also collected at specified time after TBI. Total white blood cell (WBC) counts, WBC differential counts, lymphocytes (Lym) and neutrophils (Neu), as well as red blood cell (RBC) and platelet (PLT) counts, were determined using a BC-5300Vet hematology analyzer (Mindray, China).

### Plasma granulocyte colony-stimulating factor (G-CSF) quantification

Samples of EDTA-anticoagulated blood were centrifuged at 1,500 × g for 10 minutes at 20 °C without brake. The plasma layer was then aspirated and centrifuged at 15,000 × g for 10 minutes at 4 °C to remove residual cellular debris. Plasma levels of G-CSF were quantified using an enzyme-linked immunosorbent assay (ELISA) kit (DY414-05; R&D Systems, USA; lot P466075) according to the manufacturer’s instructions.

### Bone marrow isolation and colony-forming unit (CFU) assays

Primary bone marrow cells (BMCs) were isolated as previously described in *Chrenko et al.* [8]. Briefly, femoral BMCs were flushed into 2 ml of PBS containing 2% fetal bovine serum (FBS; FB-1101/500, Biosera, France), with cells from both femurs pooled for analysis. Following erythrocyte lysis using ACK lysing buffer (A1049201, Thermo Scientific, USA), nucleated cells were counted using a Bio-Rad cell counter (Bio-Rad, CA, USA). Granulocyte-macrophage colony-forming units (GM-CFUs) were assessed using MethoCult GF M3534 medium (03534, Stemcell Technologies, Canada). Femoral BMC suspensions were plated at 1.5 × 10^5^ cells per well in 6-well plates in triplicate and incubated at 37°C in a humidified atmosphere containing 5% CO_2_. GM-CFUs were scored after 7-10 days as colonies containing at least 50 cells. GM-CFU numbers were normalized to femur cellularity and body weight.

### Gene expression

Relative gene expression of nuclear factor erythroid 2-related factor 2 (NRF2) responsive genes: NRF2, encoded by *Nfe2l2* (Mm00477784_m1), heme oxygenase-1 (*Hmox1*; Mm00516005_m1), and Glutamate-Cysteine Ligase Modifier Subunit (*Gclm*; Mm00514996_m1), and NAD(P)H Quinone Dehydrogenase 1 (*Nqo1*; Mm01253561_m1) (all from Thermo Fisher Scientific, USA) was detected as was described in [8]. In the *in vitro* experiments (Fig. 6), BMCs (3 × 10^6^ cells) were cultured under physiological oxygen conditions (1% or 4% O_2_) and treated with 2 μM nitro-fatty acid derivatives. After 1 h, the BMCs were irradiated with 2 Gy and incubated for an additional 3 h. In the *in vivo* experiments (Fig. S4), BMCs were isolated from treated C57BL/6 mice. Specifically, mice received two doses of 10-NO_2_OA (10 mg/kg body weight) or vehicle at 24 and 1 h before TBI (4 or 7.5 Gy). One hour after TBI, BMCs were isolated under 1% O_2_ conditions. BMCs from both the *in vitro* and *in vivo* experiments were then washed with PBS and lysed using a NucleoSpin RNA Plus kit (740984, Machery-Nagel; Germany). Total RNA (0.2 μg) was reversely transcribed into first strand cDNA using a RevertAid First Strand cDNA Syntesis kit according to the manufacturer’s protocol (K1622, Thermo Fisher Scientific, USA). qRT-PCR reactions were performed in a LightCycler480 instrument (Roche; Switzerland) using TaqMan™ FastAdvanced Master Mix for qPCR (4444557, Thermo Fisher Scientific, USA). Data were normalized to ribosomal protein L32 (Rpl32, Mm02528467_g1, Thermo Fisher Scientific, USA) and presented as 2-Δcq.

### Western blot analysis

BMCs or intestinal tissue explants were lysed in RIPA buffer (89901, Thermo Scientific, USA) supplemented with protease and phosphatase inhibitor cocktail (78442, Thermo Scientific, USA) for 5 min on ice. The lysates were then sonicated twice for 5 seconds and centrifuged at 15,000 × g for 10 min at 4°C. Supernatants were collected, and protein concentrations were determined using a BCA protein assay kit (23227, Thermo Scientific, USA). Equal amounts of protein were separated on 7-15% SDS-polyacrylamide gels. Following electrophoresis, proteins were transferred onto PVDF membranes (IPVH85R, Millipore, USA) using a Pierce^TM^ Power Blot System (Thermo Scientific, USA) with Pierce^TM^ 1-Step Transfer Buffer (84731, Thermo Scientific, USA). Membranes were blocked with 5% non-fat dry milk in TBST and incubated overnight at 4°C with the following primary antibodies: anti-phospho-H2AX (9718, 1:1,000), anti-SOX9 (82630, 1:500), anti-cyclin D1 (2978, 1:1,000), anti-cleaved caspase-3 (9664, 1:500), and anti-β-actin (3700, 1:1,000) (all from Cell Signaling Technology, USA), or anti-vinculin (V9264, 1:1,000; Sigma-Aldrich, USA). Membranes were then incubated for 1 h at room temperature with the appropriate HRP-conjugated secondary antibodies (anti-mouse IgG, 7076, 1:5,000; or anti-rabbit IgG, 7074, 1:2,000; both from Cell Signaling Technology, USA). Protein bands were visualized using enhanced chemiluminescence substrates (SuperSignal™ West Pico/Femto, 34577/34095, Thermo Scientific, USA), and images were acquired using an Amersham Imager 680 (GE Healthcare, USA). Band intensities were quantified using ImageJ v1.54p (NIH, USA) and normalized to vinculin or β-actin.

### Preparation of intestinal explant cultures for macroscopic, cytokine and protein analyses

C57BL/6 mice were subjected to 7.5 Gy TBI and treated with 10-NO_2_OA or vehicle 24 and 1 h before irradiation and 4 and 48 h thereafter (see Scheme 1). On day 10 post-TBI, the large intestine was collected for macroscopic analysis. Further, the protein analysis was performed on the small intestine segments on day 6 post-TBI. In detail, a 20 cm segment of the proximal small intestine was excised, cleared of external membranes, blood vessels, and adipose tissue, and maintained in cold PBS (2–8°C). The lumen was flushed with cold PBS, the intestine was opened longitudinally and washed three times. The tissue was transferred to fresh cold PBS, cut into approximately 2 mm pieces, and repeatedly rinsed by gentle pipetting followed by gravity sedimentation until the supernatant was clear. The washed tissue fragments were resuspended in RIPA buffer supplemented with inhibitors and homogenized by pipetting or passage through a needle and syringe for subsequent protein analysis. For cytokine analysis, the large intestine was collected on day 6 post-TBI and washed five times with cold PBS. Intestinal segments (3 cm in length) were gently flushed with cold PBS, opened longitudinally, and cut into 0.5 cm pieces. The tissue segments were incubated in RPMI+10% FBS with penicillin/streptomycin for 24 h. Following incubation, culture supernatants were collected and stored at −80°C until cytokine analysis.

### Cytokine array

For plasma analysis, EDTA-anticoagulated plasma samples obtained from 3–4 individual mice within each experimental group were pooled and processed according to the manufacturer’s instructions. Mice were treated with 10-NO_2_OA or vehicle 24 and 1 h before 7.5 Gy TBI and euthanized 1 h post-irradiation.

For intestinal cytokine profiling, conditioned media collected from intestinal explant cultures from 3–4 individual animals within each experimental group were pooled and analyzed. The treatment regimen and experimental endpoint for intestinal explant cultures are described in the preceding section.

Cytokine and chemokine profiling was performed using the Mouse Cytokine Array Kit, Panel A (ARY006, Biotechne, R&D Systems, USA). Membranes were imaged using an Amersham Imager 680 (GE Healthcare, USA), and spot intensities were quantified using ImageJ v1.52 (NIH, USA). Background-subtracted data were normalized to positive control spots, log-transformed, and converted to z-scores.

### Statistical analysis

Data are presented as arithmetic means ± standard deviations (SD). Significant outliers were excluded prior to analysis. Differences between two groups were evaluated using two-tailed Student’s t-test or Welch’s t-test, as appropriate. Differences among three groups were analyzed using one-way analysis of variance (ANOVA) followed by Dunnett’s or Tukey’s multiple-comparisons test, as appropriate. Survival data were analyzed using the log-rank (Mantel-Cox) test, and median survival times and hazard ratios were calculated. Statistical significance was defined as p < 0.05. Statistical analyses and graph design were performed using GraphPad Prism version 11.1 (GraphPad Software, USA; www.graphpad.com). Details of the statistical tests and numbers of animals used are provided in the corresponding figure legends.

## Results

### 10-NO₂OA improves survival after total body irradiation

To determine whether 10-NO_2_OA modulates gamma radiation-induced lethality, C57BL/6 mice were subjected to total body irradiation (TBI; 7.5 Gy) and treated with 10-NO_2_OA or vehicle 24 h and 1 h prior to irradiation, and 4 h and 48 h thereafter (Fig. 1A). Under these conditions, 10-NO_2_OA significantly increased survival compared to vehicle-treated controls (57% vs. 20%; log-rank test, p = 0.0004; hazard ratio = 2.80, 95% CI 1.50–5.19). Median survival in the vehicle-treated group was 13.5 days. Body weight analysis revealed a progressive decline in vehicle-treated animals starting around day 10 post-TBI, reaching a nadir around day 13, followed by partial stabilization in surviving mice (Fig. 1B). In contrast, 10-NO_2_OA–treated mice maintained stable body weight throughout the observed period.

**Fig. 1:**
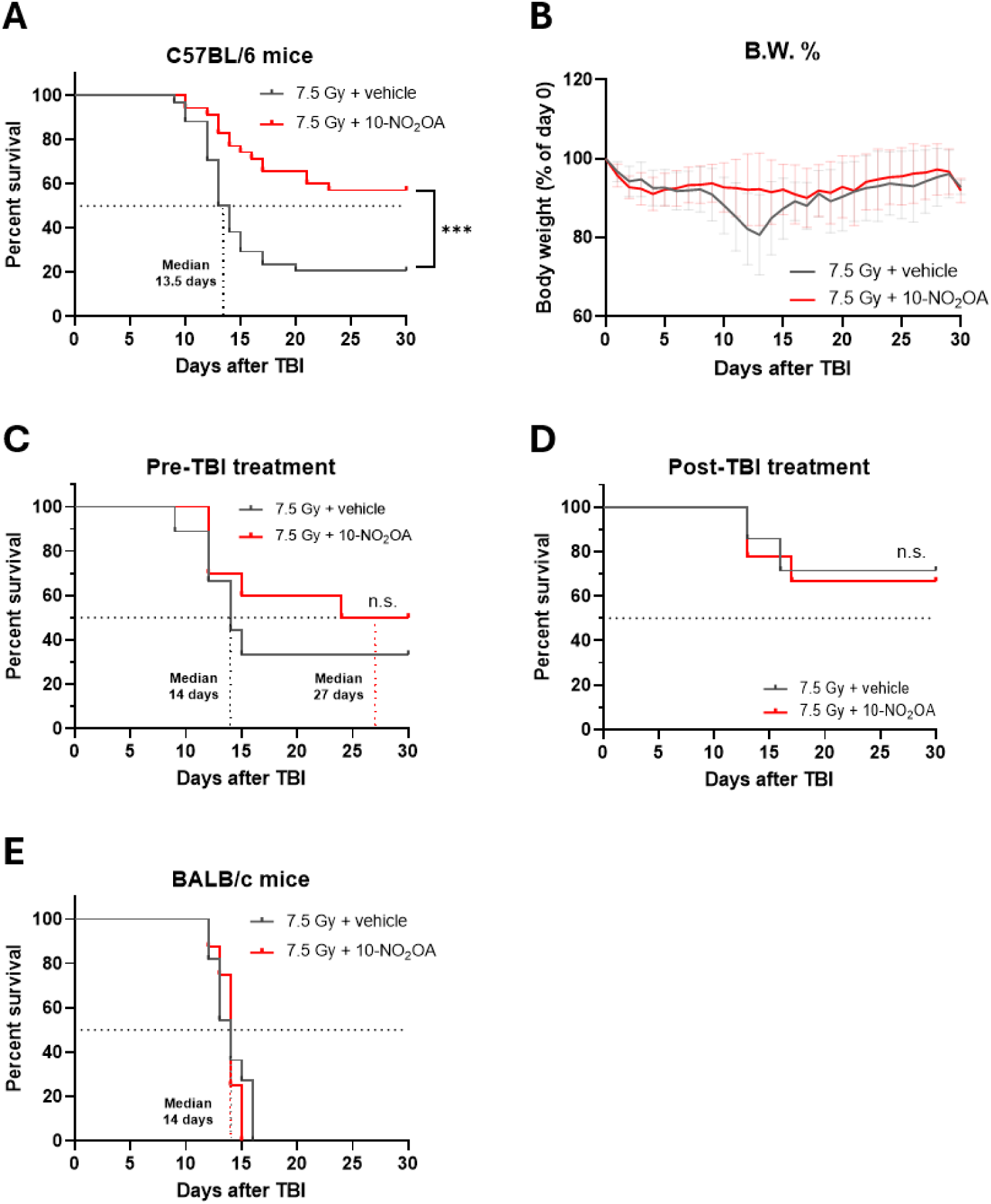
10-NO_2_OA treatment increases survival in mice after total body irradiation (TBI). Survival curves of C57BL/6 (A, C-D) or BALB/c (E) mice. (A, E) 4 doses of 10-NO_2_OA or vehicle (24 and 1 h before irradiation, 4 and 48 h after irradiation), (C) 2 doses of 10-NO_2_OA or vehicle (24 and 1 h before irradiation), (D) 2 doses of 10-NO_2_OA or vehicle (4 and 48 h after irradiation). (B) Body weight of C57BL/6 mice treated with 4 doses of 10-NO_2_OA or vehicle (24 and 1 h before irradiation, 4 and 48 h after irradiation). Data represent means ± SD, n = 34-35 (A, B), n = 9-10 (C), n = 7-9 (D), n = 8-11. Statistical analysis with Log-rank (Mantel-Cox) test (A, C-E). Body weight (B.W.); n.s. – not significant. Body weight (% of T0)

To examine the influence of treatment timing, 10-NO_2_OA was administered either before-only or after-only irradiation (Fig. 1C-D). Pre-treatment-only regimen resulted in 50% survival compared with 33% of vehicle-treated animals (not significant) and extended median survival from 14 to 27 days with stable body weight in both groups (Fig. 1C, Fig. S1A). In contrast, post-treatment-only regimen did not improve survival and did not affect mice body weight (67% in 10-NO_2_OA *vs*. 71% in vehicle; Fig. 1D, Fig.S2A).

To evaluate strain dependency, the experiment was repeated in more radiosensitive BALB/c mice. Under identical administration conditions with both pre- and post-treatment regimen, all animals succumbed around day 16 following TBI, and 10-NO_2_OA treatment had no detectable effect on survival of BALB/c mice (Fig. 1E).

### 10-NO₂OA mitigates radiation-induced neutropenia and thrombocytopenia

To assess the impact of 10-NO₂OA treatment on radiation-induced alterations in peripheral blood parameters, hematological profiles were analyzed 30 days after TBI (7.5 Gy) in surviving mice from control, TBI + vehicle, and TBI + 10-NO₂OA groups, with treatment administered 24 and 1 h prior to TBI and 4 and 48 h post-irradiation (Fig. 2A-D). Both TBI groups (TBI + vehicle and TBI + 10-NO_2_OA) exhibited significantly lower white blood cell (WBC) (Fig. 2A) and lymphocyte counts (Lym) (Fig. 2B) than controls, with Lym significantly reduced also when comparing TBI + 10-NO₂OA versus TBI + vehicle. Neutrophil (Neu) counts were significantly reduced only in TBI + vehicle group, remaining higher in control and TBI + 10-NO₂OA groups (Fig. 2C). Platelet (PLT) levels were decreased in both TBI groups compare with control, with a more pronounced reduction observed in TBI, which exhibited significantly lower counts than both control and TBI + 10-NO₂OA groups (Fig. 2D). Furthermore, the hematological profile was also analyzed in pre-treatment or post-treatment groups (Fig. S1B-E, Fig. S2B-E). Consistent with the four-dose treatment regimen, neither pre-treatment-only nor post-treatment-only with 10-NO_2_OA mitigated the decreases in WBC and Lym counts (Fig. S1B-C, Fig S2B-C). However, only pre-treatment regimen was able to significantly increase the PLT levels in TBI + 10-NO₂OA group compared with the TBI + vehicle group (Fig. S1E, Fig. S2E). Additionally, surviving mice in both treatment groups did not exhibit a decrease in neutrophils count following irradiation (Fig. S1D, Fig. S2D).

**Fig. 2:**
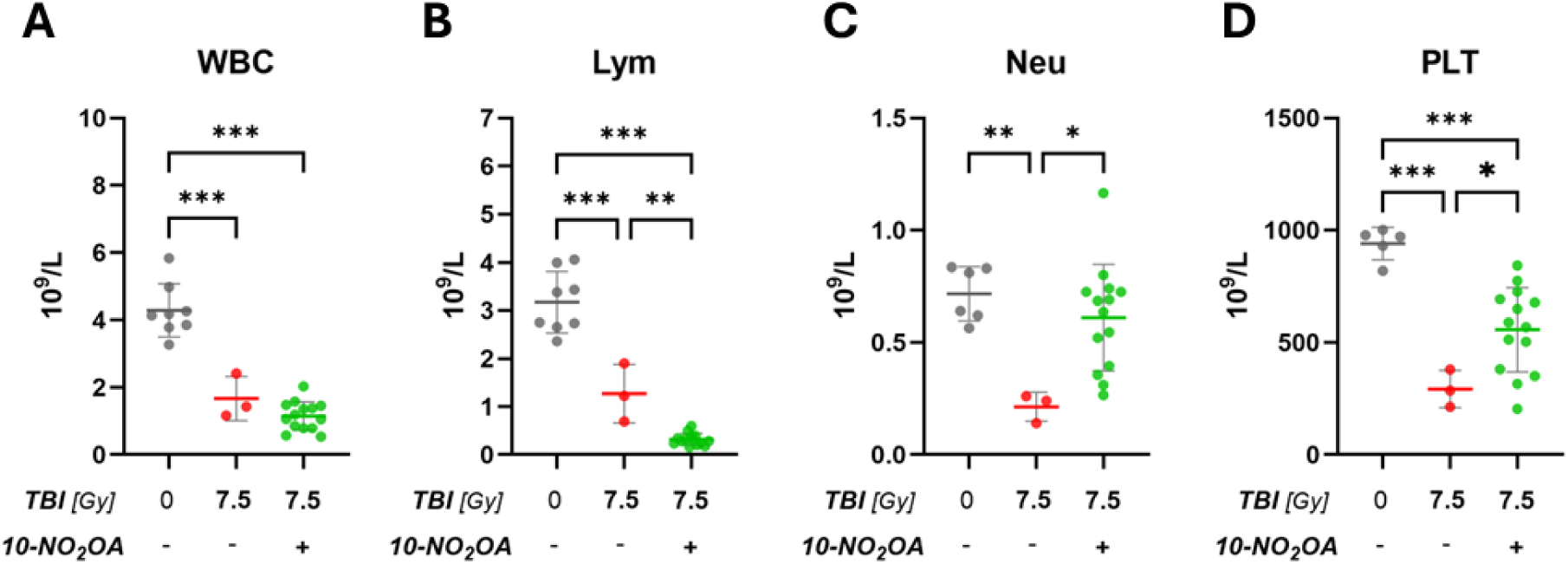
10-NO_2_OA treatment preserves neutrophil and platelet cell-counts after TBI. The level of (A) WBC, (B) Lym, (C) Neu and (D) PLT were analyzed from peripheral blood 30 days after TBI (7.5Gy) in surviving mice. C57BL/6 mice were treated with 4 doses of 10-NO_2_OA or vehicle (24 and 1 h before irradiation, 4 and 48 h after irradiation). Data represent means ± SD, n = 3-14. Statistical analysis with one-way ANOVA with Tukey’s multiple comparisons test; (*p < 0.05, **p < 0.01, ***p < 0.001). Total body irradiation (TBI); white blood cells (WBC); lymphocytes (Lym); neutrophils (Neu); platelets (PLT).

Together, these data show that four-dose 10-NO₂OA treatment not only significantly improves survival after TBI but also mitigates radiation-induced reductions of Neu and PLT in peripheral blood, highlighting its protective effects against radiation-induced hematological alterations.

### 10-NO₂OA transiently increases BMC but does not prevent sustained peripheral blood cytopenia

To further clarify the effect of 10-NO_2_OA on the hematopoietic component of ARS, mice were subjected to 4 Gy TBI, a sublethal dose selected to induce bone marrow injury while minimizing the contribution of radiation damage to other organs and administered 10-NO_2_OA or vehicle 24 and 1 h before irradiation, and 4 and 48 h thereafter (Fig. 3). Under these conditions, both irradiated groups (TBI + vehicle and TBI + 10-NO_2_OA) exhibited a significant reduction in GM-CFU (Fig. 3A), red blood cell (RBC; Fig. 3C), and WBC (Fig. 3F) counts at days 7 and 14 compared with controls. Notably, the decrease in RBC and WBC counts persisted through days 21 and 28 (Fig. 3C, F). In contrast, 10-NO_2_OA treatment led to a significant increase in BMCs on day 14 in comparison to both control and TBI + vehicle groups (Fig. 3B). However, consistent with the survival analysis, WBC counts on day 28 were lower in the TBI + 10-NO_2_OA group compared with the TBI + vehicle group (Fig. 3F). PLT counts and body weight (BW) were lower in both TBI groups compared to control group but remained comparable across all experimental groups throughout the study period (Fig. 3D, E). These findings indicate that while 10-NO_2_OA enhances bone marrow cellularity at day 14 following irradiation, it does not prevent the sustained peripheral blood cytopenia associated with H-ARS under the tested conditions.

**Fig. 3:**
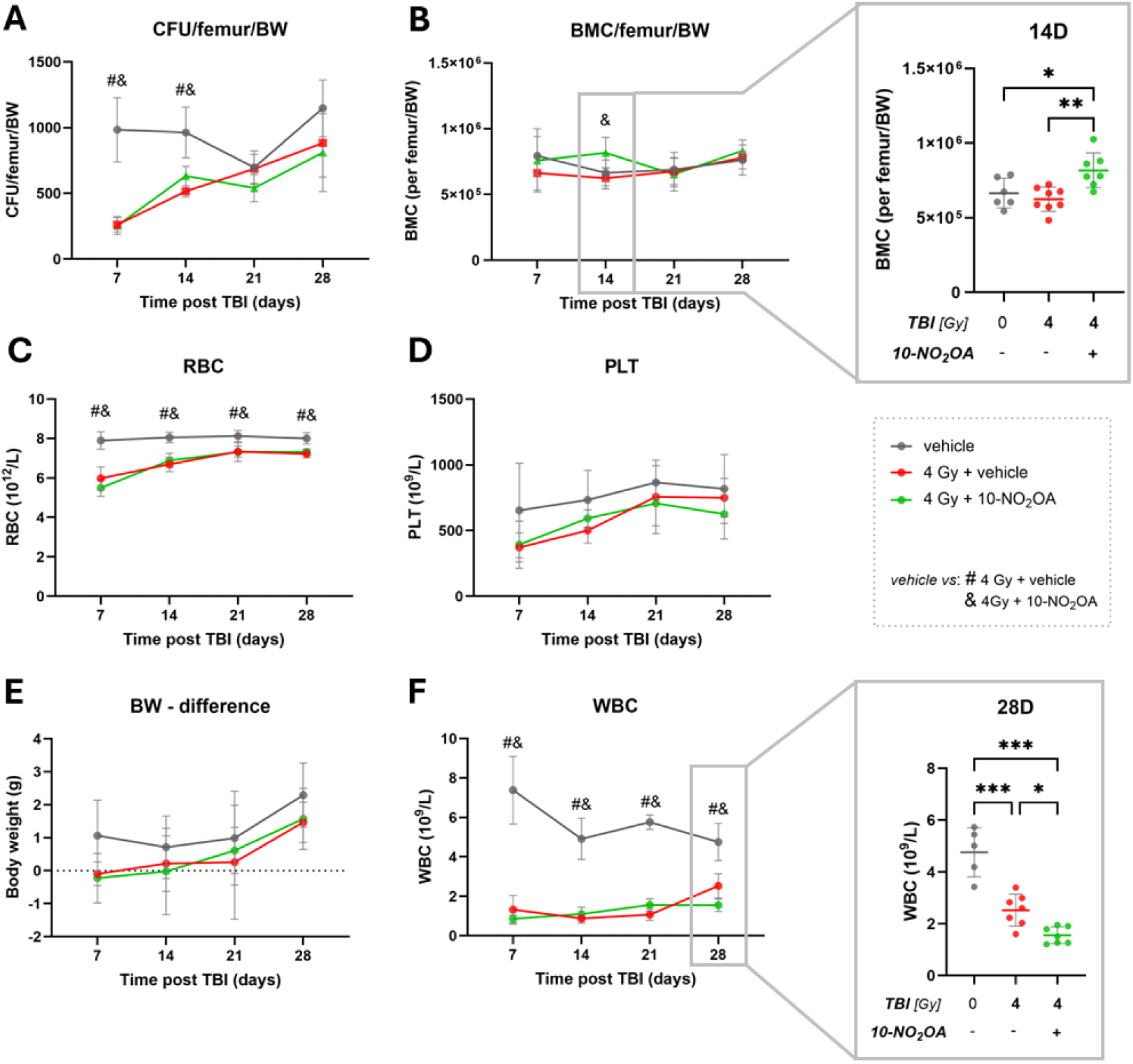
10-NO₂OA transiently increases BMCs. The number of (A) GM-CFU, (B) BMC, (C) RBC, (D) PLT, (F) WBC, and (E) body weight (BW) was analyzed in C57BL/6 mice 7, 14, 21, and 28 days after TBI (4 Gy) treated with 4 doses of 10-NO_2_OA or vehicle (24 and 1 h before irradiation, 4 and 48 h after irradiation). Data represent means ± SD, from n = 4-9, analyzed with one-way ANOVA followed by Tukey’s multiple comparisons test; ^#,&^ p < 0.05; *p < 0.05, **p < 0.01, ***p < 0.001. Symbol # means - vehicle *vs.* 4Gy+10-NO_2_OA; symbol & means - vehicle *vs.* 4Gy+vehicle. Total body irradiation (TBI); colony forming units (CFU); bone marrow cells (BMC); red blood cells (RBC); platelets (PLT); body weight (BW); white blood cells (WBC).

### 10-NO₂OA rapidly increases neutrophils, platelets, and G-CSF following TBI

To gain insight into the early hematological changes associated with the improved survival observed after 10-NO_2_OA treatment, we examined the short-term effects of 10-NO_2_OA pre-treatment following TBI (4 or 7.5 Gy) (Fig. 4). Consistent with our previous observations, TBI at both 4 and 7.5 Gy resulted in reduced WBC and lymphocyte counts 1 h post-irradiation, irrespective of 10-NO_2_OA treatment (Fig. 4A, B). In contrast, 10-NO₂OA treatment significantly increased neutrophil counts at both radiation doses relative to the corresponding TBI + vehicle groups (Fig. 4C), reaching levels that exceeded those in non-irradiated controls. Similarly, PLT counts following 7.5 Gy TBI were significantly increased by 10-NO₂OA compared with both control and TBI + vehicle (Fig. 4D) and significantly decreased in TBI + vehicle compared with control mice. Cytokine profiling (Fig. 4E) further revealed a broader early hematopoietic and inflammatory response following 10-NO₂OA treatment, characterized by increased plasma levels of BLC, IL-16, KC, MCP-1, and TIMP-1 compared with both control and TBI (7.5 Gy) + vehicle groups, whereas C5/C5a levels were reduced. Notably, G-SCF, KC and MCP-1, key mediators of neutrophil and monocyte recruitment, respectively, were markedly elevated (Fig. 4E). Further, to validate cytokine profiling data, plasma G-CSF levels were quantified by ELISA (Fig. 4F). Under these conditions 10-NO₂OA treatment significantly increased plasma levels of G-CSF compared with both control and TBI + vehicle (Fig. 4F). These early, selective effects are consistent with the pattern observed at later time points (presented at Fig. 2), where 10-NO_2_OA similarly preserves neutrophil and platelet levels without preventing the overall radiation-induced cytopenia, suggesting a lineage-specific hematological response that may contribute to the improved survival observed after irradiation.

**Fig. 4:**
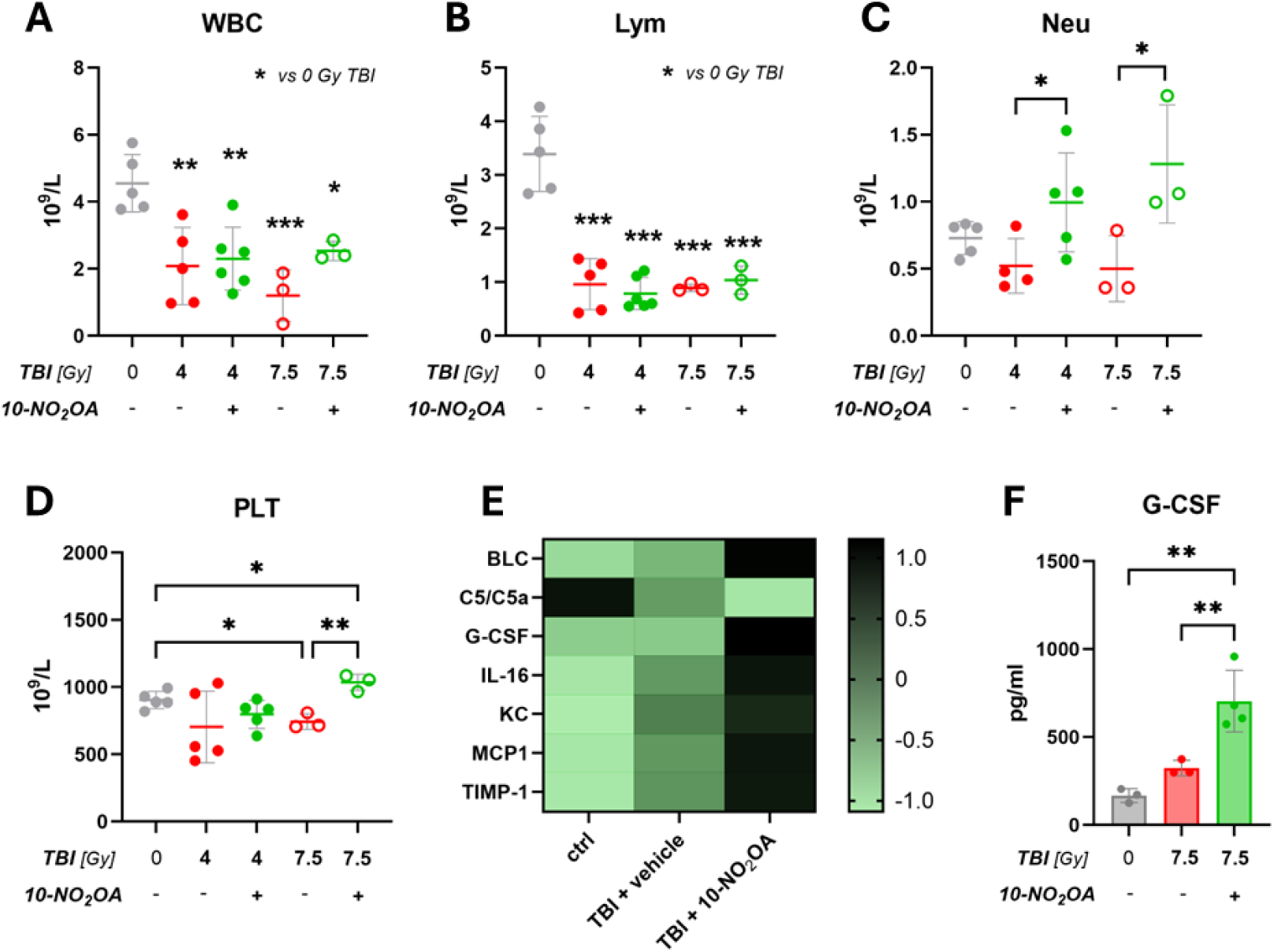
10-NO₂OA promotes an early myeloid response after TBI. The number of (A) WBC; (B) Lym; (C) Neu, and (D) PLT, cytokine assay (E) and mouse G-CSF level (F) were analyzed 1 h after TBI (4 or 7.5 Gy) in (A-D) peripheral blood or in (E-F) plasma from C57BL/6 mice treated with 2 doses of 10-NO_2_OA or vehicle (24 and 1 h before irradiation). Data (A–D, F) represent means ± SD, n = 3-6, analyzed with one-way ANOVA followed by Tukey’s multiple comparisons test; (*p < 0.05, **p < 0.01, ***p < 0.001). (E) Data from cytokine array are shown as means of the log-transformed z-scores from 3–4 pooled plasma samples per group, analyzed in duplicates. Total body irradiation (TBI); white blood cells (WBC); lymphocytes (Lym); neutrophils (Neu); platelets (PLT); B-lymphocyte chemoattractant (BLC); granulocyte colony-stimulating factor (G-CSF); interleukin 16 (IL-16); keratinocyte-derived chemokine (KC); monocyte Chemoattractant Protein-1 (MCP-1); tissue inhibitor of matrix metalloproteinase 1 (TIMP-1).

### 10-NO₂OA increases neutrophil counts in non-irradiated mice

To uncover whether 10-NO_2_OA directly modulates hematological parameters in the absence of irradiation, mice were treated with 10-NO_2_OA (24 and 1 h prior to sham irradiation), and peripheral blood cell counts were analyzed 3 days later (Fig. 5). Under these conditions, 10-NO_2_OA treatment significantly increased neutrophil counts (Fig. 5C), accompanied by a corresponding elevation in total WBC levels (Fig. 5A), while lymphocyte counts remained unchanged (Fig. 5B). Platelet counts showed a non-significant decreasing trend (Fig. 5D). Together, these findings indicate that 10-NO_2_OA can influence hematological parameters, particularly by increasing neutrophil counts independently of irradiation. This observation suggests that the elevated neutrophil levels observed after TBI may reflect, at least in part, a treatment-driven effect rather than solely protection from radiation-induced damage.

**Fig. 5:**
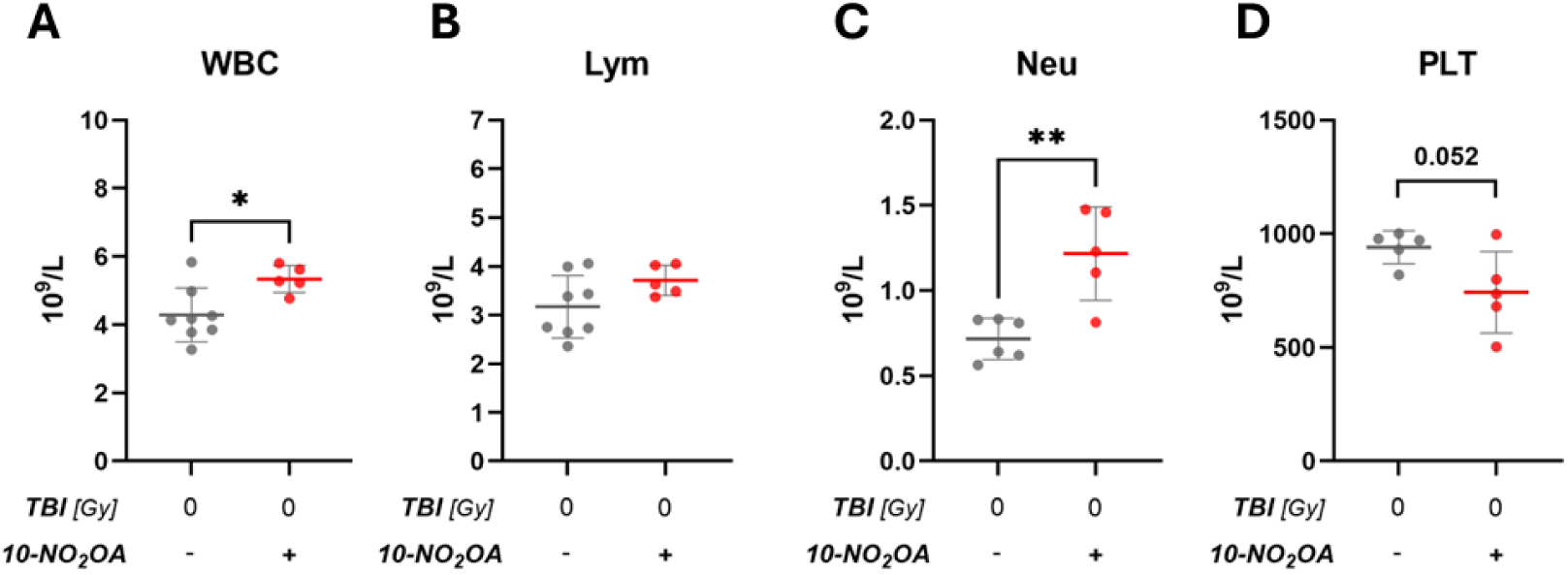
10-NO_2_OA increased peripheral neutrophil count in mice. The number of (A) WBC; (B) Lym; (C) Neu, and (D) PLT were analyzed in peripheral blood from C57BL/6 mice treated with 2 doses of 10-NO_2_OA or vehicle (24 and 1 h before sham irradiation). Data represent means ± SD, from 3 independent experiments, n = 5-8, and analyzed with unpaired Student’s t-test; (*p < 0.05, **p < 0.01). Total body irradiation (TBI); white blood cells (WBC); lymphocytes (Lym); neutrophils (Neu); platelets (PLT).

### 10-NO₂OA activates NRF2-responsive signaling in irradiated BMCs

To determine whether the radioprotective effects of 10-NO₂OA are associated with activation of the NRF2 pathway under irradiation conditions, primary BMCs isolated from C57BL/6 mice were treated *ex vivo* with 10-NO₂OA or vehicle and analyzed 3h following gamma irradiation. The chemical structures of the nitro-fatty acids included in the comparative analysis are shown in Fig. 6E. Consistent with our previous findings in non-irradiated BMCs cultured under physiological oxygen conditions [8], 10-NO₂OA elicited the strongest NRF2-associated response among all nitro-fatty acids tested (Fig. 6B-D). Specifically, 10-NO₂OA significantly increased the expression of the NRF2 target genes *Nqo1*, *Hmox1*, and *Gclm* in irradiated BMCs (Fig. 6B-D). Notably, neither 10-NO₂OA nor any of the other nitro-fatty acids tested significantly altered the expression of *Nfe2l2* itself (Fig. 6A). Furthermore, exploratory analysis of γH2AX, a marker of DNA double-strand break signaling, performed in two independent experiments, suggested that 10-NO₂OA was associated with slightly lower γH2AX levels than the irradiated control (Fig. S3).

**Fig. 6:**
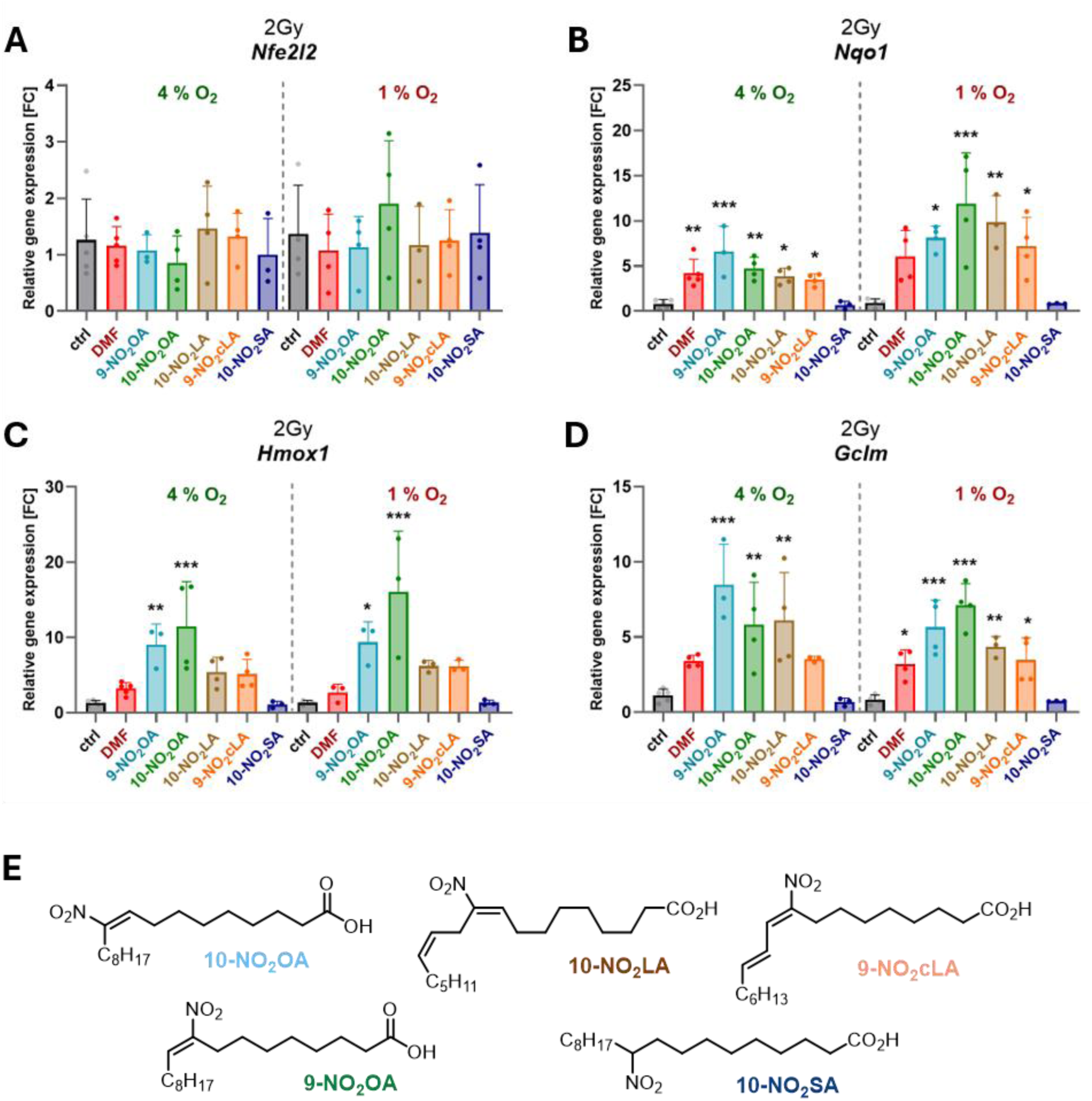
10-NO_2_OA activates NRF2-responsive signaling in irradiated BMCs. The relative expression of (A) *Nfe2l2*, (B) *Nqo1*, (C) *Hmox1*, and (D) *Gclm* was analyzed in physiological oxygen conditions (4 and 1% O_2_). Primary BMCs isolated from C57BL/6 mice were treated with the indicated nitro-fatty acids (2 µM) or vehicle for 1 h before gamma irradiation (2 Gy), then incubated for an additional 3 h, and subsequently lysed for expression analysis. Data represent means ± SD, from at least 3 independent experiments (n = 3-5), and analyzed with one-way ANOVA with Dunnett multiple comparisons test; (*p < 0.05, **p < 0.01, ***p < 0.001). (E) Chemical structures of the nitro-fatty acids included in the comparative analysis. Nuclear factor erythroid 2-related factor 2 (Nfe2l2); NAD(P)H quinone oxidoreductase 1 (Nqo1); Heme Oxygenase 1 (Hmox1); glutamate-cysteine ligase modifier subunit (Gclm); control (ctrl); dimethyl fumarate (DMF) as positive control; 9- or 10-nitro-oleic acid (9-/10-NO_2_OA); 10-nitro linoleic acid (10-NO_2_LA); 9-nitro-conjugated linoleic acid (9-NO_2_cLA); 10-nitro-stearic acid (10-NO_2_SA). (E) Structures of nitro-fatty acids tested in the present study.

To explore these findings *in vivo*, we assessed the expression of the NRF2-responsive genes in BMCs isolated from C57BL/6 mice pre-treated with 10-NO_2_OA or vehicle 24 and 1 h before TBI (7.5 Gy). In two independent experiments, 10-NO_2_OA enhanced the expression of both *Hmox1* and *Gclm* following TBI compared to TBI alone (Fig. S4). Together, these results indicate that 10-NO₂OA activates NRF2-dependent signaling following irradiation both *ex vivo* and *in vivo*. These findings are consistent with a potential contribution of NRF2 pathway activation in the radioprotective effects of 10-NO₂OA.

### 10-NO₂OA attenuates radiation-induced intestinal alterations after TBI

While 10-NO₂OA produced several beneficial hematopoietic effects following TBI, these findings may not fully explain the observed survival advantage. Therefore, we performed an exploratory assessment of gastrointestinal responses to determine whether intestinal protection might also contribute to the improved survival of 10-NO₂OA-treated mice (Fig. 7). Mice were exposed to 7.5 Gy TBI and treated with 10-NO_2_-OA or vehicle 24 and 1 h before irradiation and 4 and 48 h thereafter. At day 10 post-TBI, 10-NO_2_-OA–treated mice exhibited a better-preserved intestinal appearance than vehicle-treated mice, which displayed overt macroscopic signs of intestinal injury. Specifically, the large intestine appeared reduced in size and hemorrhagic in most TBI animals (Fig. 7A; upper representative images). Further, abdominal inspection revealed dilated intestinal segments with pale to pinkish coloration (Fig. 7A; bottom representative images). The intestinal wall appeared thin and translucent in some areas, and luminal contents were mostly unevenly distributed along the intestine. No obvious perforations were observed. These macroscopic alterations were not evident in 10-NO_2_-OA–treated mice (Fig. 7A; right images). On day 6 post-TBI, cytokine array from conditioned media of *ex vivo* explanted large intestine was analyzed (Fig. 7B). Irradiation reduced the levels of multiple signaling factors, including G-CSF, GM-CSF, and M-CSF, as well as IL-6, IP-10, CXCL1, MIP-2, and TIMP-1. Treatment with 10-NO_2_-OA reversed the irradiation-induced changes in cytokine secretion and increased the levels of G-CSF, GM-CSF, M-CSF, IL-6, IP-10, and TIMP-1. Additionally, CXCL1 and MIP-2 showed a clear shift towards control levels. Furthermore, on day 6 post-TBI, protein levels of the intestinal crypt marker SOX9, the apoptosis-associated protein cleaved caspase 3, and the cell cycle marker cyclin D1 were analyzed in *ex vivo* explanted large intestine tissue (Fig. 7C). In contrast to cleaved caspase-3, SOX9 and cyclin D1 protein levels were slightly elevated in the TBI + NO_2_OA group compared with TBI + vehicle group (Fig. 7C). Although these differences did not reach statistical significance, they may suggest improved tissue viability and regenerative capacity in 10-NO_2_OA treated mice.

**Fig. 7:**
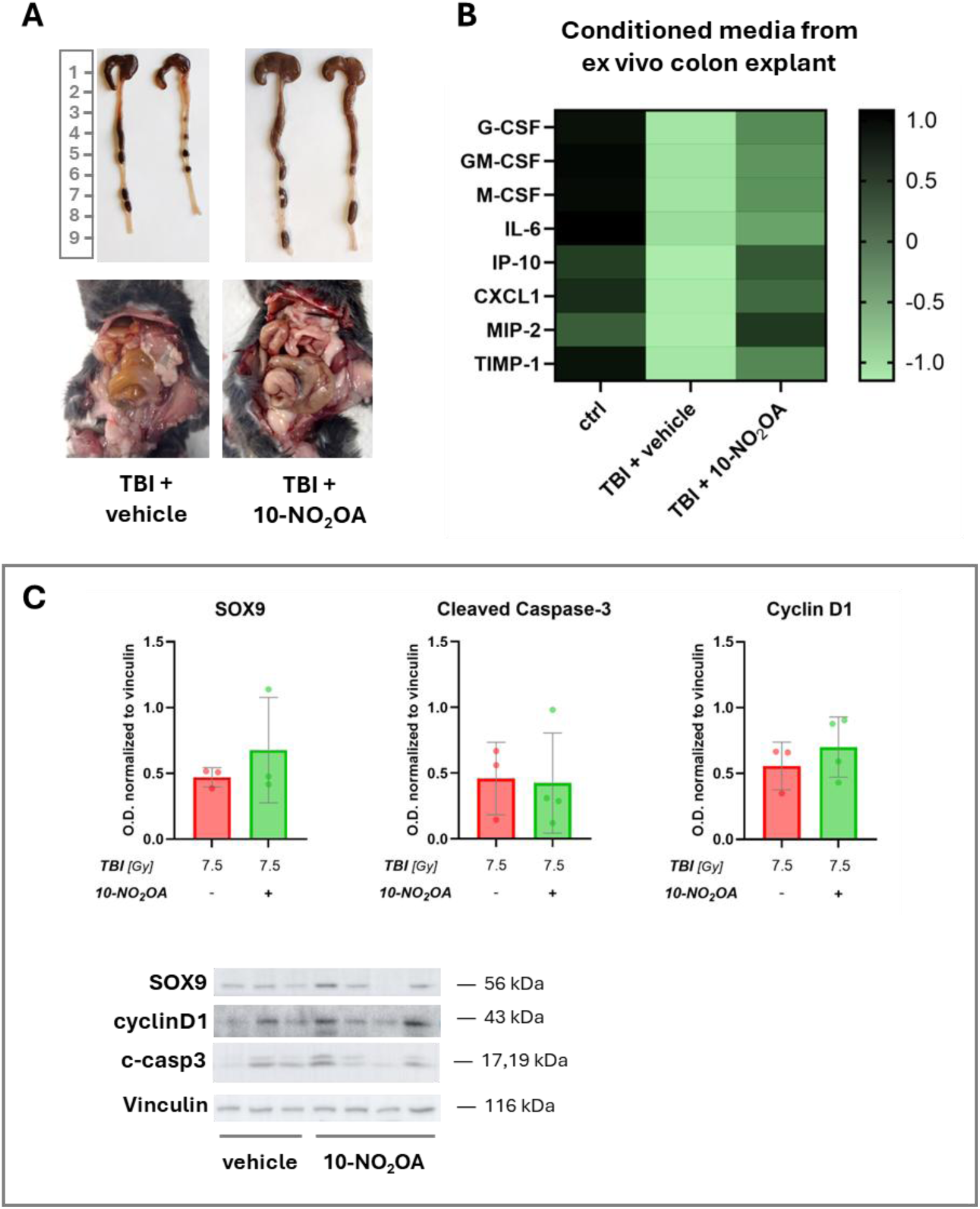
10-NO_2_OA mitigates gastrointestinal acute radiation syndrome. (A) Representative images of (upper) explanted large intestine and (bottom) intestinal tract of C57BL/6 mice (10 days post-TBI) treated with 4 doses of 10-NO_2_OA or vehicle (24 and 1 h before irradiation, 4 and 48 h after irradiation). (B) Cytokine array of colon explants from different treatment groups 6 days post-TBI: Conditioned media from colon tissue explants of 3–4 animals per group were pooled for analysis. Results are presented as means of the log-transformed z-scores from pooled conditioned media per group and analyzed in technical duplicates of the array. Total body irradiation (TBI), G-CSF: granulocyte colony-stimulating factor, GM-CSF: granulocyte-macrophage colony-stimulating factor, M-CSF: macrophage colony-stimulating factor, IL-6: interleukin-6, IP-10: interferon gamma-induced protein 10 (CXCL10), CXCL1: C-X-C motif chemokine ligand 1, MIP-2: macrophage inflammatory protein-2 (CXCL2), TIMP-1: tissue inhibitor of metalloproteinases-1. (C) Western blot analysis of SOX9 (SRY-box transcription factor 9), cleaved caspase-3 (c-casp3), and cyclin D1 protein levels in intestinal tissue explants obtained from C57BL/6 mice 6 days post-TBI treated with 4 doses of 10-NO₂OA or vehicle (24 and 1 h before irradiation, 4 and 48 h after irradiation). Representative blots and corresponding densitometric quantification are shown. Data represent means ± SD, from n = 3–4 independent experiments.

Taken together with the survival data, these results are consistent with a systemic effect of 10-NO₂OA that is not limited to the hematopoietic compartment.

## Discussion

In this study, we show that 10-NO₂OA improves survival after LD_50_-level gamma irradiation in C57BL/6 mice when administered in a combined pre- and post-irradiation regimen. This survival benefit was accompanied by early changes in hematological parameters, including increased neutrophil and platelet counts and elevated plasma G-CSF, while sustained recovery of peripheral blood cellularity remained incomplete. These findings suggest that 10-NO₂OA does not simply prevent global hematopoietic injury but instead promotes a selective and time-dependent response that may support survival during the acute phase after irradiation. In parallel, 10-NO₂OA activated NRF2-responsive signaling in irradiated BMCs and was associated with modulation of intestinal responses. Together, these data indicate that the radioprotective effects of 10-NO₂OA are likely multifactorial, involving hematological, cytoprotective, and intestinal components discussed below.

An important finding of the present study was the strong dependence of 10-NO₂OA efficacy on the treatment schedule. This effect was observed only with the combined pre- and post-irradiation treatment regimen, whereas pre-irradiation treatment alone produced only a modest, non-significant benefit and post-irradiation treatment alone was ineffective. These findings indicate that the radioprotective efficacy of 10-NO_2_OA is strongly dependent on treatment timing. Such schedule dependence is well recognized among radiation countermeasures. For example, the clinically approved radioprotector amifostine is administered before irradiation, whereas hematopoietic growth factors such as G-CSF are used after radiation exposure as radiomitigators [13]. More broadly, optimization of treatment schedules remains an important aspect of radiation-countermeasure development [14, 15]. The present findings suggest that pre-irradiation exposure to 10-NO₂OA is an important determinant of efficacy. However, although the combined pre- and post-irradiation regimen produced the greatest survival benefit, the current study does not allow discrimination between the relative contributions of continued post-irradiation dosing and prolonged overall exposure to the compound. Notably, pretreatment with several NRF2-activating compounds failed to improve survival following irradiation, suggesting that NRF2 activation before radiation exposure alone may be insufficient to confer meaningful radioprotection [16]. In addition, the absence of a survival benefit in BALB/c mice suggests that host genetic background may represent another important determinant of responsiveness to 10-NO₂OA treatment. BALB/c mice are known to be more radiosensitive than C57BL/6 mice, in part due to differences in DNA damage responses and repair capacity [17], and it is therefore possible that the magnitude of protection afforded by 10-NO₂OA was insufficient to overcome the greater radiation sensitivity of this strain.

Although reports on nitro-fatty acids as radiation countermeasures are currently limited, findings from other fatty acids further support the radiomitigative potential. Notably, omega-3 fatty acids attenuated radiation-induced myelosuppression and promoted recovery of white blood cells and platelets in irradiated mice [18], supporting the concept that lipid mediators can beneficially modulate hematopoietic responses to radiation exposure. In line with these observations, we previously demonstrated that the mixed 9-/10-NO_2_OA isomer accelerated hematopoietic recovery after sublethal total body irradiation [11], indicating its capacity to promote hematopoietic regeneration, with the most pronounced effects occurring during the early phase of recovery. Here, we provide evidence that regio- and stereo-chemically defined 10-NO₂OA enhanced hematopoietic recovery but did not fully restore peripheral blood cell counts to control levels at later time points. The transient nature of the hematopoietic response may reflect the lack of repeated administration of 10-NO_2_OA, as studies employing prolonged dosing regimens, such as captopril treatment, have demonstrated sustained mitigation of radiation-induced hematopoietic injury [14]. Improved colony-forming capacity has frequently been associated with radioprotection in preclinical models [19]. In line with this concept, 10-NO_2_OA increased GM-CFU numbers and bone marrow cellularity for up to 14 days after sublethal TBI, indicating a positive effect on hematopoietic progenitor function and bone marrow regeneration. While these improvements were not accompanied by corresponding increases in total white blood cell, erythrocyte, and platelet counts, consistent with our previous findings [11], the results suggest that 10-NO₂OA preferentially supports specific hematopoietic compartments during the recovery process.

Interestingly, LD_50_-level TBI survivors treated with 10-NO_2_OA exhibited higher neutrophil and platelet counts, whereas lymphocyte counts remained lower. Consistent with these long-term *in vivo* findings, mice treated with 10-NO_2_OA displayed increased numbers of platelets and, in particular, neutrophils shortly after TBI, accompanied by elevated G-CSF levels compared with the TBI-only group. The observed increase in G-CSF may be especially important in H-ARS, where neutropenia and thrombocytopenia are major contributors to morbidity and mortality. Stimulation of granulopoiesis by G-CSF represents an established approach for mitigating radiation-induced hematopoietic injury [1, 4]. Although 10-NO_2_OA did not fully prevent radiation-induced cytopenia, the rapid induction of G-CSF together with increased neutrophil counts may have contributed to the observed survival benefit, consistent with previous studies demonstrating an association between endogenous G-CSF induction, hematopoietic recovery, and improved survival after irradiation [19]. Rather than broadly restoring all hematological parameters, 10-NO_2_OA consistently increased G-CSF levels, neutrophil counts, and platelet counts, suggesting preferential modulation of myeloid and thrombocytic responses following irradiation. This is of importance as granulocytopenia and thrombocytopenia may contribute to reduced host defense, sepsis and hemorrhage in exposed individuals [20]. Besides recombinant growth factors (e.g. filgrastim), romiplostim, which enhances platelet production via stimulation of bone marrow megakaryocytes, was approved for management of H-ARS [21]. Collectively, these findings suggest that 10-NO₂OA promotes a selective hematopoietic response characterized by increased G-CSF production and preservation of neutrophil and platelet populations. Such targeted modulation may be more important for survival after irradiation than complete normalization of hematological parameters.

Notably, the effects on neutrophils and platelets were more pronounced at the higher radiation dose. Based on previous reports demonstrating that NRF2 signaling regulates hematopoietic stem and progenitor cell function and promotes differentiation toward granulocyte-monocyte lineages while limiting erythroid and lymphoid commitment [22, 23], we aimed to determine whether NRF2 signaling mediates these effects.

NRF2 has emerged as an important regulator of the radiation response, coordinating antioxidant and cytoprotective defenses, DNA damage responses, and inflammatory signaling pathways that collectively mitigate radiation-induced tissue injury and promote recovery (reviewed in [24, 25]). Importantly, NRF2 activity must be tightly regulated in hematopoiesis: transient activation can facilitate hematopoietic stem cell cycle entry and differentiation during regenerative stress, whereas sustained activation progressively erodes quiescence and self-renewal, culminating in hematopoietic stem cell exhaustion and impaired bone marrow reconstitution [26, 27]. Previous studies have shown that NRF2 promotes hematopoietic recovery following irradiation, whereas NRF2 deficiency increases susceptibility to radiation-induced injury [22, 28]. Consistent with this concept, several nitro-fatty acids have been reported to activate NRF2 signaling [29, 30], including in primary bone marrow cells [8]. In the present study, we showed that 10-NO₂OA activates NRF2 signaling in gamma-irradiated bone marrow cells and elicits the strongest NRF2-associated response among the nitro-fatty acids tested, as evidenced by increased expression of the canonical NRF2 target genes *Nqo1*, *Hmox1*, and *Gclm*. Together, these findings are compatible with a cytoprotective contribution of NRF2 signaling to the radioprotective effects of 10-NO_2_OA. However, the causal requirement of NRF2 for the observed radioprotection was not addressed in the present study. Notably, the substantial survival benefit observed following irradiation was accompanied by only modest effects on long-term hematopoietic recovery, suggesting that mechanisms beyond preservation of bone marrow cellularity and peripheral blood counts may contribute to the observed radioprotection. This possibility prompted us to explore whether modulation of intestinal responses might represent an additional component of the protective effects of 10-NO_2_OA.

Because gastrointestinal ARS typically develops at radiation doses above 6 Gy and becomes the dominant life-threatening syndrome at ≥10 Gy, preservation of intestinal integrity represents a critical determinant of survival following high-dose TBI (reviewed in [31]). Given the beneficial effects of 10-NO_2_OA on mouse survival, we performed exploratory investigations of the gastrointestinal tract. Although colon length was not significantly affected by 10-NO_2_OA treatment, cytokine profiling of colon explants revealed partial restoration of irradiation-suppressed cytokines and growth factors. The absence of detectable differences in colon length may reflect the lower radiation dose, differences in tissue collection timing compared with dedicated gastrointestinal injury studies [32], and the use of tissues obtained from LD_50_ survival experiments rather than specialized gastrointestinal-ARS models. Notably, restoration of the cytokine profile in colon explants collected 6 days after irradiation may indicate modulation of both inflammatory and regenerative processes, as this time point corresponds to a dynamic phase of intestinal injury and crypt regeneration in experimental gastrointestinal-ARS models [33]. Therefore, future studies should investigate whether oral administration of 10-NO_2_OA can provide additional protection against radiation-induced gastrointestinal injury.

Collectively, our data support a model in which 10-NO_2_OA exerts radioprotective activity through multiple complementary mechanisms. Early modulation of myeloid responses, activation of cytoprotective signaling pathways, and preservation of intestinal homeostasis may each contribute to improved survival. The absence of complete hematopoietic recovery despite substantial survival improvement argues against a mechanism exclusively dependent on bone marrow protection.

Several limitations of the present study should be acknowledged. First, although 10-NO₂OA significantly improved survival following LD₅₀-level irradiation, the relationship between survival and the observed hematopoietic, intestinal, and molecular changes remains associative rather than causal. In particular, although 10-NO₂OA activated NRF2-responsive signaling, the present study was not designed to determine whether NRF2 activation is required for its radioprotective effects. Nevertheless, the robust survival benefit, together with consistent biological responses across multiple endpoints, provides a strong foundation for future mechanistic studies. Second, the intestinal analyses were exploratory in nature and did not include histopathological evaluation, intestinal permeability measurements, or functional assessment of gastrointestinal injury. Despite these limitations, the findings broaden the current understanding of 10-NO_2_OA in radiation biology and provide an important starting point for investigating its effects on radiation-induced intestinal injury *in vivo*. Finally, the strain-dependent differences observed between C57BL/6 and BALB/c mice indicate that host-specific factors may influence the efficacy of 10-NO₂OA and warrant further investigation, particularly in the context of developing more personalized radioprotective strategies. Future studies should therefore focus on defining the relative contribution of hematopoietic, intestinal, and NRF2-dependent mechanisms to the survival benefit conferred by 10-NO₂OA.

**Graphical abstract.**
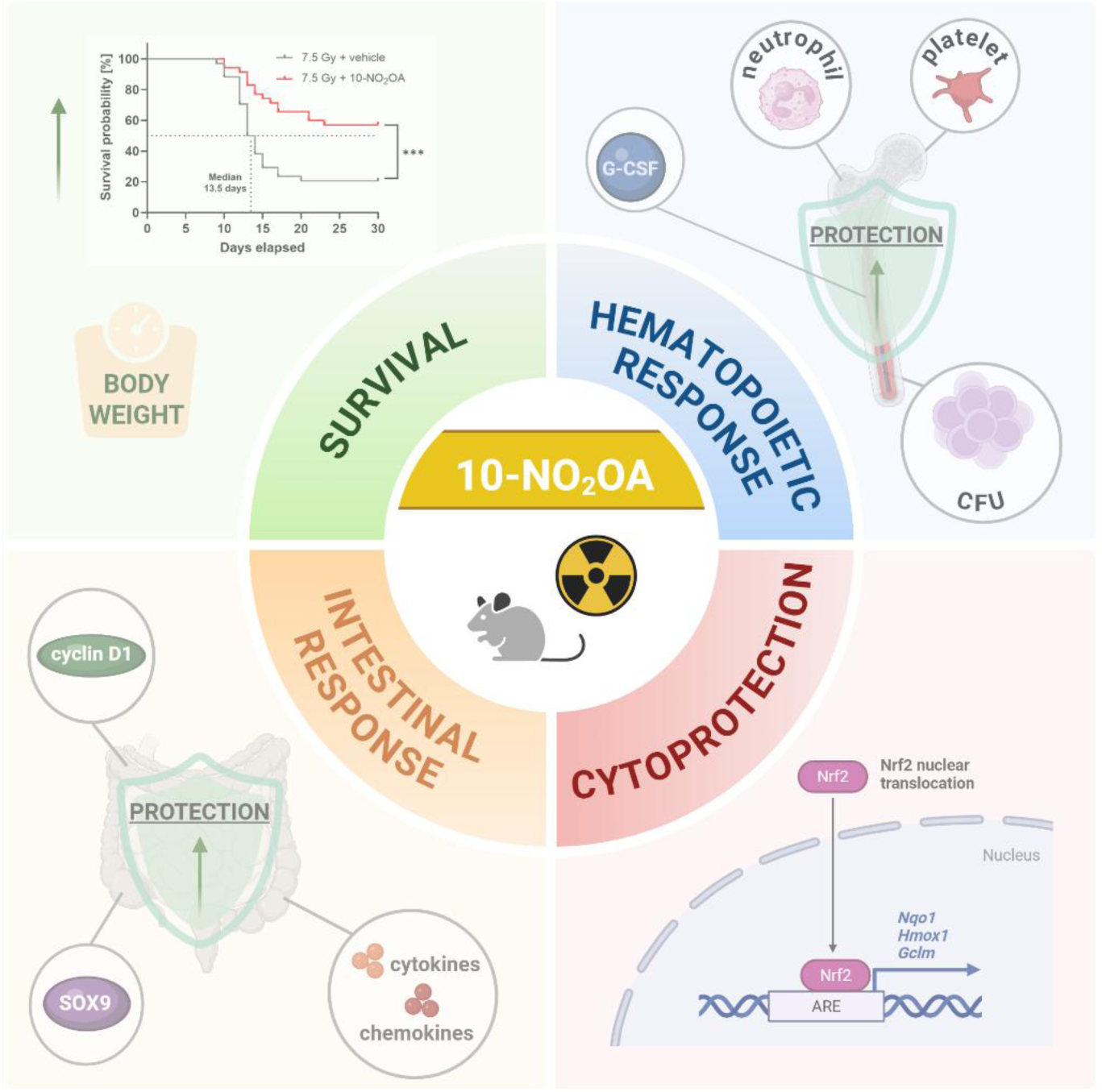
Created in BioRender. Pereckova, J. (2026) https://BioRender.com/wxgseko.

## Conclusions

In conclusion, 10-NO_2_OA improved survival following LD_50_-level gamma irradiation in C57BL/6 mice when administered in a combined pre- and post-irradiation regimen. The survival benefit was associated with selective hematopoietic effects, including increased neutrophil and platelet counts and elevated plasma G-CSF levels, rather than complete restoration of hematopoiesis. In addition, 10-NO_2_OA activated NRF2-responsive signaling and was associated with modulation of intestinal responses following irradiation. Together, these findings identify 10-NO₂OA as a promising radioprotective agent and suggest that its survival benefit is associated with multiple biological responses extending beyond the hematopoietic compartment.

## Supporting information

Supplementary file 2_Synthesis

Publication License Sep-25-2026- Scheme 1

Publication License Sep-25-2026-graphical abstract

## Declarations and Statements

### Authorship contribution statement

**Jana Pereckova:** Conceptualization; Investigation; Methodology; Data Analysis; Writing – original draft; Writing - review & editing. **Rachel Vichova:** Conceptualization; Investigation; Methodology; Writing - review & editing. **Jiri Toufar:** Investigation; Methodology; Writing - review & editing. **Daniel Chrenko**: Investigation; Methodology; Writing - review & editing. **Jiří Pospíšil**: Investigation; Methodology; Writing - review & editing. **Tomas Perecko:** Conceptualization; Data curation; Formal analysis; Investigation; Methodology; Writing - original draft; Writing - review & editing; Funding acquisition.

## Acknowledgments

We thank Zuzana Hoferova for technical assistance with mouse experiments. Scheme 1 (Pereckova, J. (2026) https://BioRender.com/hvl3xom) and graphical abstract (Pereckova, J. (2026) https://BioRender.com/wxgseko) were created with BioRender.com.

## Data Availability Statement

The data that support the biological findings of this study are openly available in Zenodo at https://doi.org/10.5281/zenodo.22938972. The data that support the findings of this study related to chemical synthesis part are openly available in Zenodo at https://doi.org/10.5281/zenodo.22940442.

## Use of Generative AI

A large language model (Microsoft Copilot, GPT-5 chat model) was used for language editing and stylistic refinement. The authors confirm the originality and accuracy of the content and take full responsibility for it.

## Supplementary Information

Supplementary file 1_SI Figures (PDF format)

Contains Figures S1–S4 showing surviving experiments with pre-irradiation and post-irradiation treatment regimens (S1, S2), and mechanistic experiments on DNA damage signaling (S3) and NRF-target genes expression (S4) *in vivo*.

Supplementary file 2_Synthesis (PDF format)

Contains detailed synthetic procedures, purification methods, and characterization data of tested nitro-fatty acids.

## References

1. Arnautou P, Garnier G, Maillot J, Konopacki J, Brachet M, Bonnin A, Amabile JC, Malfuson JV. Management of acute radiation syndrome. Transfus Clin Biol. 2024;31(4):253–9. 10.1016/j.tracli.2024.07.002.

2. Freeman ML. Gastrointestinal acute radiation syndrome: current knowledge and perspectives. Cell Death Discov. 2025;11(1):235. 10.1038/s41420-025-02525-6.

3. Anno GH, Baum SJ, Withers HR, Young RW. Symptomatology of acute radiation effects in humans after exposure to doses of 0.5-30 Gy. Health Phys. 1989;56(6):821–38. 10.1097/00004032-198906000-00001.

4. Lazarus HM, Gale RP. Cytokine therapy of acute radiation syndrome. Best Pract Res Clin Haematol. 2024;37(4):101599. 10.1016/j.beha.2025.101599.

5. Khoo NKH, Schopfer FJ. Nitrated fatty acids: from diet to disease. Curr Opin Physiol. 2019;9:67–72. 10.1016/j.cophys.2019.04.013.

6. Rom O, Khoo NKH, Chen YE, Villacorta L. Inflammatory signaling and metabolic regulation by nitro-fatty acids. Nitric Oxide. 2018. 10.1016/j.niox.2018.03.017.

7. Rubbo H. Nitro-fatty acids: novel anti-inflammatory lipid mediators. Braz J Med Biol Res. 2013;46(9):728–34. 10.1590/1414-431X20133202.

8. Chrenko D, Pereckova J, Zatloukalova M, Vacek J, Pospisil J, Perecko T. Regio- and Stereoselective Synthesis of Nitro-fatty Acids as NRF2 Pathway Activators Working under Ambient or Hypoxic Conditions. J Med Chem. 2025;68(11):12172–84. 10.1021/acs.jmedchem.5c00982.

9. Kansanen E, Kuosmanen SM, Ruotsalainen AK, Hynynen H, Levonen AL. Nitro-Oleic Acid Regulates Endothelin Signaling in Human Endothelial Cells. Mol Pharmacol. 2017;92(4):481–90. 10.1124/mol.117.109751.

10. Salvatore SR, Vitturi DA, Fazzari M, Jorkasky DK, Schopfer FJ. Evaluation of 10-Nitro Oleic Acid Bio-Elimination in Rats and Humans. Sci Rep. 2017;7:39900. 10.1038/srep39900.

11. Perecko T, Hoferova Z, Hofer M, Pereckova J, Falk M. Administration of nitro-oleic acid mitigates radiation-induced hematopoietic injury in mice. Life Sci. 2022;310:121106. 10.1016/j.lfs.2022.121106.

12. Bon DJYD, Chrenko D, Kováč O, Ferugová V, Lasák P, Fuksová M, Zálešák F, Pospíšil J. Julia-Kocienski-Like Connective C−C and C=C Bond-Forming Reaction. Advanced Synthesis & Catalysis. 2024;366(3):480–7. 10.1002/adsc.202301054.

13. Plett PA, Sampson CH, Chua HL, Joshi M, Booth C, Gough A, Johnson CS, Katz BP, Farese AM, Parker J, MacVittie TJ, Orschell CM. Establishing a murine model of the hematopoietic syndrome of the acute radiation syndrome. Health Phys. 2012;103(4):343–55. 10.1097/HP.0b013e3182667309.

14. McCart EA, Lee YH, Jha J, Mungunsukh O, Rittase WB, Summers TA, Jr., Muir J, Day RM. Delayed Captopril Administration Mitigates Hematopoietic Injury in a Murine Model of Total Body Irradiation. Sci Rep. 2019;9(1):2198. 10.1038/s41598-019-38651-2.

15. Singh VK, Serebrenik AA, Wise SY, Petrus SA, Fatanmi OO, Kaytor MD. BIO 300: A Prophylactic Radiation Countermeasure for Acute Radiation Syndrome. Mil Med. 2024;189(Suppl 3):390–8. 10.1093/milmed/usae156.

16. McDonald JT, Kim K, Norris AJ, Vlashi E, Phillips TM, Lagadec C, Della Donna L, Ratikan J, Szelag H, Hlatky L, McBride WH. Ionizing radiation activates the Nrf2 antioxidant response. Cancer Res. 2010;70(21):8886–95. 10.1158/0008-5472.CAN-10-0171.

17. Williams JP, Brown SL, Georges GE, Hauer-Jensen M, Hill RP, Huser AK, Kirsch DG, Macvittie TJ, Mason KA, Medhora MM, Moulder JE, Okunieff P, Otterson MF, Robbins ME, Smathers JB, McBride WH. Animal models for medical countermeasures to radiation exposure. Radiat Res. 2010;173(4):557–78. 10.1667/RR1880.1.

18. Sun M, Pang L, Ju X, Sun H, Yu J, Zhao H, Yao W, Wei M. Attenuating effects of omega-3 fatty acids (Omegaven) on irradiation-induced intestinal injury in mice. Food Chem Toxicol. 2014;64:275–80. 10.1016/j.fct.2013.11.051.

19. Yi L, Tian M, Piao C, Gao G, Wu L, Pan Y, Liu J. The protective effects of 1,2-propanediol against radiation-induced hematopoietic injury in mice. Biomed Pharmacother. 2019;114:108806. 10.1016/j.biopha.2019.108806.

20. Heslet L, Bay C, Nepper-Christensen S. Acute radiation syndrome (ARS) - treatment of the reduced host defense. Int J Gen Med. 2012;5:105–15. 10.2147/IJGM.S22177.

21. Singh VK, Seed TM. Radiation countermeasures for hematopoietic acute radiation syndrome: growth factors, cytokines and beyond. Int J Radiat Biol. 2021;97(11):1526–47. 10.1080/09553002.2021.1969054.

22. Merchant AA, Singh A, Matsui W, Biswal S. The redox-sensitive transcription factor Nrf2 regulates murine hematopoietic stem cell survival independently of ROS levels. Blood. 2011;118(25):6572–9. 10.1182/blood-2011-05-355362.

23. Murakami S, Shimizu R, Romeo PH, Yamamoto M, Motohashi H. Keap1-Nrf2 system regulates cell fate determination of hematopoietic stem cells. Genes Cells. 2014;19(3):239–53. 10.1111/gtc.12126.

24. Anuranjani, Bala M. Concerted action of Nrf2-ARE pathway, MRN complex, HMGB1 and inflammatory cytokines - implication in modification of radiation damage. Redox Biol. 2014;2:832-46. 10.1016/j.redox.2014.02.008.

25. Moloudi K, Sangsuwan T, Monzen S, Fujishima Y, Anderson D, Frey B, Miura T, Azariasl S, Yasuda H, Haghdoost S. Nuclear Factor Erythroid 2-Related Factor 2 (NRF2) as a Biomarker for Radiation Dosimetry and Health Risk Assessment: A Review. Antioxidants (Basel). 2025;14(12). 10.3390/antiox14121393.

26. Murakami S, Suzuki T, Harigae H, Romeo PH, Yamamoto M, Motohashi H. NRF2 Activation Impairs Quiescence and Bone Marrow Reconstitution Capacity of Hematopoietic Stem Cells. Mol Cell Biol. 2017;37(19). 10.1128/MCB.00086-17.

27. Tsai JJ, Dudakov JA, Takahashi K, Shieh JH, Velardi E, Holland AM, Singer NV, West ML, Smith OM, Young LF, Shono Y, Ghosh A, Hanash AM, Tran HT, Moore MA, van den Brink MR. Nrf2 regulates haematopoietic stem cell function. Nat Cell Biol. 2013;15(3):309–16. 10.1038/ncb2699.

28. Kim JH, Thimmulappa RK, Kumar V, Cui W, Kumar S, Kombairaju P, Zhang H, Margolick J, Matsui W, Macvittie T, Malhotra SV, Biswal S. NRF2-mediated Notch pathway activation enhances hematopoietic reconstitution following myelosuppressive radiation. J Clin Invest. 2014;124(2):730–41. 10.1172/JCI70812.

29. Hellmuth N, Brat C, Awad O, George S, Kahnt A, Bauer T, Huynh Phuoc HP, Steinhilber D, Angioni C, Hassan M, Hock KJ, Manolikakes G, Zacharowski K, Roos J, Maier TJ. Structural Modifications Yield Novel Insights Into the Intriguing Pharmacodynamic Potential of Anti-inflammatory Nitro-Fatty Acids. Front Pharmacol. 2021;12:715076. 10.3389/fphar.2021.715076.

30. Khoo NKH, Li L, Salvatore SR, Schopfer FJ, Freeman BA. Electrophilic fatty acid nitroalkenes regulate Nrf2 and NF-kappaB signaling:A medicinal chemistry investigation of structure-function relationships. Sci Rep. 2018;8(1):2295. 10.1038/s41598-018-20460-8.

31. Winters TA, Marzella L, Molinar-Inglis O, Price PW, Han NC, Cohen JE, Wang SJ, Fotenos AF, Sullivan JM, Esker JI, Lapinskas PJ, DiCarlo AL. Gastrointestinal Acute Radiation Syndrome: Mechanisms, Models, Markers, and Medical Countermeasures. Radiat Res. 2024;201(6):628–46. 10.1667/RADE-23-00196.1.

32. Xiao HW, Cui M, Li Y, Dong JL, Zhang SQ, Zhu CC, Jiang M, Zhu T, Wang B, Wang HC, Fan SJ. Gut microbiota-derived indole 3-propionic acid protects against radiation toxicity via retaining acyl-CoA-binding protein. Microbiome. 2020;8(1):69. 10.1186/s40168-020-00845-6.

33. Pejchal J, Tichy A, Kmochova A, Fikejzlova L, Kubelkova K, Milanova M, Lierova A, Filipova A, Muckova L, Cizkova J. Mitigation of Ionizing Radiation-Induced Gastrointestinal Damage by Insulin-Like Growth Factor-1 in Mice. Front Pharmacol. 2022;13:663855. 10.3389/fphar.2022.663855.

