## Supplementary file 1_SI Figures for "10-Nitro-oleic acid improves survival following gamma irradiation and modulates hematopoietic and intestinal responses"

Supplementary Figures:
Figure S1
Figure S2
Figure S3
Figure S4

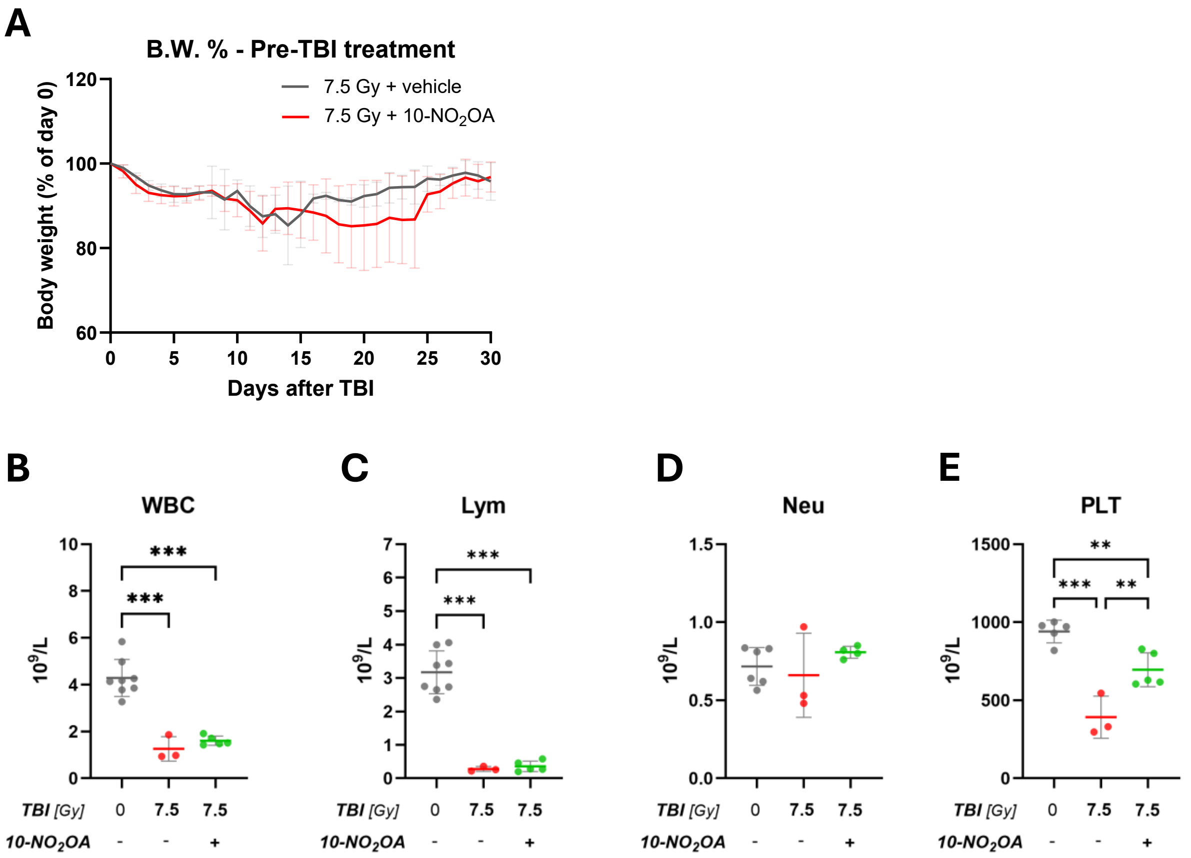


**Fig. S1: Effects of 10-$NO_2$OA pre-treatment on mice body weight and hematological profile after total body irradiation (TBI).** Two doses of 10-$NO_2$OA or vehicle (24 and 1 h before irradiation) were administered prior 7.5 Gy TBI. Body weight of C57BL/6 mice (A), the level of (B) WBC, (C) Lym, (D) Neu and (E) PLT were analyzed from peripheral blood of surviving mice 30 days after TBI (7.5 Gy). Data represent means ± SD, n = 9 (TBI + vehicle) and 10 (TBI + 10-$NO_2$OA) for Body weight, n = 3-7 for hematological analyses. Statistical analysis with one-way ANOVA with Tukey's multiple comparisons test; (**p < 0.01, ***p < 0.001). Total body irradiation (TBI); B.W. % – Body weight relative to day 0 in percentage; white blood cells (WBC); lymphocytes (Lym); neutrophils (Neu); platelets (PLT).

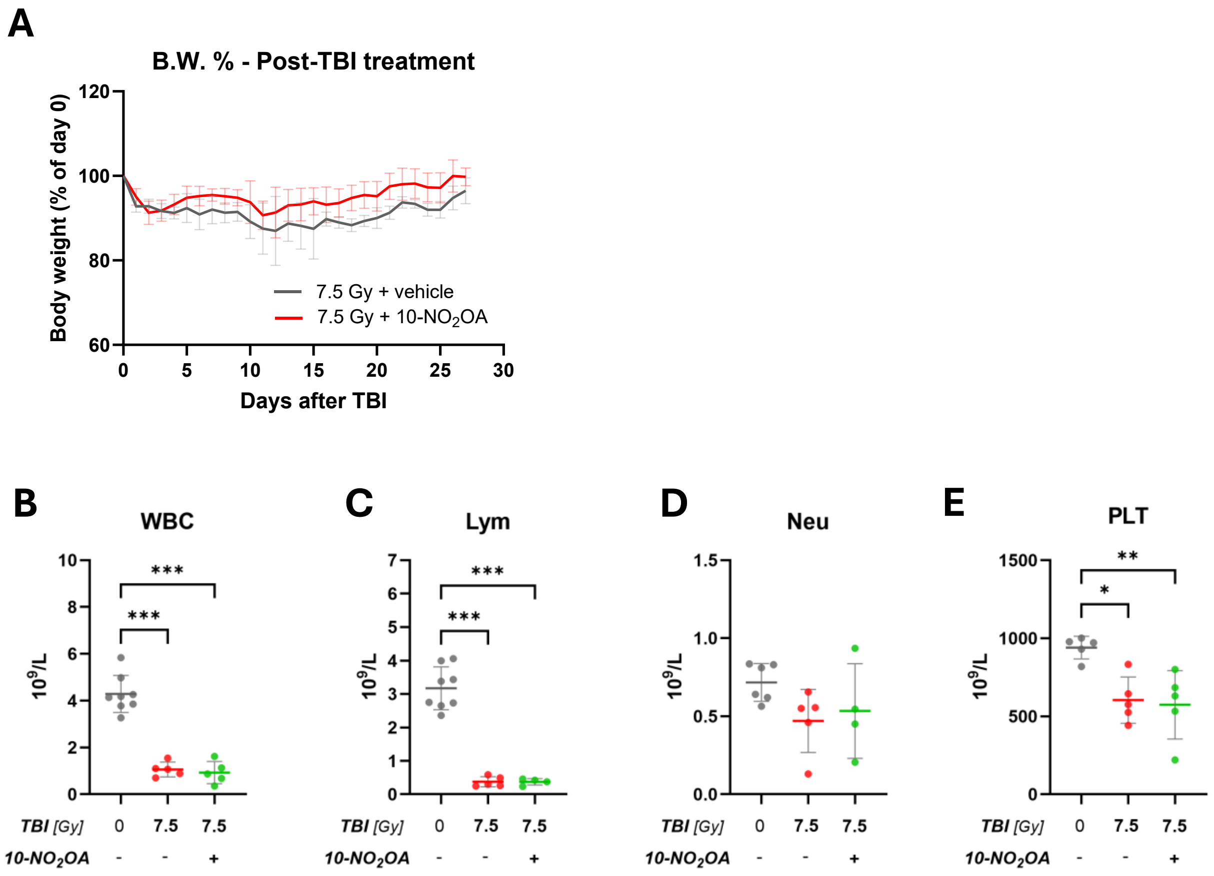


**Fig. S2: Effects of 10-$NO_2$OA post-treatment on mice body weight and hematological profile after total body irradiation (TBI).** Two doses of 10-$NO_2$OA or vehicle (4 and 48 h after irradiation) were administered post 7.5 Gy TBI. Body weight of C57BL/6 mice (A), the level of (B) WBC, (C) Lym, (D) Neu and (E) PLT were analyzed from peripheral blood of surviving mice 30 days after TBI (7.5 Gy). Data represent means ± SD, n = 7 (TBI + vehicle) and 9 (TBI + 10-$NO_2$OA) for Body weight, n = 4-8 for hematological analyses. Statistical analysis with one-way ANOVA with Tukey's multiple comparisons test; (*$p<0.05$, **$p < 0.01$, ***$p < 0.001$). Total body irradiation (TBI); B.W. % – Body weight relative to day 0 in percentage; white blood cells (WBC); lymphocytes (Lym); neutrophils (Neu); platelets (PLT).

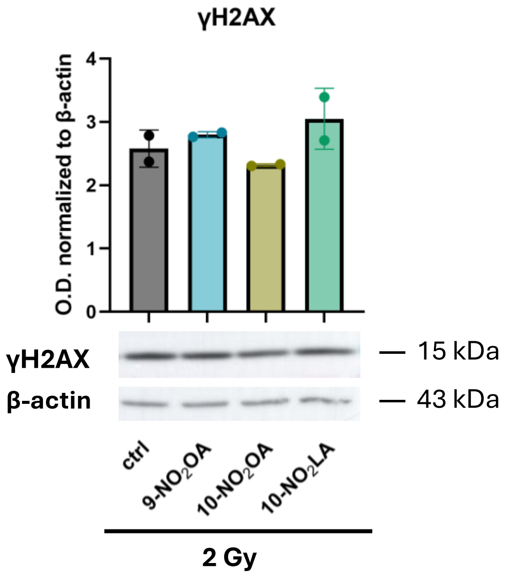


**Fig. S3: Effects of nitro-fatty acids on irradiation-induced γH2AX levels *in vitro*.** Isolated mouse bone marrow cells were treated with medium (control) or 2 µM 9-$NO_2$OA, 10-$NO_2$OA, or 10-$NO_2$LA for 1 h in 4% of oxygen condition and then irradiated with 2 Gy. Samples were collected 30 min after irradiation. γH2AX protein levels were assessed by Western blotting, band intensities were quantified by densitometry and normalized to β-actin. Data are representative of two independent experiments.

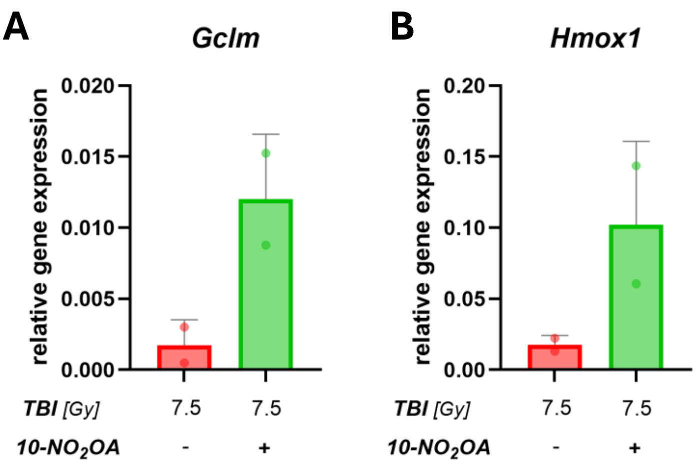


**Fig. S4: 10-NO$_2$OA pre-treatment enhances NRF2-responsive gene expression following total body irradiation (TBI).** C57BL/6 mice received 10-NO$_2$OA or vehicle 24 and 1 h before TBI (7.5 Gy). Bone marrow cells were isolated 3 h after TBI, and the relative expression of *Gclm* (A) and *Hmox1* (B) was assessed. 10-NO$_2$OA increased the expression of both genes compared with vehicle treatment. Data are representative of two independent experiments.
