## Supplementary file 2_Synthesis for "10-Nitro-oleic acid improves survival following gamma irradiation and modulates hematopoietic and intestinal responses"

### General information

All starting materials were purchased from commercial suppliers and used without further purification unless otherwise stated. Anhydrous tetrahydrofuran (THF), diethyl ether ($Et_2O$), and dichloromethane ($CH_2Cl_2$) were purchased from Sigma-Aldrich. Unless otherwise stated, reactions requiring an inert atmosphere were performed in round-bottom flasks fitted with rubber septa under a positive pressure of argon (Air Liquide, >99.5% purity).

Reaction products were purified by column chromatography using standard-grade silica gel (60 Å, 230–400 mesh) or by semipreparative HPLC using an Agilent 1290 Infinity II system equipped with a UV–Vis detector and an Agilent InfinityLab LC/MSD mass detector. Separations were performed on a reversed-phase C18 column (Agilent 5Prep-C18, 10 μm × 21.2 mm) using gradients of water and methanol at a flow rate of 15 or 20 mL/min.

Analytical thin-layer chromatography (TLC) was performed on aluminum-backed plates pre-coated with silica gel 60 $F^{254}$. Compounds were visualized under UV light or by staining with phosphomolybdic acid or potassium permanganate, followed by heating.

$^{1}H$ and $^{13}C\{^{1}H\}$ NMR spectra were recorded on JEOL ECA400II (400 MHz for $^{1}H$ and 101 MHz for $^{13}C$) or JEOL ECA500 (500 MHz for $^{1}H$ and 126 MHz for $^{13}C$) spectrometers using $CDCl_3$ as the solvent. Chemical shifts (δ) are reported in parts per million (ppm) and were referenced to the residual solvent signal at δ 7.26 ppm for $^{1}H$ NMR and to the central resonance of $CDCl_3$ at δ 77.16 ppm for $^{13}C\{^{1}H\}$ NMR. Proton signal multiplicities are abbreviated as follows: s (singlet), d (doublet), dd (doublet of doublets), t (triplet), tt (triplet of triplets), and m (multiplet).

High-resolution mass spectra (HRMS) were recorded on an Agilent 6230 high-resolution mass spectrometer equipped with an electrospray ionization (ESI) source and a time-of-flight (TOF) analyzer operating in positive- or negative-ion full-scan mode over the *m/z* range 100–1700. High-performance liquid chromatography (HPLC) analyses were performed using an Agilent 1290 Infinity II system equipped with a UV–Vis detector and an Agilent InfinityLab LC/MSD mass detector.

Lipase B *Candida antarctica* (CAL-B Lipase) immobilized from *Candida antarctica*, beads, white, >2 units/g, EC number 232-619-9, batch BCCH3575, was purchased from Sigma-Aldrich. Aqueous phosphate buffer (pH = 7.4) was prepared using “Quest Calculate™ PBS (Phosphate Buffered Saline) (1X, pH 7.4) Preparation and Recipe” protocol (https://www.aatbio.com/resources/buffer-preparations-and-recipes/pbs-phosphate-buffered-saline).

### Nitro fatty acid synthesis

#### **10-NO₂OA** synthesis

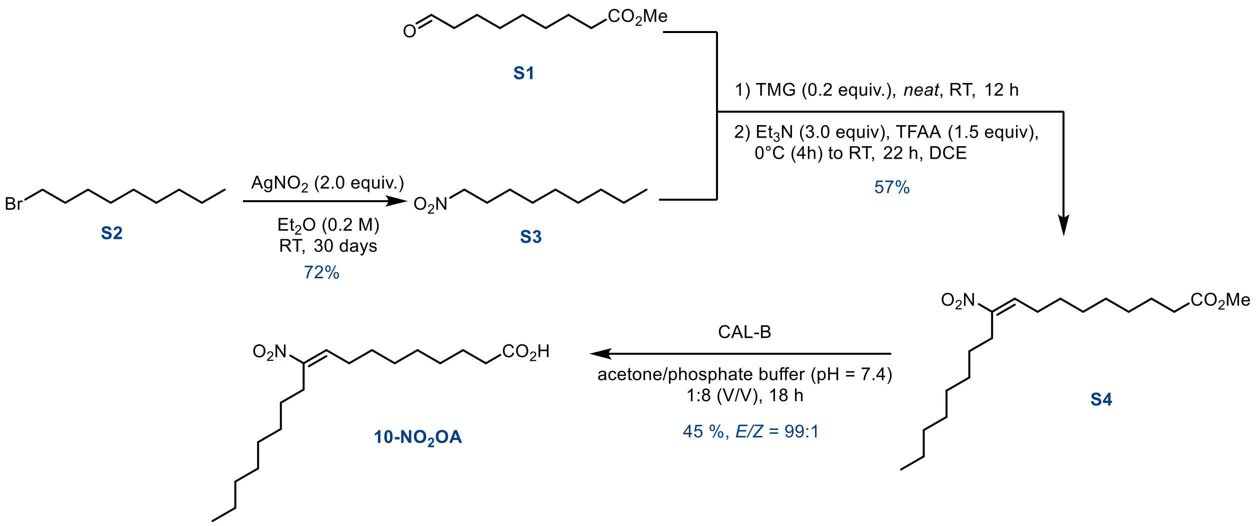

*1-nitrononane (**S3**)*

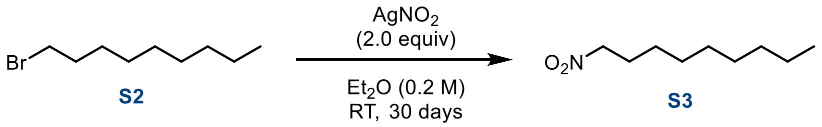

*The reaction flask was wrapped in aluminum foil to protect the reaction mixture from light.*

To a solution of bromide **S2** (4.0 g, 19.3 mmol, 1.0 equiv.) in $Et_2O$ (96 mL, 0.2 M) was added $AgNO_2$ (6.0 g, 38.6 mmol, 2.0 equiv.). The flask was sealed with a stopper and wrapped in aluminum foil to protect the reaction mixture from light. The resulting suspension was stirred at room temperature for 30 days. The reaction mixture was then diluted with EtOAc (50 mL) and filtered through a short pad of Celite®. The filter pad was washed with EtOAc (2 × 100 mL), and the combined filtrates were concentrated under reduced pressure. The crude product was purified by flash column chromatography ($SiO_2$, petroleum ether/EtOAc = 1000:1) to afford **S3** (2.43 g, 72%) as a colorless oil.

**$^1$H NMR (500 MHz, Chloroform-*d*) δ (ppm):** 4.37 (t, J = 7.1 Hz, 2H), 2.03 – 1.96 (m, 2H), 1.41 – 1.19 (m, 12H), 0.87 (t, J = 6.9 Hz, 3H).

**$^{13}$C {$^1$H} NMR (126 MHz, Chloroform-*d*) δ (ppm):** 75.9, 31.9, 29.4, 29.3, 29.0, 27.6, 26.4, 22.8, 14.2.

The spectroscopic data were consistent with those reported in the literature.[1]

*Methyl (9E)-10-nitrooctadec-9-enoate (**S4**)*

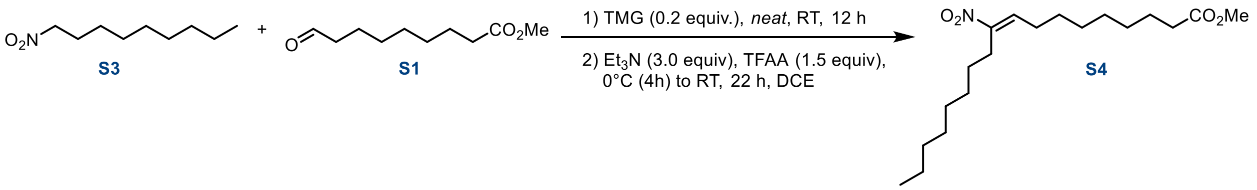

A neat mixture of 1-nitrononane **S3** (0.614 g, 3.54 mmol, 1.0 equiv.) and aldehyde S1 (0.800 g, 4.25 mmol, 1.2 equiv.) was cooled to 0 °C, and 1,1,3,3-tetramethylguanidine (TMG; 0.083 g, 0.71 mmol, 0.2 equiv.) was added. The resulting mixture was stirred at room temperature for 12 h, cooled to 0 °C, and quenched by the addition of $H_2O$ (15 mL). The mixture was extracted with EtOAc (3 × 30 mL), and the combined organic layers were dried over $Na_2SO_4$, filtered, and concentrated under reduced pressure. The crude product was purified by flash column chromatography ($SiO_2$, petroleum ether/EtOAc = 50:1) to afford the β-hydroxy adduct as a pale-yellow oil, which was used immediately in the subsequent step. $^1$H NMR (500 MHz, Chloroform-*d*) *δ* (ppm): 4.43 (ttd, *J* = 9.3, 4.0, 1.7 Hz, 1H), 4.00 (ddq, *J* = 8.7, 4.7, 2.0 Hz, 0.5H), 3.85 (pd, *J* = 7.3, 6.4, 2.8 Hz, 0.5H), 3.66 (s, 3H), 2.34 (dd, *J* = 4.9, 3.0 Hz, 0.5H), 2.30 (t, *J* = 7.5 Hz, 2H), 2.18 (dd, *J* = 8.0, 3.4 Hz, 0.5H), 2.14 – 1.95 (m, 1H), 1.83 – 1.72 (m, 1H), 1.63 – 1.59 (m, 2H), 1.53 – 1.40 (m, 3H), 1.33 – 1.23 (m, 18H), 0.87 (t, *J* = 7.0 Hz, 3H). The spectroscopic data were consistent with those reported in the literature.[2]

The resulting β-hydroxy adduct was dissolved in DCE (18 mL, 0.2 M) and cooled to 0 °C. $Et_3N$ (1.41 mL, 10.62 mmol, 3.0 equiv.) and trifluoroacetic anhydride (TFAA; 0.74 mL, 5.31 mmol, 1.5 equiv.) were added sequentially. The reaction mixture was stirred at 0 °C for 4 h and then at room temperature for an additional 22 h. The mixture was cooled to 0 °C and quenched by the addition of $H_2O$ (50 mL). The resulting mixture was extracted with EtOAc (3 × 75 mL), and the combined organic layers were dried over $MgSO_4$, filtered, and concentrated under reduced pressure. The crude product was purified by semipreparative reversed-phase HPLC (C18 column, MeOH/$H_2O$) to afford nitroalkene **S4** (0.870 g, 72%, E/Z >99:1) as a colorless oil. The E/Z ratio was determined by $^1$H NMR analysis.

**$^1$H NMR (500 MHz, Chloroform-*d*) *δ* (ppm):** 7.07 (t, *J* = 7.9 Hz, 1H), 3.67 (d, *J* = 1.0 Hz, 3H), 2.60 – 2.53 (m, 2H), 2.31 (t, *J* = 7.5 Hz, 2H), 2.21 (q, *J* = 7.6 Hz, 2H), 1.62 (p, *J* = 7.2 Hz, 2H), 1.48 (h, *J* = 8.8, 8.2 Hz, 4H), 1.38 – 1.23 (m, 16H), 0.88 (t, *J* = 7.0 Hz, 3H).

**$^{13}$C {$^1$H} NMR (126 MHz, Chloroform-*d*) *δ* (ppm):** 174.4, 152.1, 136.4, 51.6, 34.1, 32.0, 29.4, 29.32, 29.29, 29.14, 29.11, 28.6, 28.12, 28.05, 26.5, 25.0, 22.8, 14.2.

**HRMS (ESI) *m/z*:** $[M+H]^+$ calculated for $C_{19}H_{36}NO_4$: 342.2639; found: 342.2643.

*(9E)-10-nitrooctadec-9-enoic acid (**10-NO₂OA**)*

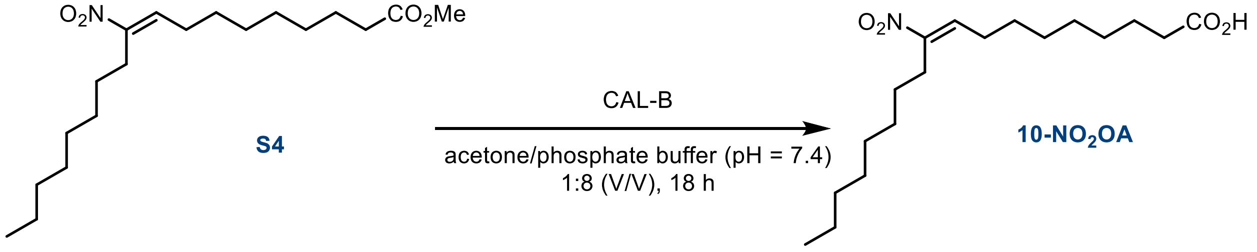

**CAL-B** (400 mg) was added to a solution of methyl ester **S4** (0.300 g, 0.88 mmol, 1.0 equiv.) in acetone (11 mL, 0.08 M) and aqueous phosphate buffer (88 mL, 0.01 M, pH 7.4). The resulting mixture was stirred vigorously (1000 rpm) at room temperature for 18 h. The pH was then adjusted to 3 using 1.0 M aqueous HCl, and the mixture was extracted with EtOAc (5 × 20 mL). The combined organic layers were dried over $MgSO_4$, filtered, and concentrated under reduced pressure. The crude product was purified by semipreparative reversed-phase HPLC (C18 column, MeOH/$H_2O$) to afford **10-$NO_2$OA** (0.130 g, 45%, *E/Z* >99:1) as a colorless oil. The E/Z ratio was determined by $^1$H NMR analysis.

**$^1$H NMR (500 MHz, Chloroform-*d*) *δ* (ppm):** 10.50 (broad s, 1H), 7.07 (t, *J* = 7.9 Hz, 1H), 2.59 – 2.54 (m, 2H), 2.36 (t, *J* = 7.5 Hz, 2H), 2.21 (q, *J* = 7.6 Hz, 2H), 1.67 – 1.60 (m, 2H), 1.53 – 1.44 (m, 4H), 1.39 – 1.32 (m, 6H), 1.32 – 1.24 (m, 10H), 0.88 (t, *J* = 6.8 Hz, 3H).

**$^{13}$C {$^1$H} NMR (126 MHz, Chloroform-*d*) *δ* (ppm):** 178.7, 152.1, 136.4, 33.8, 32.0, 29.4, 29.38, 29.32, 29.1, 29.0, 28.6, 28.12, 28.05, 26.5, 24.7, 22.8, 14.2.

**MS (ESI) *m/z* (%):** 327 [M-H]$^-$.

**HRMS (ESI) *m/z*:** [M-H]$^-$ calculated for $C_{18}H_{32}NO_4$: 326.2337; found: 326.2340.

#### 9-$NO_2$OA synthesis

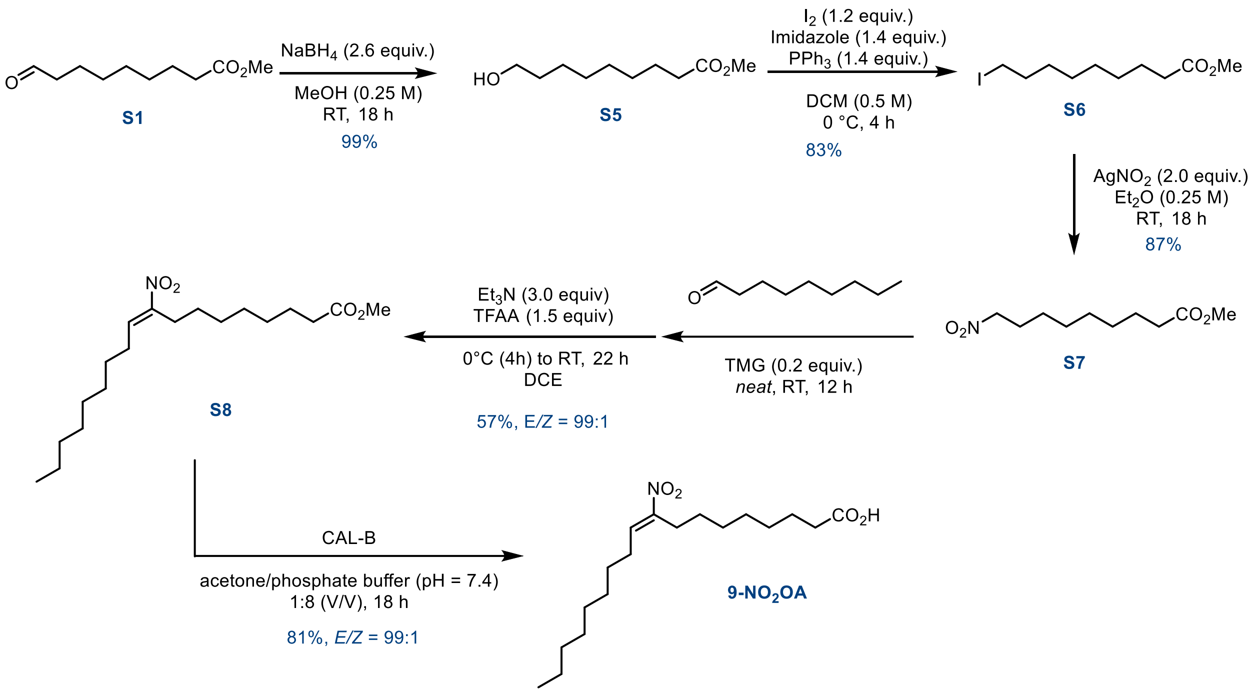

*Methyl 9-hydroxynonanoate (**S5**)*

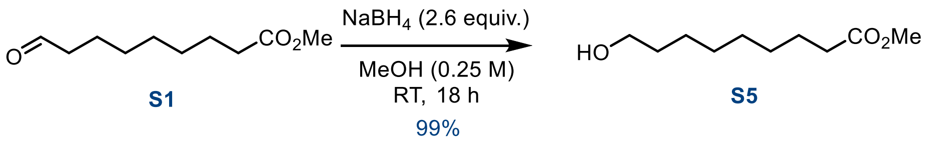

$NaBH_4$ (1.7 g, 44.2 mmol, 2.6 equiv.) was added portionwise to a cold (0 °C, ice-water bath) solution of aldehyde **S1** (3.2 g, 17 mmol, 1.0 equiv.) in MeOH (68 mL, 0.25 M). The resulting mixture was stirred at room temperature for 18 h. Water (50 mL) and EtOAc (100 mL) were added, and the layers were separated. The aqueous layer was extracted with EtOAc (3 × 50 mL), and the combined organic layers were dried over $MgSO_4$, filtered, and concentrated under reduced pressure. The crude product was purified by flash column chromatography ($SiO_2$; petroleum ether/EtOAc, 10:1) to afford alcohol **S5** (3.17 g, 99%) as a colorless oil.

**$^1$H NMR (500 MHz, Chloroform-*d*) *δ* (ppm):** 3.66 (d, *J* = 2.0 Hz, 3H), 3.63 (t, *J* = 6.6 Hz, 2H), 2.30 (td, *J* = 7.4, 2.0 Hz, 2H), 1.64 – 1.59 (m, 2H), 1.59 – 1.52 (m, 2H), 1.37 – 1.26 (m, 8H).

**$^{13}$C {$^1$H} NMR (126 MHz, Chloroform-*d*) *δ* (ppm):** 174.5, 63.2, 51.6, 34.2, 32.9, 29.32, 29.28, 29.17, 25.8, 25.0.

The spectroscopic data were consistent with those reported in the literature.[3]

*Methyl 9-iodononanoate (**S6**)*

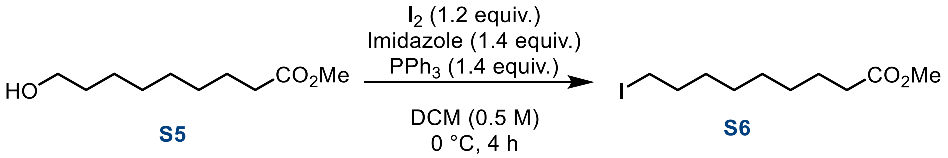

*The reaction flask was wrapped in aluminum foil to protect the reaction mixture from light.*

To a solution of alcohol **S5** (3.10 g, 16.5 mmol, 1.0 equiv.) in $CH_2Cl_2$ (33 mL, 0.5 M) at 0 °C (ice–water bath) were sequentially added imidazole (1.57 g, 22.8 mmol, 1.4 equiv.), $PPh_3$ (6.05 g, 22.8 mmol, 1.4 equiv.), and $I_2$ (5.00 g, 19.6 mmol, 1.2 equiv.). The reaction flask was wrapped in aluminum foil to protect the reaction mixture from light. The resulting mixture was stirred at 0 °C for 3.5 h and then diluted with EtOAc (100 mL). The mixture was filtered through a short pad of silica gel, and the filtrate was concentrated under reduced pressure. The crude product was purified by flash column chromatography ($SiO_2$, petroleum ether/EtOAc = 10:1) to afford iodide **S6** (4.08 g, 83%) as a pale-orange oil.

**$^1$H NMR (500 MHz, Chloroform-*d*) *δ* (ppm):** 3.67 (s, 3H), 3.18 (t, J = 7.0 Hz, 2H), 2.30 (t, J = 7.5 Hz, 2H), 1.81 (p, J = 7.1 Hz, 2H), 1.64 – 1.59 (m, 2H), 1.41 – 1.35 (m, 2H), 1.34 – 1.23 (m, 6H).

**$^{13}$C {$^1$H} NMR (126 MHz, Chloroform-*d*) *δ* (ppm):** 174.4, 51.6, 34.2, 33.6, 30.5, 29.17, 29.15, 28.5, 25.0, 7.4.

The spectroscopic data were consistent with those reported in the literature.[2]

*Methyl 9-nitrononanoate (**S7**)*

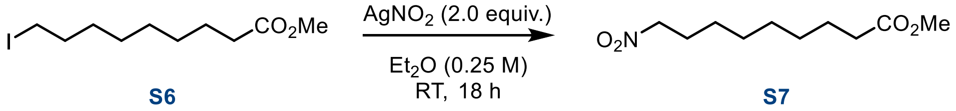

*The reaction flask was wrapped in aluminum foil to protect the reaction mixture from light.*

To a solution of iodide **S6** (4.00 g, 13.3 mmol, 1.0 equiv.) in $Et_2O$ (66 mL, 0.2 M) was added $AgNO_2$ (4.10 g, 26.6 mmol, 2.0 equiv.). The flask was sealed with a stopper, and the resulting suspension was stirred overnight at room temperature. The reaction mixture was diluted with EtOAc (100 mL) and filtered through a short pad of Celite®. The filter pad was washed with EtOAc (2 × 100 mL), and the combined filtrates were concentrated under reduced pressure. The crude product was purified by flash column chromatography ($SiO_2$, petroleum ether/EtOAc = 4:1) to afford nitroalkane **S7** (2.54 g, 87%) as a colorless oil.

**$^1$H NMR (500 MHz, Chloroform-*d*) *δ* (ppm):** 4.37 (t, *J* = 7.0 Hz, 2H), 3.66 (s, 3H), 2.30 (t, *J* = 7.5 Hz, 2H), 2.00 (p, *J* = 7.1 Hz, 2H), 1.62 (q, *J* = 7.4 Hz, 2H), 1.41 – 1.26 (m, 8H).

**$^{13}C$ {$^{1}H$} NMR (126 MHz, Chloroform-*d*) *δ* (ppm):** 174.4, 75.8, 51.62, 51.58, 34.1, 29.03, 29.01, 28.8, 27.5, 26.3, 24.9.

*Methyl (E)-9-nitrooctadec-9-enoate (**S8**)*

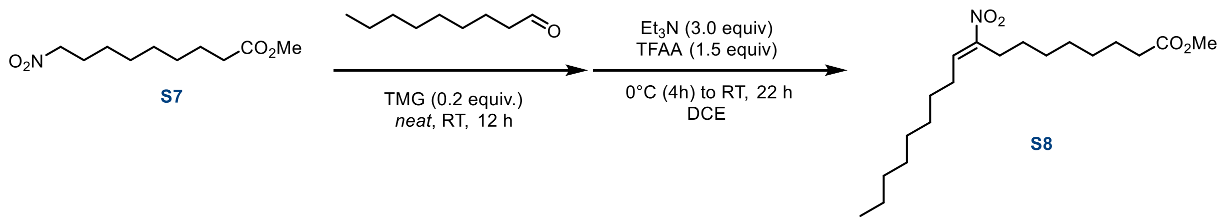

A mixture of nitroalkane **S7** (1.20 g, 5.47 mmol, 1.0 equiv.) and nonanal (0.96 g, 6.56 mmol, 1.2 equiv.) was cooled to 0 °C, and 1,1,3,3-tetramethylguanidine (TMG; 0.14 g, 1.1 mmol, 0.2 equiv.) was added. The resulting mixture was stirred at room temperature for 12 h, cooled to 0 °C, and quenched by the addition of $H_2O$ (25 mL). The mixture was extracted with EtOAc (3 × 30 mL), and the combined organic layers were dried over $MgSO_4$, filtered, and concentrated under reduced pressure. The resulting mixture was dissolved in DCE (27 mL, 0.2 M) and cooled to 0 °C. $Et_3N$ (2.28 mL, 16.4 mmol, 3.0 equiv.) and TFAA (1.14 mL, 8.2 mmol, 1.5 equiv.) were added sequentially. The resulting mixture was stirred at 0 °C for 4 h and then at room temperature for an additional 22 h. The reaction mixture was cooled to 0 °C (ice–water bath) and quenched by the addition of $H_2O$ (75 mL). The mixture was extracted with EtOAc (3 × 100 mL), and the combined organic layers were dried over $MgSO_4$, filtered, and concentrated under reduced pressure. The crude product was purified by semipreparative reversed-phase HPLC (C18 column, MeOH/$H_2O$) to afford methyl (*E*)-9-nitrooleate **S8** (1.06 g, 57%, *E/Z* >99:1). The *E/Z* ratio was determined by $^{1}H$ NMR analysis.

**$^{1}H$ NMR (500 MHz, Chloroform-*d*) *δ* (ppm):** 7.08 (t, *J* = 7.9 Hz, 1H), 3.66 (s, 3H), 2.60 – 2.53 (m, 2H), 2.30 (t, *J* = 7.5 Hz, 2H), 2.21 (q, *J* = 7.6 Hz, 2H), 1.61 (p, *J* = 7.2 Hz, 2H), 1.48 (h, *J* = 7.5 Hz, 4H), 1.37 – 1.22 (m, 16H), 0.90 – 0.83 (m, 3H).

**$^{13}C$ {$^{1}H$} NMR (126 MHz, Chloroform-*d*) *δ* (ppm):** 174.4, 151.9, 136.7, 51.6, 34.2, 31.9, 29.5, 29.4, 29.3, 29.2, 29.12, 29.05, 28.7, 28.2, 28.0, 26.5, 25.0, 22.8, 14.2.

**HRMS (ESI) *m/z:*** $[M+H]^+$ calculated for $C_{19}H_{36}NO_4$: 342.2639; found: 342.2638.

*(9E)-9-nitrooctadec-9-enoic acid (**9-NO₂OA**)*

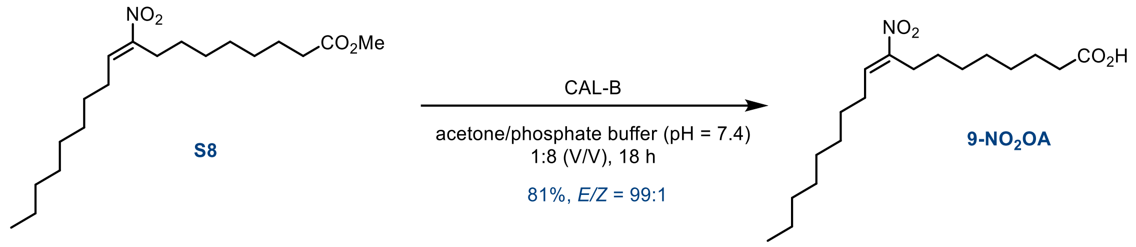

CAL-B (400 mg) was added to a solution of methyl ester **S8** (0.130 g, 0.38 mmol, 1.0 equiv.) in acetone (4.7 mL, 0.08 M) and aqueous phosphate buffer (38 mL, 0.01 M, pH 7.4). The resulting mixture was stirred vigorously (1000 rpm) at room temperature for 18 h. The pH was then adjusted to 3 using 1.0 M aqueous HCl, and the mixture was extracted with EtOAc (5 × 20 mL). The combined organic layers were dried over $MgSO_4$, filtered, and concentrated under reduced pressure. The crude product was purified by semipreparative reversed-phase HPLC (C18 column, MeOH/$H_2O$) to afford **9-$NO_2$OA** (0.100 g, 81%, E/Z ≥99:1). The *E/Z* ratio was determined by $^1$H NMR analysis.

**$^1$H NMR (500 MHz, Chloroform-*d*) *δ* (ppm):** 11.53 (broad s, 1H), 7.07 (t, *J* = 7.9 Hz, 1H), 2.60 – 2.51 (m, 2H), 2.34 (t, *J* = 7.5 Hz, 2H), 2.20 (q, *J* = 7.6 Hz, 2H), 1.62 (p, *J* = 7.4 Hz, 2H), 1.48 (h, *J* = 7.2 Hz, 4H), 1.37 – 1.29 (m, 6H), 1.32 – 1.21 (m, 10H), 0.92 – 0.83 (m, 3H).

**$^{13}$C {$^1$H} NMR (126 MHz, Chloroform-*d*) *δ* (ppm):** 178.0, 152.1, 136.4, 33.9, 32.0, 29.4, 29.32, 29.27, 29.1, 29.0, 28.6, 28.11, 28.05, 26.5, 24.7, 22.8, 14.2.

**HRMS (ESI) *m/z:*** [M-H]$^-$ calculated for $C_{18}H_{32}NO_4$: 326.2337; found: 326.2326.

#### 10-$NO_2$LA synthesis

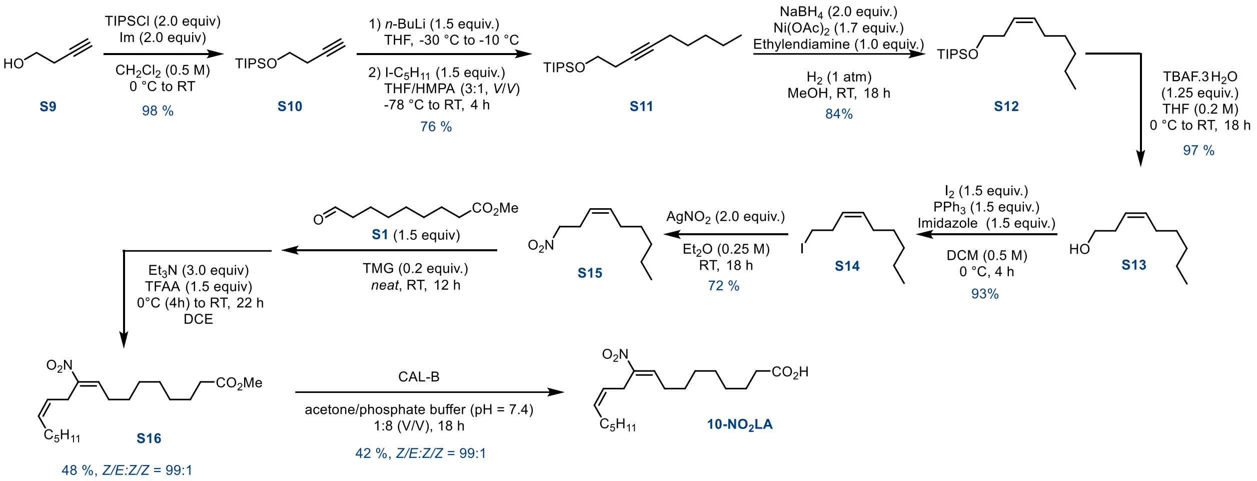

*(But-3-yn-1-yloxy)triisopropylsilane (**S10**)*

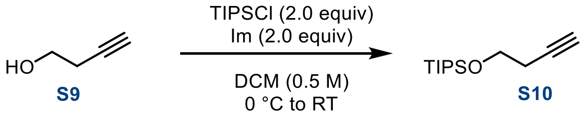

A solution of imidazole (18.2 g, 263 mmol, 2.0 equiv.) in $CH_2Cl_2$ (250 mL, 0.5 M) was cooled to 0 °C, and but-3-yn-1-ol **S9** (10 mL, 131 mmol, 1.0 equiv.) was added. After 5 min, TIPSCl (59 mL, 263 mmol, 2.0 equiv.) was added over 5 min, and the resulting mixture was stirred at room temperature for 16 h. $H_2O$ (50 mL) was then added, and the layers were separated. The aqueous layer was extracted with $CH_2Cl_2$ (3 × 150 mL), and the combined organic layers were washed with brine (100 mL), dried over $Na_2SO_4$, filtered, and concentrated under reduced pressure. The crude product was purified by flash column chromatography ($SiO_2$, petroleum ether/EtOAc = 100:1) to afford silyl ether **S10** (29.1 g, 98%) as a colorless oil.

**$^1$H NMR (500 MHz, Chloroform-*d*) δ (ppm):** 3.82 (t, *J* = 7.3 Hz, 2H), 2.44 (td, *J* = 7.3, 2.7 Hz, 2H), 1.96 (t, *J* = 2.7 Hz, 1H), 1.10 – 1.03 (m, 21H).

**$^{13}$C {$^1$H} NMR (126 MHz, Chloroform-*d*) δ (ppm):** 81.7, 69.4, 62.1, 23.0, 18.1, 12.1.

**MS (ESI) *m/z* (%):** 227 [M+H]$^+$.

**$R_f$** = 0.45 (petroleum ether/EtOAc = 10:1; visualized with vanillin, green spot).

The analytical data were consistent with those previously reported.[4]

*Triisopropyl(oct-3-yn-1-yloxy)silane (**S11**)*

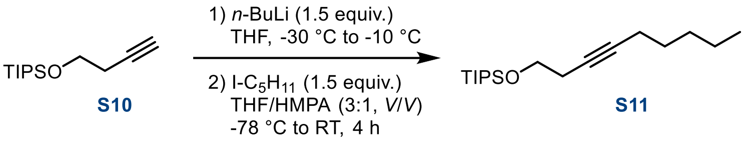

A solution of TIPS-protected alkyne **S10** (10.0 g, 43.7 mmol, 1.0 equiv.) in THF (109 mL, 0.4 M) was cooled to −30 °C (dry ice/acetone bath, external temperature), and *n*-BuLi (41 mL, 64 mmol, 1.5 equiv.; 1.6 M solution in hexanes) was added dropwise. The resulting mixture was warmed to −10 °C (external temperature) and stirred for 1 h. The reaction mixture was then cooled to −78 °C (dry ice/acetone bath, external temperature), and a solution of 1-iodopentane (8.7 mL, 65.6 mmol, 1.5 equiv.) in THF/HMPA (3:1, V/V; 88 mL) was added dropwise. The resulting mixture was stirred at −78 °C for 1 h and then allowed to warm to room temperature over 4 h. $H_2O$ (50 mL) was added, and the mixture was extracted with $Et_2O$ (3 × 100 mL). The combined organic layers were washed with brine (50 mL), dried over $Na_2SO_4$, filtered, and concentrated under reduced pressure. The crude product was purified by flash column chromatography ($SiO_2$, petroleum ether/EtOAc = 100:1) to afford alkyne **S11** (9.86 g, 76%) as a colorless oil.

**$^1$H NMR (500 MHz, Chloroform-*d*) δ (ppm):** 3.77 (t, *J* = 7.4 Hz, 2H), 2.42 – 2.36 (m, 2H), 2.13 (tt, *J* = 7.2, 2.4 Hz, 2H), 1.48 (p, *J* = 7.0 Hz, 2H), 1.39 – 1.27 (m, 3H), 1.14 – 1.01 (m, 20H), 0.90 (t, *J* = 7.1 Hz, 3H).

**$^{13}$C {$^1$H} NMR (126 MHz, Chloroform-*d*) δ (ppm):** 81.6, 77.1, 62.9, 31.3, 28.95, 23.5, 22.5, 19.0, 18.2, 14.2, 12.2.

**HRMS (ESI) *m/z:*** $[M+H]^+$ calculated for $C_{18}H_{39}OSi$: 297.2608; found: 297.2605.

**R*f*** = = 0.45 (petroleum ether/EtOAc = 10:1; visualized with vanillin, orange spot).

*(Z)-triisopropyl(non-3-en-1-yloxy)silane (**S12**)*

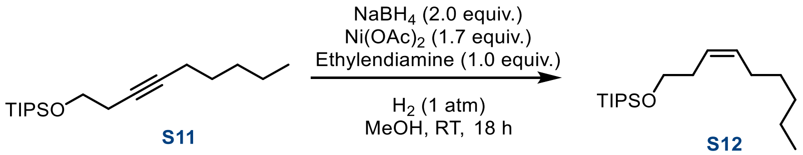

$NaBH_4$ (2.49 g, 65.8 mmol, 2.0 equiv.) was added to a stirred solution of $Ni(OAc)_2$ (14.2 g, 56 mmol, 1.7 equiv.) in anhydrous MeOH (330 mL, 0.1 M) at room temperature under an argon atmosphere. The argon atmosphere (balloon) was replaced with hydrogen (double-layer balloon), and ethylenediamine (3.29 mL, 32.9 mmol, 1.0 equiv.) was added. After 5 min, a solution of alkyne **S11** (9.86 g, 32.9 mmol, 1.0 equiv.) in MeOH (20 mL) was added via cannula, and the resulting mixture was stirred for 4 h. The reaction mixture was filtered through a pad of Celite®, and the filter pad was washed with $Et_2O$ (200 mL). The combined filtrates were diluted with brine (150 mL) and EtOAc (150 mL), and the layers were separated. The aqueous layer was extracted with EtOAc (3 × 150 mL), and the combined organic layers were washed with brine (50 mL), dried over $MgSO_4$, filtered, and concentrated under reduced pressure. The crude product was purified by flash column chromatography ($SiO_2$, petroleum ether/EtOAc = 100:1) to afford alkene **S12** (8.25 g, 84%, *E/Z* = 1:>99) as a colorless oil.

**$^1$H NMR (500 MHz, Chloroform-*d*) δ (ppm):** 5.48 – 5.35 (m, 2H), 3.68 – 3.65 (m, 2H), 2.30 (q, *J* = 7.2 Hz, 2H), 2.04 (q, *J* = 7.3 Hz, 2H), 1.35 – 1.27 (m, 6H), 1.06 (dd, *J* = 5.5, 2.3 Hz, 21H), 0.90 – 0.86 (m, 3H).

**$^{13}$C{$^1$H} NMR (126 MHz, Chloroform-*d*) δ (ppm):** 132.0, 125.7, 63.4, 31.7, 31.4, 29.6, 27.5, 22.7, 18.2, 14.2, 12.2.

**HRMS (ESI) *m/z:*** $[M+H]^+$ calculated for $C_{18}H_{39}OSi$: 299.2765; found: 299.2768.

**R*f*** = 0.55 (petroleum ether/EtOAc = 10:1; visualized with PMA).

*(Z)-non-3-en-1-ol (**S13**)*

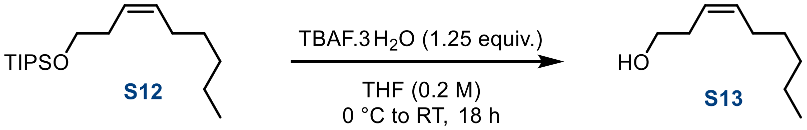

Alkene **S12** (5.00 g, 16.6 mmol, 1.0 equiv.) was dissolved in THF (83 mL, 0.2 M), and the resulting solution was cooled to 0 °C (ice–water bath, external temperature). Solid TBAF·3$H_2O$ (6.67 g, 20.7 mmol, 1.25 equiv.) was added portionwise, and the resulting mixture was stirred at 0 °C for 3 h. Saturated aqueous $NH_4Cl$ (50 mL) was added, and the mixture was extracted with EtOAc (3 × 100 mL). The combined organic layers were washed with brine (50 mL), dried over $MgSO_4$, filtered, and concentrated under reduced pressure. The crude product was purified by flash column chromatography ($SiO_2$, petroleum ether/EtOAc = 10:1) to afford alcohol **S13** (2.30 g, 97%) as a colorless oil.

**$^1$H NMR (500 MHz, Chloroform-*d*) δ (ppm):** 5.62 – 5.51 (m, 1H), 5.42 – 5.29 (m, 1H), 3.67 – 3.57 (m, 2H), 2.33 (qd, *J* = 6.6, 1.5 Hz, 2H), 2.06 (qd, *J* = 7.3, 1.6 Hz, 2H), 1.39 – 1.23 (m, 7H), 0.88 (t, *J* = 6.9 Hz, 3H).

**$^{13}$C {$^1$H} NMR (126 MHz, Chloroform-*d*) δ (ppm):** 133.8, 125.1, 62.5, 31.6, 30.9, 29.5, 27.5, 22.7, 18.7, 14.2.

**MS (ESI) *m/z* (%):** 143 $[M+H]^+$.

**HRMS (ESI) *m/z:*** $[M+H]^+$ calculated for $C_9H_{19}O$: 143.1430; found: 143.1429.

**R*f*** = = 0.30 (petroleum ether/EtOAc = 4:1; visualized with vanillin, blue-green spot).

*(Z)-1-iodonon-3-ene (**S14**)*

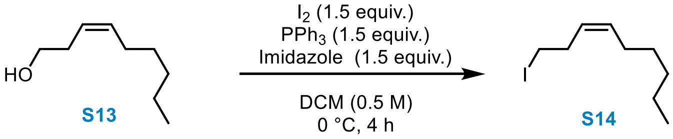

*The reaction flask was wrapped in aluminum foil to protect the reaction mixture from light.*

A solution of alcohol **S13** (2.11 g, 14.7 mmol, 1.0 equiv.) in $CH_2Cl_2$ (147 mL, 0.1 M) was cooled to 0 °C (ice–water bath, external temperature), and $PPh_3$ (5.84 g, 22.0 mmol, 1.5 equiv.), imidazole (1.51 g, 22.0 mmol, 1.5 equiv.), and $I_2$ (5.65 g, 22.0 mmol, 1.5 equiv.) were added sequentially. The resulting mixture was allowed to warm to room temperature and stirred for 4 h. $H_2O$ (50 mL) was added to quench the reaction, and the layers were separated. The aqueous layer was extracted with EtOAc (3 × 50 mL), and the combined organic layers were washed with brine (50 mL), dried over $MgSO_4$, filtered, and concentrated under reduced pressure. The crude product was purified by flash column chromatography ($SiO_2$, petroleum ether/EtOAc = 10:1) to afford iodide **S14** (3.45 g, 93%) as a colorless oil.

**$^1$H NMR (500 MHz, Chloroform-*d*) δ (ppm):** 5.57 – 5.50 (m, 1H), 5.34 – 5.28 (m, 1H), 3.13 (t, *J* = 7.3 Hz, 2H), 2.63 (q, *J* = 7.3 Hz, 2H), 2.02 (q, *J* = 7.4 Hz, 2H), 1.38 – 1.25 (m, 6H), 0.89 (t, *J* = 6.8 Hz, 3H).

**$^{13}$C{$^1$H} NMR (126 MHz, Chloroform-*d*) δ (ppm):** 132.9, 127.8, 31.7, 31.6, 29.3, 27.6, 22.7, 14.2, 5.7.

**MS (ESI) *m/z* (%):** 253 $[M+H]^+$.

**HRMS (ESI) *m/z:*** $[M+H]^+$ calculated for $C_9H_{18}I$: 253.0448; found: 253.0450.

**R*f*** = 0.80 (petroleum ether/EtOAc = 10:1; visualized with vanillin).

*(Z)-1-nitronon-3-ene (**S15**)*

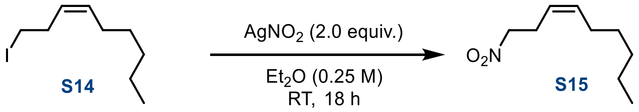

*The reaction flask was wrapped in aluminum foil to protect the reaction mixture from light.*

At room temperature, silver nitrite (4.2 g, 27.1 mmol, 2.0 equiv.) was added in one portion to a stirred solution of **S14** (3.45 g, 13.5 mmol, 1.0 equiv.) in $Et_2O$ (54 mL, 0.25 M). The resulting mixture was stirred at room temperature for 18 h and then filtered through a short pad of Celite®. The filter cake was washed with EtOAc (3 × 50 mL), and the combined organic layers were washed with brine, dried over $MgSO_4$, filtered, and concentrated under reduced pressure. The crude product was purified by flash column chromatography ($SiO_2$; petroleum ether/EtOAc, 10:1) to afford the desired nitro compound **S15** (1.67 g, 72%).

**$^1$H NMR (500 MHz, Chloroform-*d*) *δ* (ppm):** 5.58 (dtt, *J* = 10.6, 7.5, 1.5 Hz, 1H), 5.29 (dtt, *J* = 10.7, 7.5, 1.8 Hz, 1H), 4.37 (t, *J* = 7.2 Hz, 2H), 2.83 – 2.68 (m, 2H), 2.08 – 2.01 (m, 2H), 1.36 – 1.25 (m, 6H), 0.89 (t, *J* = 6.9 Hz, 3H).

**$^{13}$C {$^1$H} NMR (126 MHz, Chloroform-*d*) *δ* (ppm):** 135.1, 122.2, 75.3, 31.6, 29.2, 27.4, 25.6, 22.7, 14.2.

**MS (ESI) *m/z* (%):** 172 $[M+H]^+$.

**HRMS (ESI) *m/z:*** $[M+H]^+$ calculated for $C_9H_{18}NO_2$: 172.1332; found: 172.1335.

**R*f*** = 0.40 (petroleum ether/EtOAc, 10:1).

*Methyl (9E,12Z)-10-nitrooctadeca-9,12-dienoate (**S16**)*

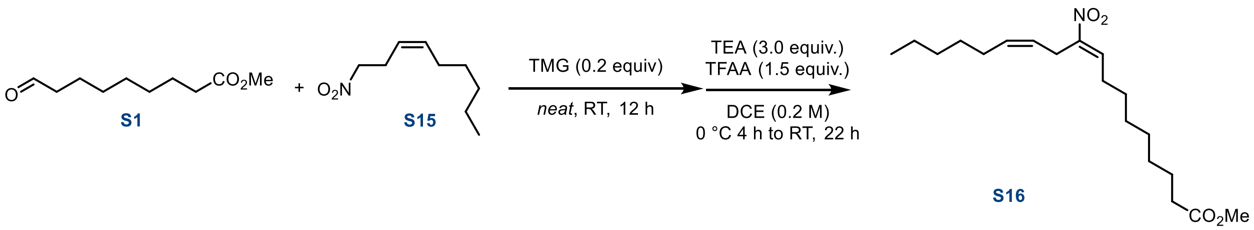

A neat mixture of nitroalkane **S15** (0.300 g, 1.73 mmol, 1.0 equiv.) and aldehyde **S1** (0.49 g, 2.60 mmol, 1.5 equiv.) was cooled to 0 °C (ice–water bath, external temperature), and 1,1,3,3-tetramethylguanidine (TMG; 0.040 g, 0.35 mmol, 0.2 equiv.) was added. The resulting mixture was stirred at room temperature for 12 h, cooled to 0 °C, and quenched by the addition of $H_2O$ (5 mL). The mixture was extracted with EtOAc (3 × 15 mL), and the combined organic layers were dried over $MgSO_4$, filtered, and concentrated under reduced pressure. The crude product was purified by flash column chromatography ($SiO_2$, petroleum ether/EtOAc = 50:1) to afford the β-hydroxy adduct (0.500 g, 81%, d.r. = 1.2:1), which was used in the subsequent step. $^1$H NMR (500 MHz, Chloroform-*d*) *δ* (ppm): 5.62 – 5.53 (m, 1H), 5.33 – 5.24 (m, 1H), 4.48 – 4.39 (m, 1H), 4.05 (dq, *J* = 8.5, 4.2 Hz, 0.5H, *major diastereoisomer*), 3.91 – 3.85 (m, 0.5H, *minor diastereoisomer*), 3.66 (s, 3H), 2.94 – 2.76 (m, 1H), 2.62 – 2.52 (m, 1H), 2.36 (d, *J* = 4.8 Hz, 0.5H), 2.30 (t, *J* = 7.5 Hz, 2H), 2.21 (d, *J* = 8.2 Hz, 0.5H), 2.02 (q, *J* = 7.4 Hz, 2H), 1.62 (p, *J* = 7.3 Hz, 2H), 1.58 – 1.39 (m, 4H), 1.34 – 1.26 (m, 12H), 0.88 (t, *J* = 6.9 Hz, 3H); $R_f$ = 0.20 (petroleum ether/EtOAc = 10:1; visualized with PMA).

The resulting β-hydroxy adduct was dissolved in DCE (7 mL, 0.2 M) and cooled to 0 °C (ice–water bath, external temperature). $Et_3N$ (0.59 mL, 4.2 mmol, 3.0 equiv.) and TFAA (0.30 mL, 2.1 mmol, 1.5 equiv.) were added sequentially. The reaction mixture was stirred at 0 °C for 4 h and then at room temperature for an additional 22 h. The mixture was cooled to 0 °C and quenched by the addition of $H_2O$ (10 mL). The resulting mixture was extracted with EtOAc (3 × 25 mL), and the combined organic layers were dried over $MgSO_4$, filtered, and concentrated under reduced pressure. The crude product was purified by flash column chromatography ($SiO_2$, petroleum ether/EtOAc = 20:1) to afford nitroalkene **S16** (0.282 g, 48%, E/Z ≥99:1) as a colorless oil. The *E/Z* ratio was determined by $^1$H NMR analysis.

**$^1$H NMR (500 MHz, Chloroform-*d*) *δ* (ppm):** 7.08 (t, J = 7.9 Hz, 1H), 5.49 (dtt, J = 10.8, 7.3, 1.8 Hz, 1H), 5.25 (dtt, J = 10.5, 7.0, 1.7 Hz, 1H), 3.66 (s, 3H), 3.33 (dd, J = 7.0, 1.8 Hz, 2H), 2.30 (t, J = 7.5 Hz, 2H), 2.24 (q, J = 7.7 Hz, 2H), 2.12 (qd, J = 7.3, 1.6 Hz, 2H), 1.62 (dd, J = 9.9, 4.6 Hz, 2H), 1.49 (dt, J = 11.5, 7.2 Hz, 2H), 1.41 – 1.23 (m, 12H), 0.91 – 0.87 (m, 3H).

**$^{13}$C {$^1$H} NMR (126 MHz, Chloroform-*d*) *δ* (ppm):** 174.3, 150.8, 136.6, 133.2, 123.4, 51.6, 34.1, 31.7, 29.29, 29.23, 29.12, 29.10, 28.5, 28.1, 27.5, 25.0, 22.7, 14.2.

**MS (ESI) *m/z* (%):** 340 $[M+H]^+$ (100).

**HRMS (ESI) *m/z*:** $[M+H]^+$ calculated for $C_{19}H_{34}NO_4$: 340.2482; found: 340.2486.

*(9E,12Z)-10-nitrooctadeca-9,12-dienoic acid (**10-NO₂LA**)*

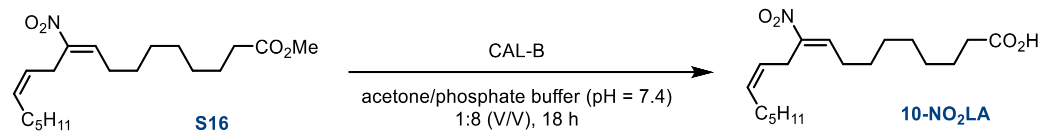

CAL-B (400 mg) was added to a solution of methyl ester **S16** (0.081 g, 0.24 mmol, 1.0 equiv.) in acetone (3 mL, 0.080 M) and aqueous phosphate buffer (24 mL, 0.01 M, pH 7.4). The resulting mixture was stirred vigorously (1000 rpm) at room temperature for 18 h. The pH was then adjusted to 3 using 1.0 M aqueous HCl, and the mixture was extracted with EtOAc (5 × 20 mL). The combined organic layers were dried over $MgSO_4$, filtered, and concentrated under reduced pressure. The crude product was purified by semipreparative reversed-phase HPLC (C18 column, MeOH/$H_2O$) to afford **10-$NO_2$LA** (0.033 g, 42%, *E,Z/Z,Z* ≥99:1). The stereoisomeric ratio was determined by $^1$H NMR analysis.

**$^{1}$H NMR (500 MHz, Chloroform-*d*) *δ* (ppm):** 11.14 (bs, 1H), 7.08 (t, *J* = 7.9 Hz, 1H), 5.49 (dtt, *J* = 10.9, 7.3, 1.8 Hz, 1H), 5.25 (dtt, *J* = 10.7, 6.9, 1.7 Hz, 1H), 3.34 (dd, *J* = 7.0, 1.8 Hz, 2H), 2.35 (t, *J* = 7.4 Hz, 2H), 2.24 (q, *J* = 7.6 Hz, 2H), 2.12 (q, *J* = 7.3 Hz, 2H), 1.65 – 1.61 (m, 2H), 1.53 – 1.47 (m, 2H), 1.38 – 1.29 (m, 12H), 0.90 (t, *J* = 6.8 Hz, 3H).

**$^{13}$C {$^{1}$H} NMR (126 MHz, Chloroform-*d*) *δ* (ppm):** 179.6, 150.8, 136.6, 133.2, 123.4, 34.0, 31.7, 29.3, 29.2, 29.1, 29.0, 28.5, 28.1, 27.6, 25.0, 24.7, 22.7, 14.2.

**MS (ESI) *m/z* (%):** 324 $[M-H]^-$ (100).

**HRMS (ESI) *m/z:*** $[M-H]^-$ calculated for $C_{18}H_{30}NO_4$: 324.2169; found: 324.2179.

#### 9-NO₂cLA synthesis

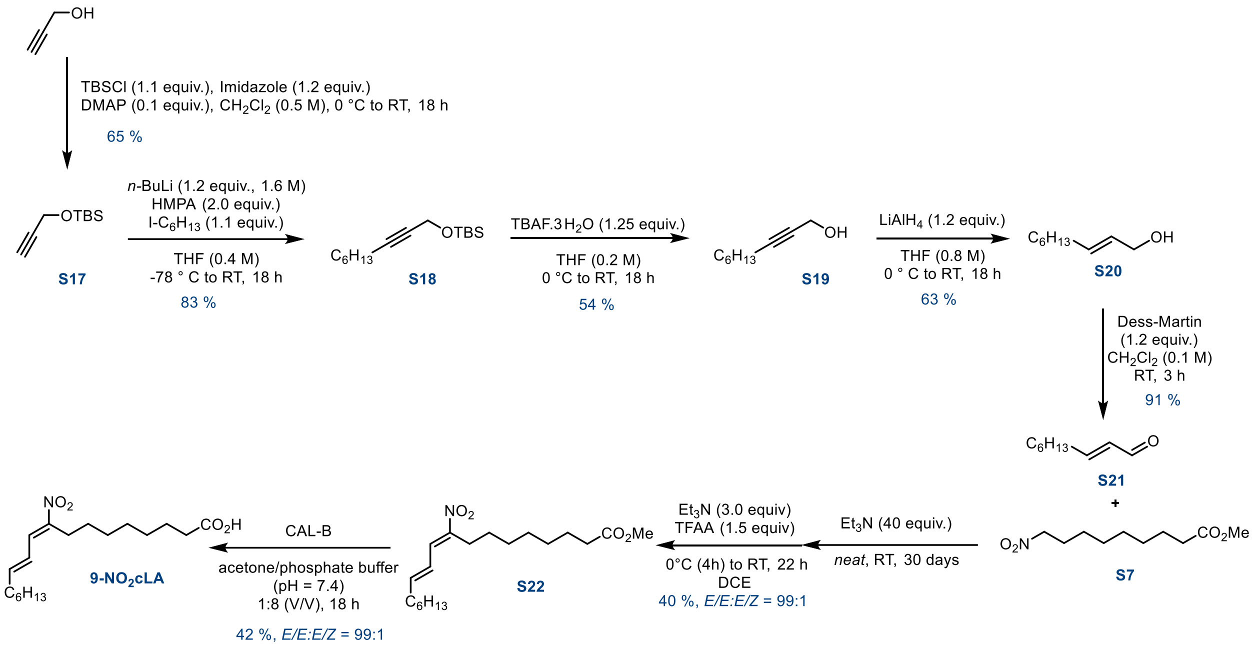

*tert-butyldimethyl(prop-2-yn-1-yloxy)silane (**S17**)*

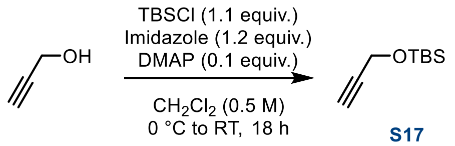

A solution of imidazole (22.0 g, 323 mmol, 1.2 equiv.) in $CH_2Cl_2$ (539 mL, 0.5 M) was cooled to 0 °C (ice–water bath, external temperature), and propargyl alcohol (15.1 g, 269 mmol, 1.0 equiv.) was added. DMAP (3.36 g, 26.9 mmol, 0.1 equiv.) and TBSCl (44.6 g, 296 mmol, 1.1 equiv.) were then added sequentially, and the resulting mixture was stirred at room temperature for 18 h. Saturated aqueous $NH_4Cl$ (150 mL) was added, and the layers were separated. The aqueous layer was extracted with $CH_2Cl_2$ (3 × 150 mL), and the combined organic layers were washed with brine (100 mL), dried over $Na_2SO_4$, filtered, and concentrated under reduced pressure. The crude product was purified by flash column chromatography ($SiO_2$, petroleum ether/EtOAc = 100:1) to afford TBS ether S17 (30.0 g, 65%) as a colorless oil.

**$^1$H NMR (500 MHz, Chloroform-*d*) *δ* (ppm):** 4.31 (m, 2H), 2.39 (td, *J* = 2.4, 0.8 Hz, 1H), 0.91 (m, 9H), 0.13 – 0.12 (m, 6H).

**$^{13}$C {$^1$H} NMR (126 MHz, Chloroform-*d*) *δ* (ppm):** 82.5, 73.0, 51.7, 26.1, 25.9, 18.4, -5.1.

**MS (ESI) *m/z* (%):** 171 [M+H]$^+$ (100).

**$R_f$** = 0.55 (hexane/EtOAc = 10:1).

*tert-Butyldimethyl(oct-2-yn-1-yloxy)silane (**S18**)*

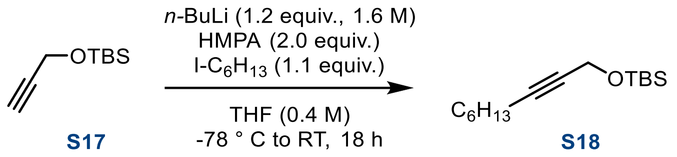

A solution of TBS-protected alkyne **S17** (6.85 g, 40.0 mmol, 1.0 equiv.) in THF (100 mL, 0.4 M) was cooled to −78 °C (dry ice/acetone bath, external temperature), and *n*-BuLi (29.9 mL, 47.8 mmol, 1.2 equiv.; 1.6 M solution in hexanes) was added dropwise. The resulting mixture was stirred at −78 °C for 1 h, after which 1-iodohexane (6.46 mL, 43.8 mmol, 1.1 equiv.) and HMPA (13.9 mL, 79.6 mmol, 2.0 equiv.) were added sequentially. The reaction mixture was allowed to warm to room temperature and stirred for 18 h. Saturated aqueous $NH_4Cl$ (100 mL) was then added, and the layers were separated. The aqueous layer was extracted with EtOAc (3 × 100 mL), and the combined organic layers were dried over $MgSO_4$, filtered, and concentrated under reduced pressure. The crude product was purified by flash column chromatography ($SiO_2$, petroleum ether/EtOAc = 100:1) to afford alkyne **S18** (8.45 g, 83%) as a colorless oil.

**$^1$H NMR (500 MHz, Chloroform-*d*) *δ* (ppm):** 4.30 (t, *J* = 2.2 Hz, 2H), 2.19 (tt, *J* = 7.1, 2.2 Hz, 2H), 1.53 – 1.45 (m, 2H), 1.42 – 1.34 (m, 2H), 1.33 – 1.24 (m, 4H), 0.91 (d, *J* = 0.5 Hz, 9H), 0.88 (t, *J* = 7.0 Hz, 3H), 0.12 (d, *J* = 0.5 Hz, 6H).

**$^{13}$C {$^1$H} NMR (126 MHz, Chloroform-*d*) *δ* (ppm):** 85.7, 78.7, 52.2, 31.5, 28.7, 26.0, 25.9, 22.7, 18.9, 18.5, 14.2, -5.0.

**MS (ESI) *m/z* (%):** 256 [M+H]$^+$.

*Non-2-yn-1-ol (**S19**)*

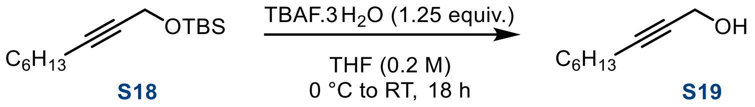

Alkyne **S18** (8.45 g, 32.9 mmol, 1.0 equiv.) was dissolved in THF (164 mL, 0.2 M), and the resulting solution was cooled to 0 °C (ice–water bath, external temperature). Solid TBAF·3$H_2O$ (13.2 g, 41.1 mmol, 1.25 equiv.) was added portionwise, and the resulting mixture was stirred at 0 °C for 3 h. Saturated aqueous $NH_4Cl$ (100 mL) was added, and the mixture was extracted with EtOAc (5 × 50 mL). The combined organic layers were washed with brine (25 mL), dried over $MgSO_4$, filtered, and concentrated under reduced pressure. The crude product was purified by flash column chromatography ($SiO_2$, petroleum ether/EtOAc = 40:1) to afford alcohol **S19** (2.45 g, 54%) as a yellowish oil.

**$^1$H NMR (500 MHz, Chloroform-*d*) *δ* (ppm):** 4.25 (dtd, *J* = 5.9, 2.2, 1.3 Hz, 2H), 2.21 (dddd, *J* = 7.2, 5.1, 2.2, 1.1 Hz, 2H), 1.58 – 1.55 (m, 1H), 1.54 – 1.45 (m, 2H), 1.40 – 1.34 (m, 1H), 1.33 – 1.24 (m, 3H), 0.91 (s, 2H), 0.89 (d, *J* = 7.2 Hz, 3H).

**$^{13}$C {$^1$H} NMR (126 MHz, Chloroform-*d*) *δ* (ppm):** 86.9, 78.4, 51.6, 31.5, 28.7, 25.8, 22.7, 18.9, 14.2.

**MS (ESI) *m/z* (%):** 141 $[M+H]^+$.

*(E)-non-2-en-1-ol (**S20**)*

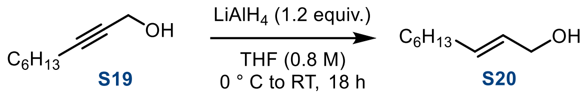

A suspension of $LiAlH_4$ (0.89 g, 23.3 mmol, 1.2 equiv.) in anhydrous THF (24.3 mL, 0.8 M) was cooled to 0 °C (ice–water bath, external temperature), and a solution of alcohol **S19** (2.75 g, 19.4 mmol, 1.0 equiv.) in anhydrous THF (19.4 mL, 1.0 M) was added slowly. The reaction mixture was stirred at 0 °C for 10 min, allowed to warm to room temperature, and stirred for an additional 18 h. The mixture was then cooled to 0 °C, and excess $LiAlH_4$ was carefully quenched by the sequential addition of $H_2O$ (2 mL) and 3.0 M aqueous NaOH (10 mL). Additional $H_2O$ (3 × 15 mL) was added, and the resulting slurry was filtered. The filter cake was washed with $Et_2O$ (5 × 20 mL) and $CH_2Cl_2$ (3 × 50 mL), and the combined filtrates were concentrated under reduced pressure. The crude product was purified by flash column chromatography ($SiO_2$, petroleum ether/EtOAc = 4:1) to afford alcohol **S20** (1.75 g, 63%, *E/Z* >99:1) as a yellowish oil. The *E/Z* ratio was determined by $^1$H NMR analysis.

**$^1$H NMR (500 MHz, Chloroform-*d*) *δ* (ppm):** 5.66 (qt, *J* = 15.4, 6.0 Hz, 2H), 4.08 (d, *J* = 5.5 Hz, 2H), 2.04 (q, *J* = 7.0 Hz, 2H), 1.40 – 1.33 (m, 2H), 1.32 – 1.22 (m, 6H), 0.88 (t, *J* = 6.7 Hz, 3H).

**$^{13}$C {$^1$H} NMR (126 MHz, Chloroform-*d*) *δ* (ppm):** 133.8, 128.9, 64.0, 32.4, 31.9, 29.2, 29.0, 22.8, 14.2.

**MS (ESI) *m/z* (%):** 143 $[M+H]^+$.

**$R_f$** = 0.69 (petroleum ether/EtOAc = 4:1).

*(E)-non-2-enal (**S21**)*

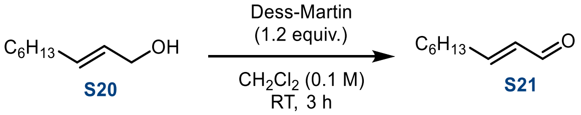

To a stirred solution of alcohol **S20** (1.50 g, 11.7 mmol, 1.0 equiv.) in $CH_2Cl_2$ (117 mL, 0.1 M) was added Dess–Martin periodinane (6.37 g, 14.3 mmol, 1.22 equiv.) at room temperature. The resulting milky suspension was stirred at room temperature for 3 h. Saturated aqueous $NaHCO_3$ (50 mL) was then added, and the mixture was stirred for an additional 25 min, during which time the suspension became clear and two phases formed. The layers were separated, and the aqueous layer was extracted with $CH_2Cl_2$ (3 × 50 mL). The combined organic layers were dried over $MgSO_4$, filtered, and concentrated under reduced pressure. The crude product was purified by flash column chromatography ($SiO_2$, petroleum ether/EtOAc = 4:1) to afford aldehyde **S21** (1.50 g, 91%, E/Z >99:1) as a yellowish oil. The E/Z ratio was determined by $^1$H NMR analysis.

**$^1$H NMR (500 MHz, Chloroform-*d*) *δ* (ppm):** 9.51 (d, *J* = 7.9 Hz, 1H), 6.85 (dt, *J* = 15.6, 6.8 Hz, 1H), 6.12 (ddt, *J* = 15.6, 7.9, 1.6 Hz, 1H), 2.38 – 2.29 (m, 2H), 1.51 (p, *J* = 7.3 Hz, 2H), 1.37 – 1.24 (m, 6H), 0.90 – 0.87 (m, 3H).

**$^{13}$C {$^1$H} NMR (126 MHz, Chloroform-*d*) *δ* (ppm):** 194.4, 159.3, 133.1, 32.9, 31.7, 28.9, 27.9, 22.7, 14.2.

**HRMS (ESI) *m/z:*** $[M+H]^+$ calculated for $C_9H_{16}O$: 141.1274; found: 141.1277.

*Methyl (9E,11Z)-9-nitrooctadeca-9,11-dienoate (**S22**)*

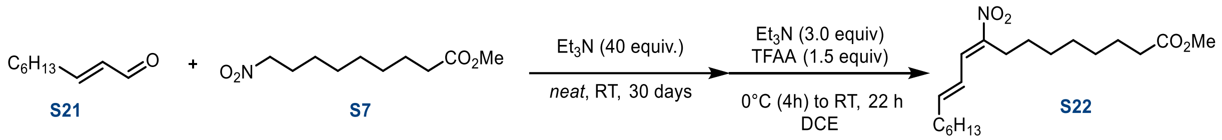

Aldehyde **S21** (0.95 g, 6.70 mmol, 1.7 equiv.) and nitro ester **S7** (0.86 g, 3.94 mmol, 1.0 equiv.) were placed in a reaction flask equipped with a magnetic stir bar, and $Et_3N$ (22 mL, 158 mmol, 40 equiv.) was added. The reaction was monitored until consumption of the starting materials. After 30 days, excess $Et_3N$ was removed under a stream of nitrogen, and the resulting crude product was dissolved in $Et_2O$ (20 mL). The organic layer was sequentially washed with 0.1 M aqueous HCl (2 × 30 mL), $H_2O$ (2 × 30 mL), and brine (30 mL), dried over $Na_2SO_4$, and filtered through a short pad containing silica gel and Celite® (3 g, $SiO_2$/Celite® = 1:1, w/w). The filtrate was concentrated under reduced pressure, and the crude product was purified by gradient flash column chromatography ($SiO_2$, petroleum ether/EtOAc = 30:1 → 1:1) to afford the β-hydroxy adduct (d.r. = 1.1:1, E/Z >99:1), together with recovered aldehyde S21 (0.20 g) and nitro ester **S7** (0.18 g). The isolated β-hydroxy adduct was used directly in the subsequent step. For reagent stoichiometry, the amount of the adduct was calculated based on the initial amount of S7 (3.94 mmol, 1.0 equiv.). Thus, the adduct was dissolved in DCE (16 mL, 0.2 M) and cooled to 0 °C (ice–water bath, external temperature). $Et_3N$ (1.69 mL, 11.82 mmol, 3.0 equiv.) and TFAA (1.01 mL, 5.91 mmol, 1.5 equiv.) were added sequentially. The resulting mixture was stirred at 0 °C for 4 h and then at room temperature for an additional 22 h. The reaction mixture was cooled to 0 °C and quenched by the addition of $H_2O$ (25 mL). The mixture was extracted with EtOAc (3 × 50 mL), and the combined organic layers were dried over $MgSO_4$, filtered, and concentrated under reduced pressure. The crude product was purified by flash column chromatography ($SiO_2$, petroleum ether/EtOAc = 20:1) to afford nitroalkene **S22** (0.54 g, 40%, *(9E,11Z)/(9Z,11Z)* ≥99:1) as a colorless oil. The stereoisomeric ratio was determined by $^1$H NMR analysis.

**$^1$H NMR (500 MHz, Chloroform-*d*) *δ* (ppm):** 7.54 (d, *J* = 11.3 Hz, 1H), 6.34 (dt, *J* = 14.5, 7.0 Hz, 1H), 6.19 (ddt, *J* = 14.8, 11.4, 1.4 Hz, 1H), 3.66 (s, 3H), 2.71 – 2.60 (m, 2H), 2.30 (t, *J* = 7.5 Hz, 2H), 2.25 (q, *J* = 7.8, 7.2 Hz, 2H), 1.62 (dd, *J* = 10.1, 4.5 Hz, 2H), 1.50 (q, *J* = 6.3, 5.4 Hz, 2H), 1.50 – 1.37 (m, 2H), 1.39 – 1.22 (m, 12H), 0.89 (t, *J* = 6.8 Hz, 3H).

**$^{13}$C {$^1$H} NMR (126 MHz, Chloroform-*d*) *δ* (ppm):** 174.4, 149.5, 149.2, 134.1, 123.7, 51.6, 34.2, 33.8, 31.7, 29.1, 29.1, 29.1, 29.0, 28.7, 28.2, 26.7, 25.0, 22.7, 14.2.

**HRMS (ESI) *m/z:*** $[M+H]^+$ calculated for $C_{19}H_{33}NO_4$: 340.2482; found: 340.2480.

*(9E,11Z)-9-nitrooctadeca-9,11-dienoic acid (**9-NO₂cLA**)*

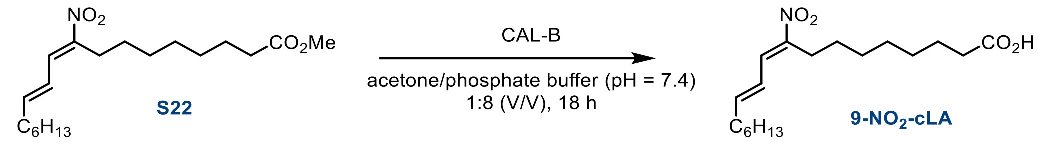

CAL-B (600 mg) was added to a solution of **S22** (1.086 g, 3.20 mmol, 1.0 equiv.) in acetone (21 mL, 0.15 M) and aqueous phosphate buffer (171 mL, 0.01 M, pH 7.4). The resulting mixture was stirred vigorously (1000 rpm) at room temperature for 18 h. The pH was then adjusted to 3 using 1.0 M aqueous HCl, and the mixture was extracted with EtOAc (5 × 75 mL). The combined organic layers were dried over $MgSO_4$, filtered, and concentrated under reduced pressure. The crude product was purified by semipreparative reversed-phase HPLC (C18 column, MeOH/$H_2O$) to afford **9-NO₂cLA** (0.44 g, 42%, *(9E,11Z)/(9Z,11Z)* ≥99:1). The stereoisomeric ratio was determined by $^1$H NMR analysis.

**$^1$H NMR (500 MHz, Chloroform-*d*) *δ* (ppm):** 7.56 – 7.52 (m, 1H), 6.35 (dt, *J* = 14.5, 7.0 Hz, 1H), 6.19 (ddt, *J* = 15.0, 11.4, 1.4 Hz, 1H), 2.69 – 2.62 (m, 2H), 2.35 (t, *J* = 7.5 Hz, 2H), 2.27 – 2.22 (m, 2H), 1.63 (p, *J* = 7.9, 7.1 Hz, 2H), 1.52 (d, *J* = 7.3 Hz, 2H), 1.48 – 1.42 (m, 2H), 1.37 – 1.25 (m, 12H), 0.94 – 0.85 (m, 3H).

**$^{13}$C {$^1$H} NMR (126 MHz, Chloroform-*d*) *δ* (ppm):** 178.3, 149.5, 149.2, 134.1, 123.7, 33.83, 33.79, 31.8, 29.12, 29.06, 29.04, 28.99, 28.7, 28.2, 26.7, 24.7, 22.7, 14.2.

**HRMS (ESI) *m/z:*** $[M\text{-}H]^-$ calculated for $C_{18}H_{30}NO_4$: 324.2180; found: 324.2178.

**R*f*** = 0.33 (hexane/EtOAc = 3:1).

#### **10-NO₂SA** synthesis

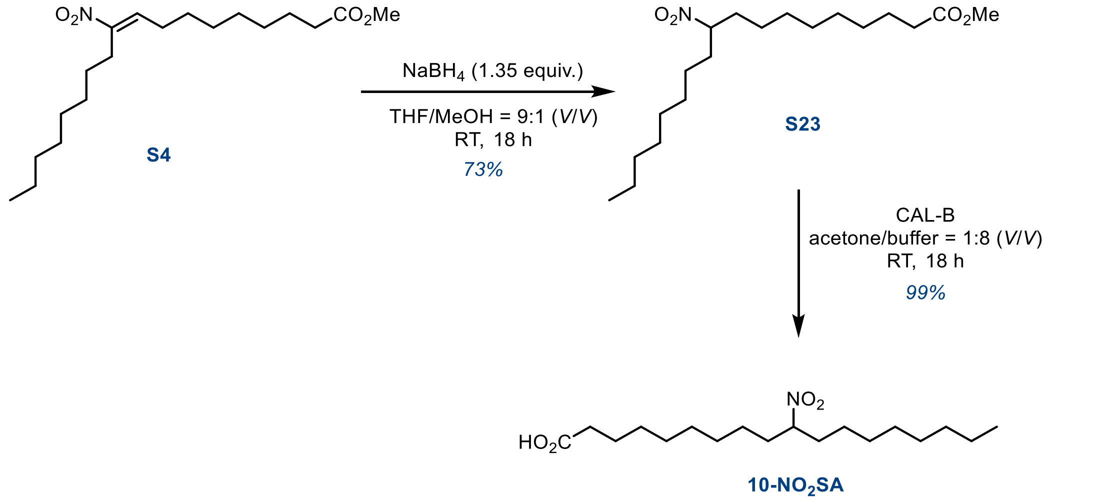

*Methyl 10-nitrooctadecanoate (**S23**)*

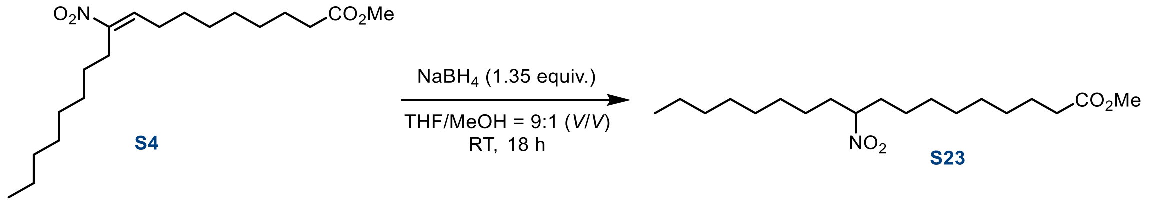

Solid $NaBH_4$ (0.058 g, 1.5 mmol, 1.35 equiv.) was added to a cold solution of **S4** (0.400 g, 1.1 mmol, 1.0 equiv.) in THF/MeOH (5.5 mL, 9:1, v/v) at 0 °C (ice-water bath, external temperature). The cooling bath was removed, and the resulting mixture was stirred at room temperature for 14 h. $H_2O$ (5 mL) and EtOAc (5 mL) were added, and the layers were separated. The aqueous layer was extracted with EtOAc (5 × 20 mL), and the combined organic layers were dried over $MgSO_4$, filtered, and concentrated under reduced pressure. The crude product was purified by flash column chromatography ($SiO_2$; petroleum ether/EtOAc, 10:1) to afford the desired nitro compound **S23** (0.276 g, 73%) as a colorless oil.

**$^1$H NMR (500 MHz, Chloroform-*d*) *δ* (ppm):** 4.45 (tt, *J* = 9.3, 4.5 Hz, 1H), 3.66 (s, 3H), 2.30 (t, *J* = 7.5 Hz, 2H), 1.94 (dtd, *J* = 14.0, 8.9, 4.3 Hz, 2H), 1.67 (dp, *J* = 15.2, 5.4, 5.0 Hz, 2H), 1.59 (dt, *J* = 12.4, 6.1 Hz, 2H), 1.34 – 1.20 (m, 22H), 0.87 (t, *J* = 6.9 Hz, 3H).

**$^{13}$C {$^1$H} NMR (126 MHz, Chloroform-*d*) *δ* (ppm):** 174.5, 89.2, 51.6, 34.2, 34.1, 34.0, 31.9, 29.4, 29.3, 29.20, 29.17, 29.11, 29.0, 25.93, 25.91, 25.0, 22.8, 14.2.

**MS (ESI) *m/z* (%):** 345 $[M+H]^+$.

**HRMS (ESI) *m/z:*** $[M+H]^+$ calculated for $C_{19}H_{38}NO_4$: 344.2795; found: 344.2796.

*10-nitrooctadecanoic acid (**10-NO₂SA**)*

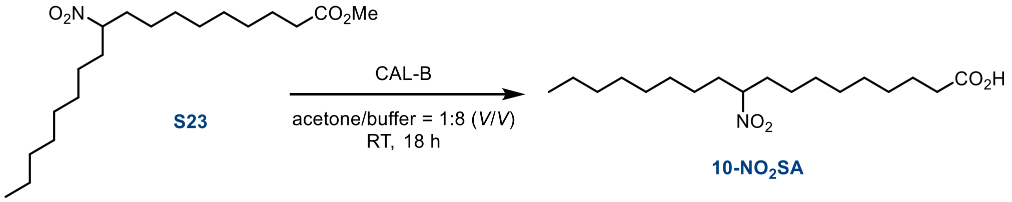

CAL-B (400 mg) was added to a solution of methyl ester **S23** (0.180 g, 0.52 mmol, 1.0 equiv.) in acetone (6.5 mL, 0.08 M) and aqueous phosphate buffer (52.4 mL, 0.01 M, pH 7.4). The resulting mixture was stirred vigorously (1000 rpm) at room temperature for 18 h. The pH was then adjusted to 3 using 1.0 M aqueous HCl, and the mixture was extracted with EtOAc (5 × 20 mL). The combined organic layers were dried over $MgSO_4$, filtered, and concentrated under reduced pressure. The crude product was purified by flash column chromatography ($SiO_2$, petroleum ether/EtOAc = 10:1) to afford **10-$NO_2$SA** (0.170 g, 99%) as a colorless oil.

**$^1$H NMR (500 MHz, Chloroform-*d*) *δ* (ppm):** 11.15 (bs, 1H), 4.45 (tt, *J* = 9.3, 4.5 Hz, 1H), 2.34 (t, *J* = 7.5 Hz, 2H), 1.99 – 1.89 (m, 2H), 1.72 – 1.58 (m, 4H), 1.36 – 1.22 (m, 22H), 0.87 (t, *J* = 7.0 Hz, 3H).

**$^{13}$C {$^1$H} NMR (126 MHz, Chloroform-*d*) *δ* (ppm):** 179.7, 89.2, 34.1, 34.0, 31.9, 29.4, 29.3, 29.19, 29.17, 29.11, 29.06, 29.02, 25.93, 25.90, 24.7, 22.8, 14.2.

**MS (ESI) *m/z* (%):** 329 $[M-H]^-$.

**HRMS (ESI) *m/z:*** $[M-H]^-$ calculated for $C_{18}H_{34}NO_4$: 328.2493; found: 328.2496.

### Literature

(1) Marčeková, M.; Gerža, P.; Šoral, M.; Moncol, J.; Berkeš, D.; Kolarovič, A.; Jakubec, P. Crystallization Does It All: An Alternative Strategy for Stereoselective Aza-Henry Reaction. *Org. Lett.* **2019**, *21* (12), 4580–4584. https://doi.org/10.1021/ACS.ORGLETT.9B01489

(2) Chrenko, D.; Pereckova, J.; Zatloukalová, M.; Vacek, J.; Pospíšil, J.; Perecko, T. Regio- and Stereoselective Synthesis of Nitro-Fatty Acids as NRF2 Pathway Activators Working under Ambient or Hypoxic Conditions. *J. Med. Chem.* **2025**, *68* (11), 12172–12184. https://doi.org/10.1021/acs.jmedchem.5c00982

(3) Reinalda, L.; Heieis, G. A.; Edgerton, L. B.; Bertheussen, K.; Torres-García, D.; Boujadayni, M. el; Steuten, K.; Punt, J. M.; Driever, W. P. F.; Corrigan, C.; Wang, X.; Everts, B.; Finlay, D.; van der Stelt, M.; van Kasteren, S. I. Synthesis of Cyclopropene-Modified Fatty Acids Allows Single-Cell Quantification of Uptake by Immune Cells. *Angew. Chem. Int. Ed.* **2026**, *65* (31), e25040. https://doi.org/10.1002/anie.202525040

(4) Davies, S. G.; Fletcher, A. M.; Foster, E. M.; Lee, J. A.; Roberts, P. M.; Thomson, J. E. Asymmetric Syntheses of APTO and AETD: The β-Amino Acid Fragments within Microsclerodermins C, D, and E. *J. Org. Chem.* **2013**, *78* (6), 2500–2510. https://doi.org/10.1021/jo302731m

### Copy of $^{1}H$ and $^{13}C\{^{1}H\}$ NMR spectra

### Copy of $^{1}$H and $^{13}$C{$^{1}$H} spectra of **S3**

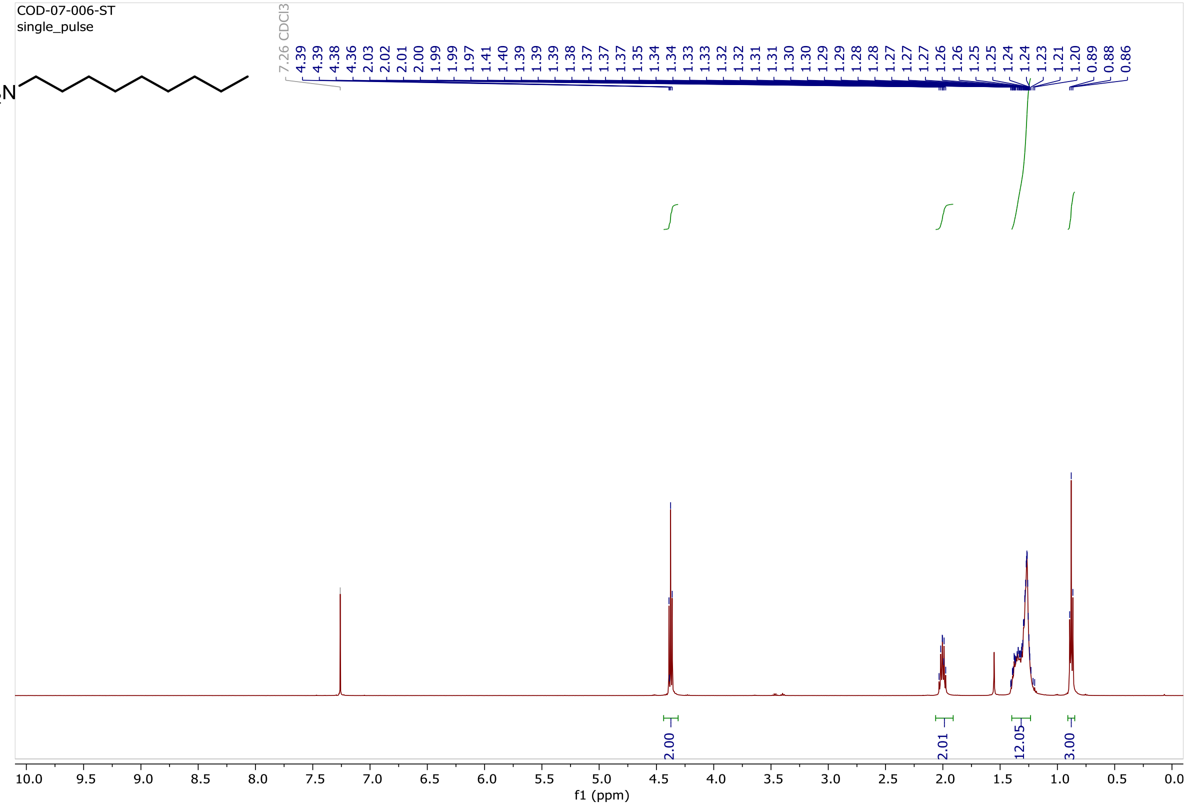

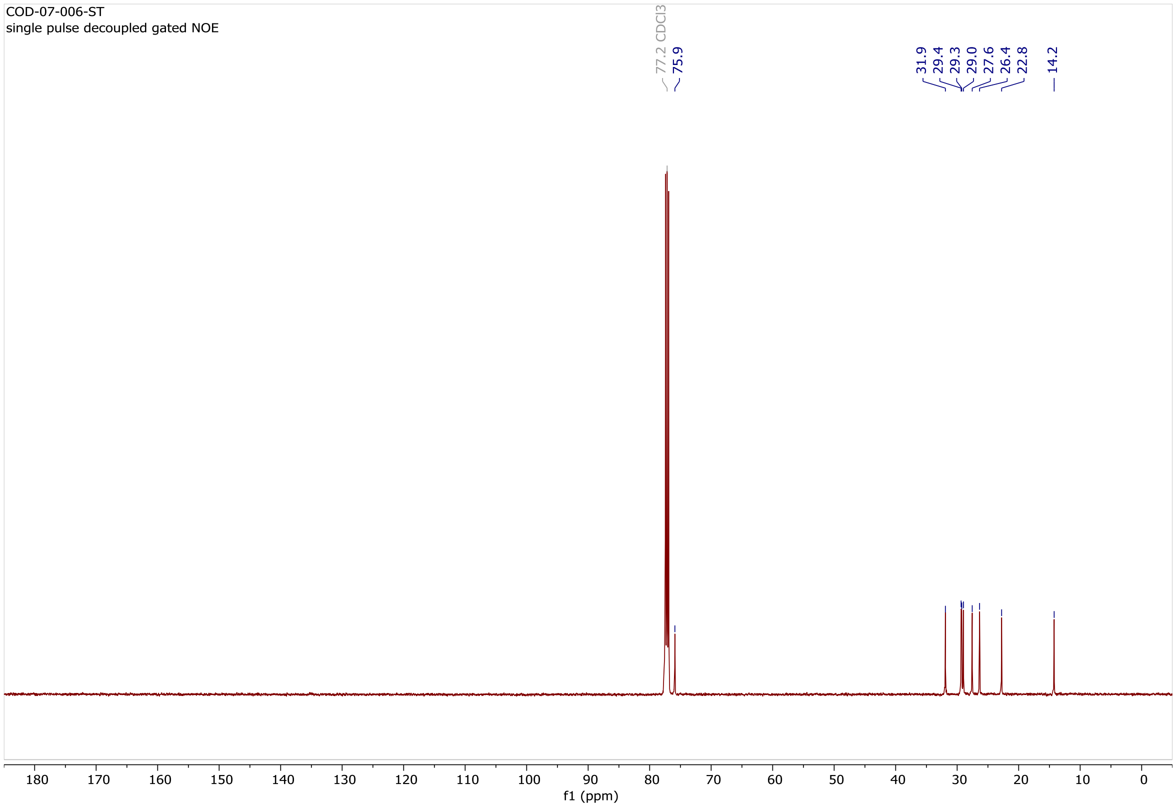

### Copy of $^1$H and $^{13}$C{$^1$H} spectra of **S4**

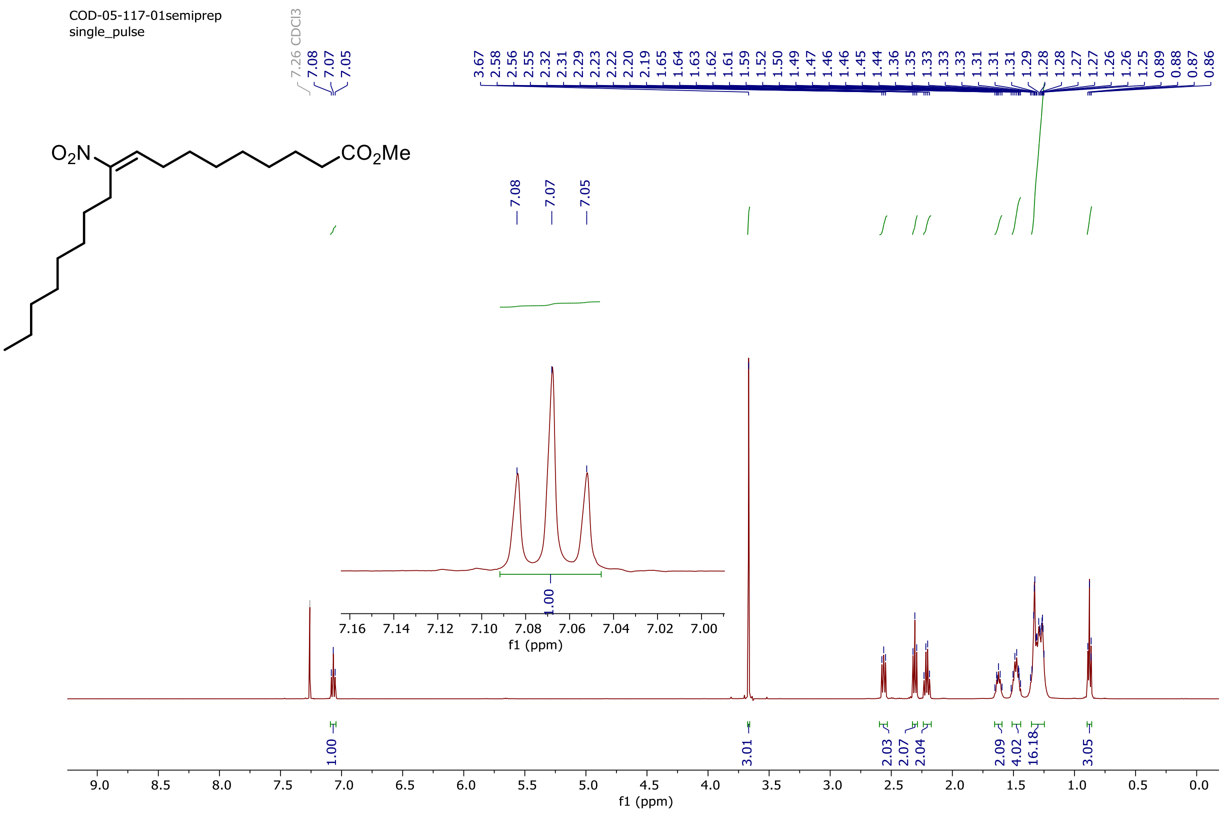

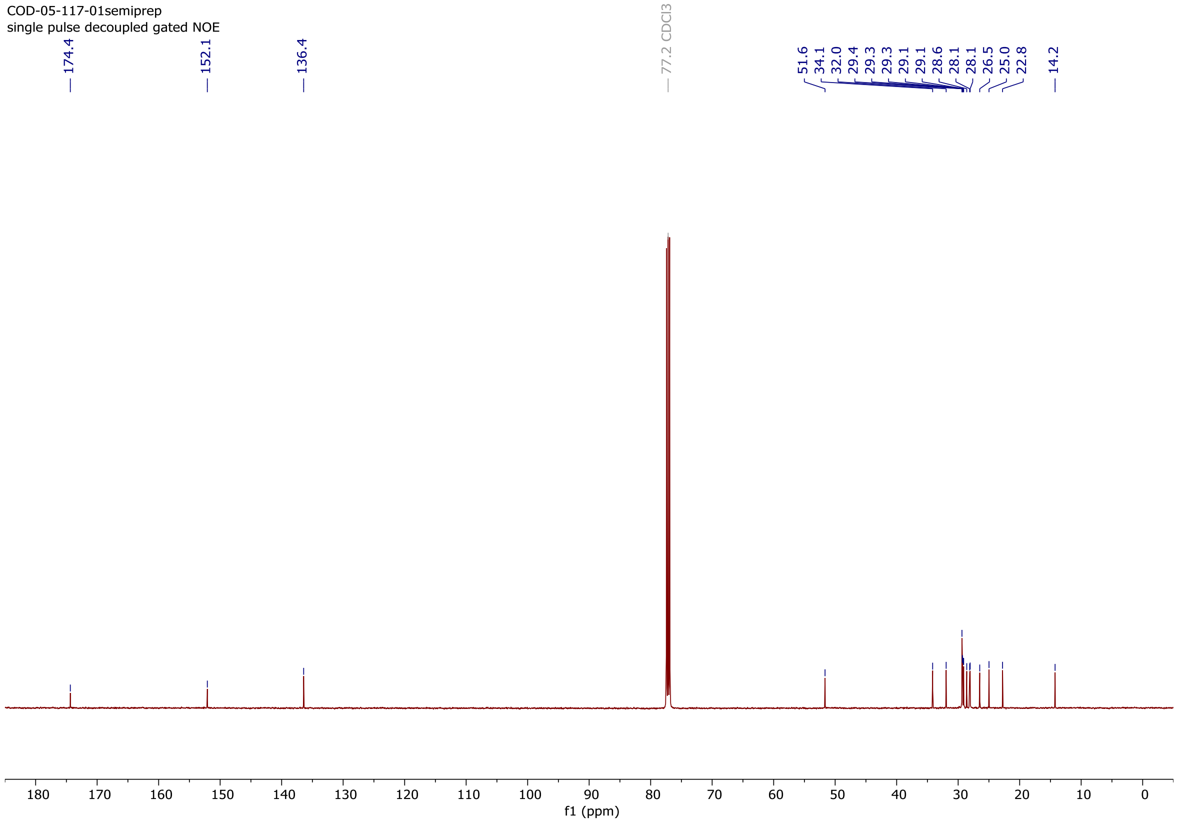

### Copy of $^{1}H$ and $^{13}C\{^{1}H\}$ spectra of **10-NO₂OA**

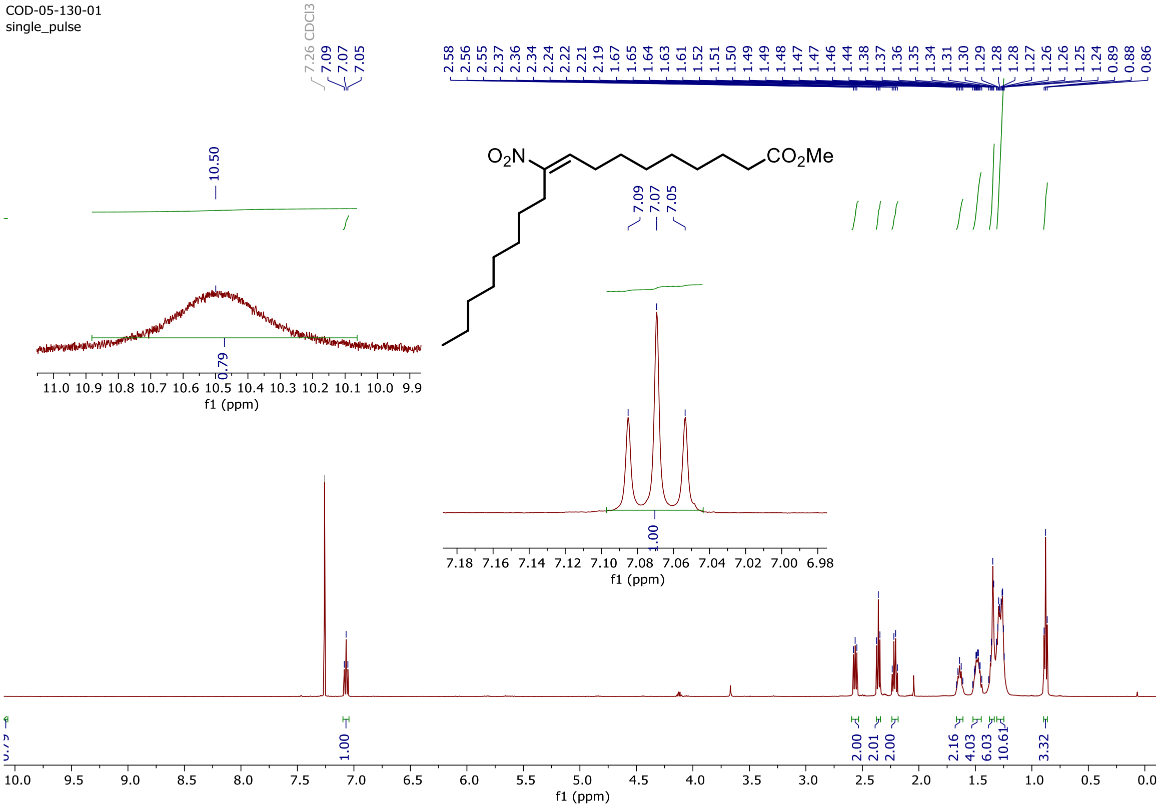

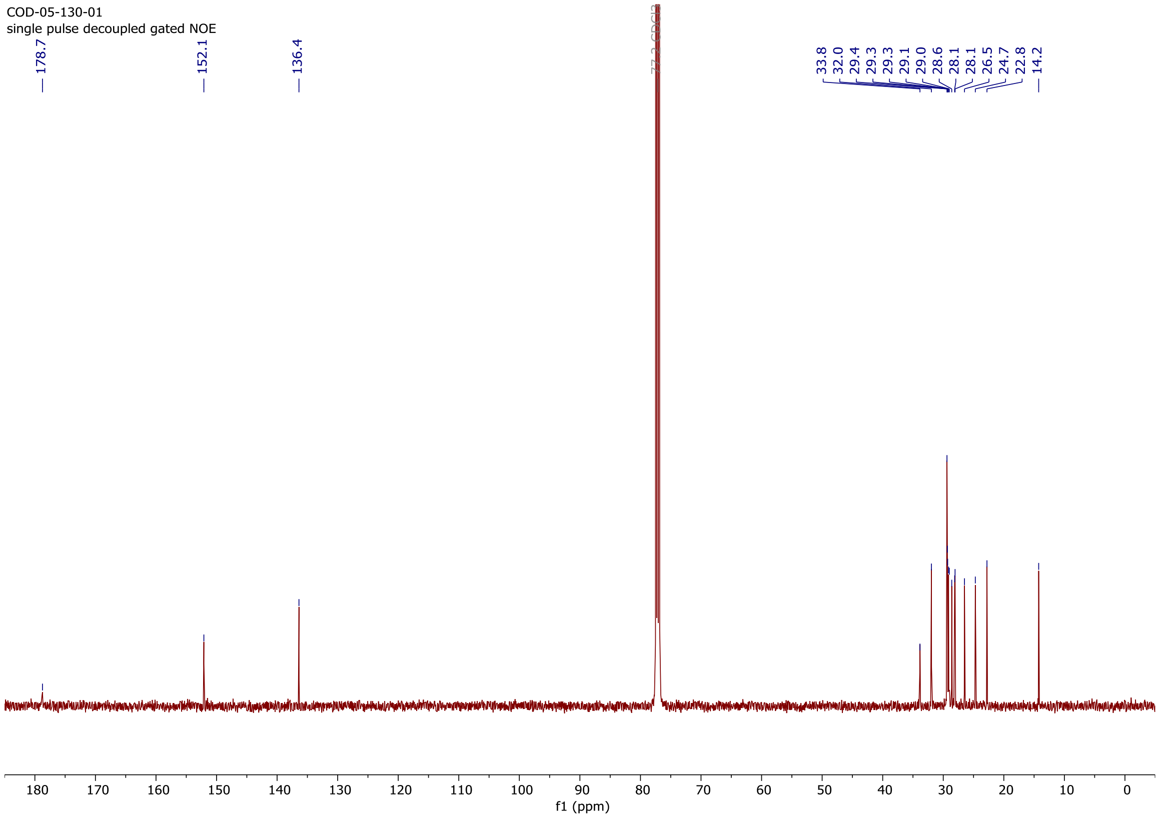

### Copy of $^{1}H$ and $^{13}C\{^{1}H\}$ spectra of **S5**

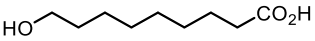

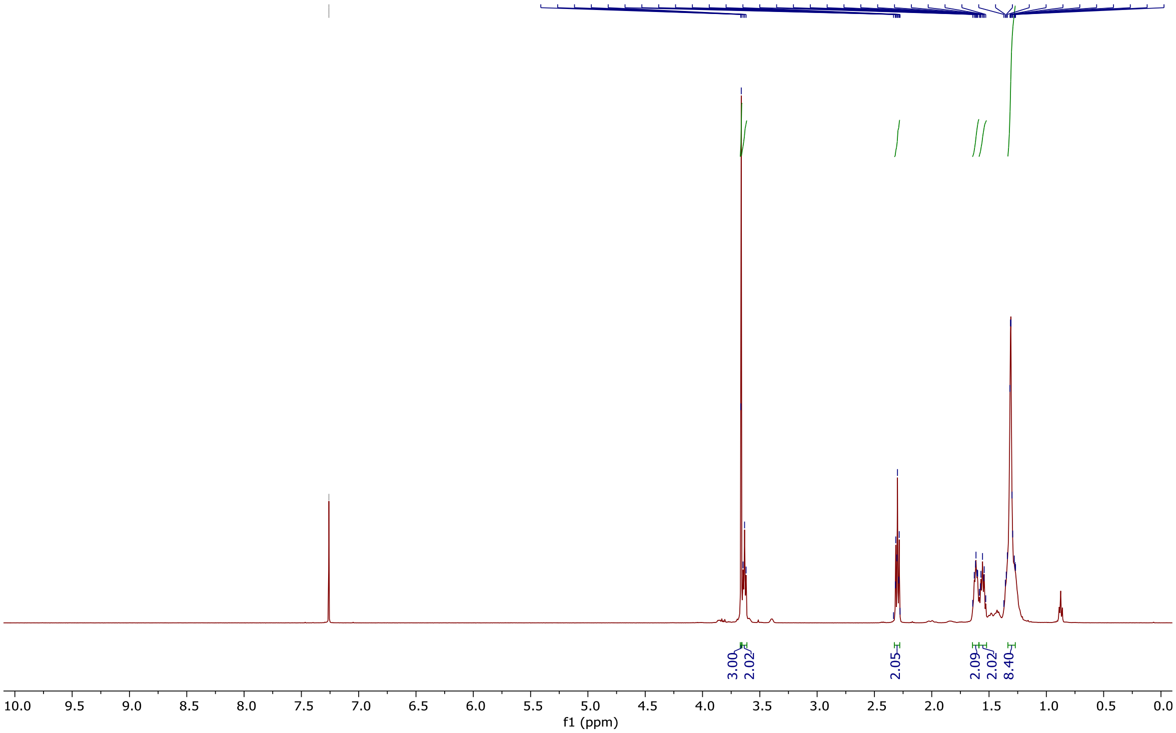

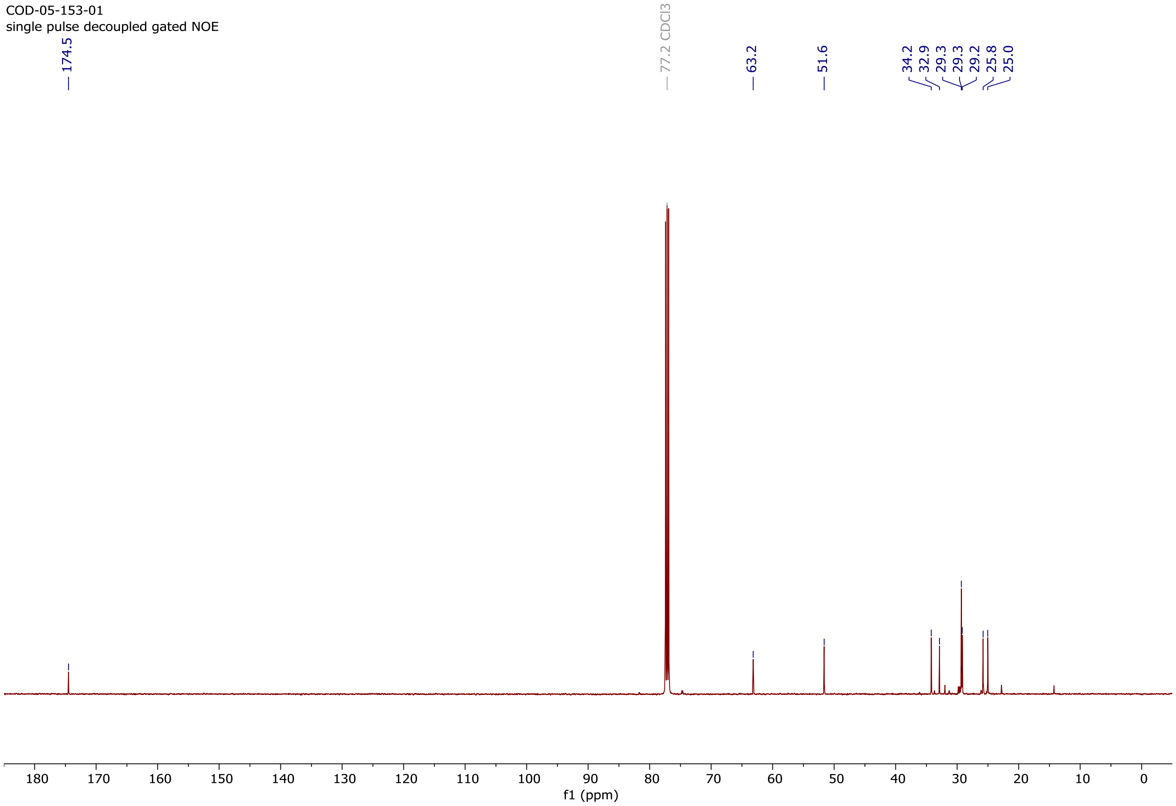

### Copy of $^{1}H$ and $^{13}C\{^{1}H\}$ spectra of **S6**

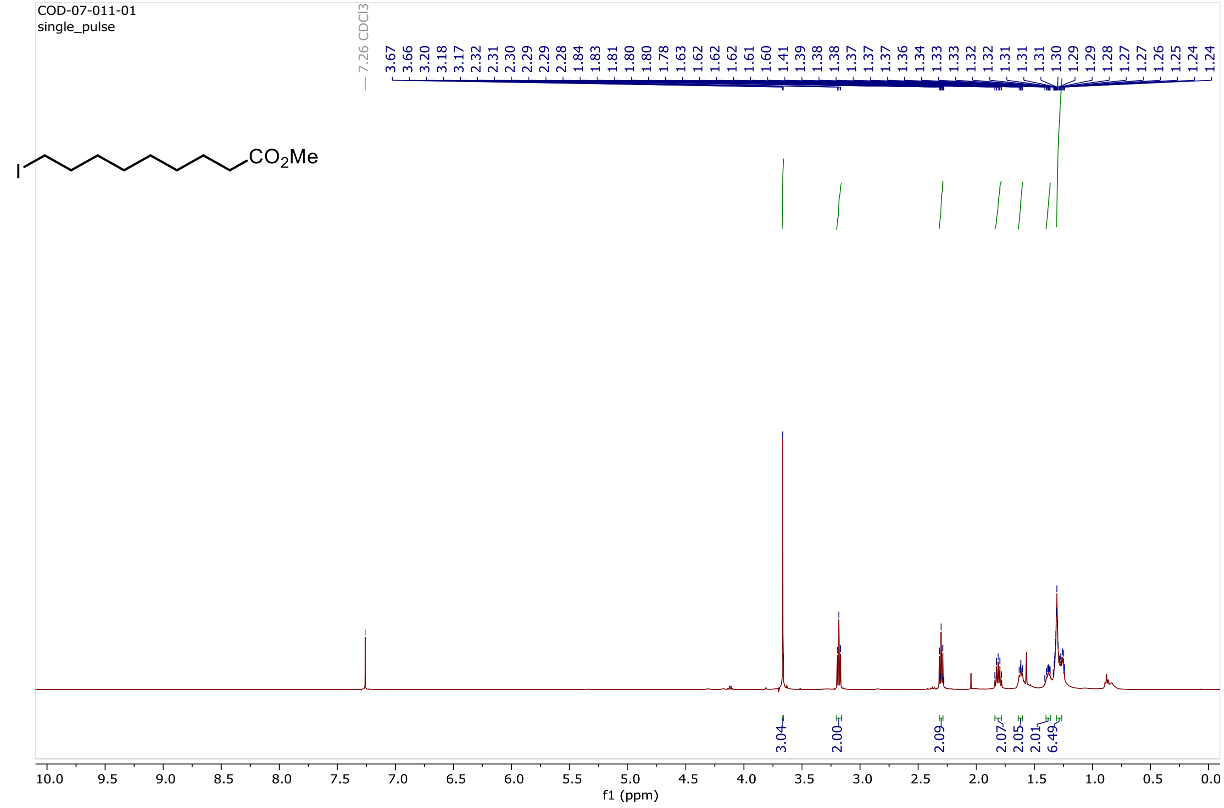

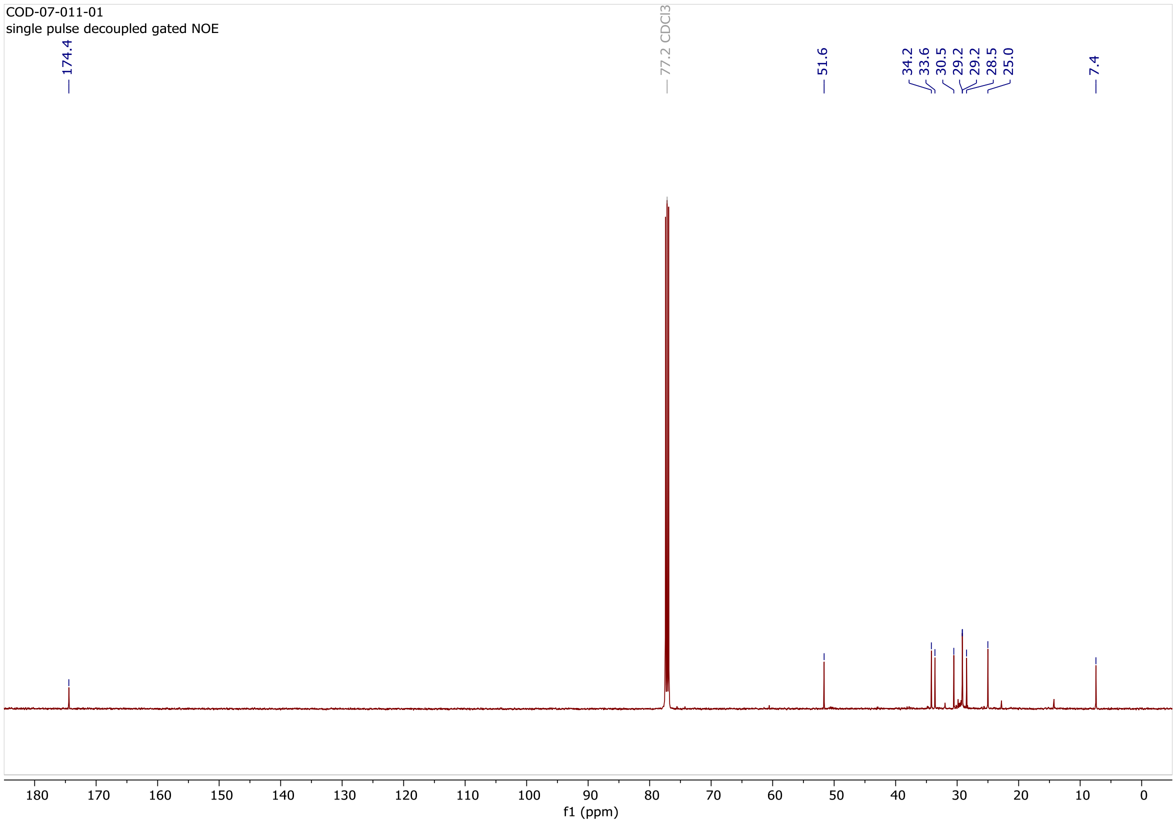

### Copy of $^{1}H$ and $^{13}C\{^{1}H\}$ spectra of **S7**

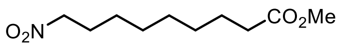

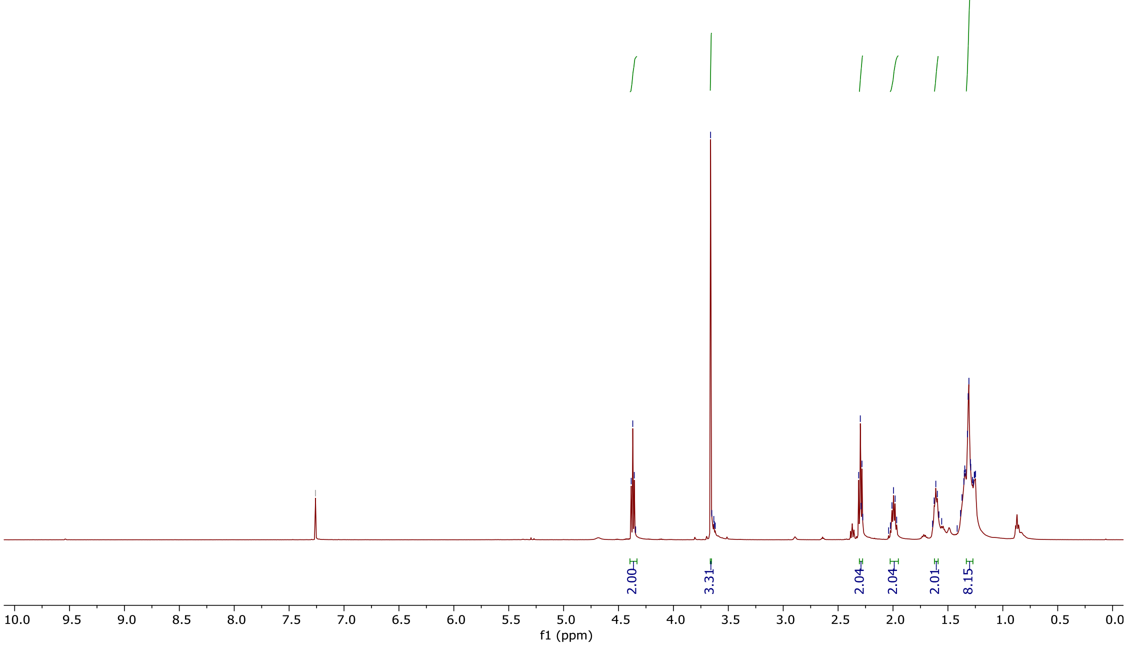

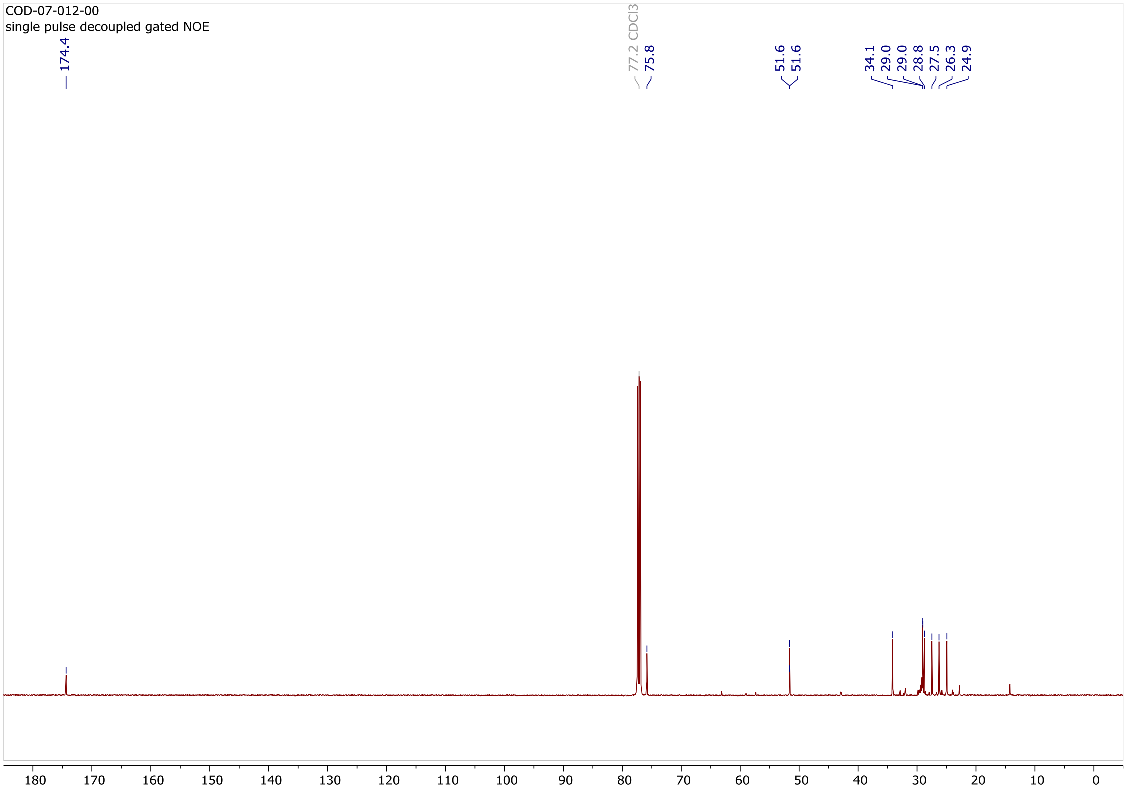

### Copy of $^1H$ and $^{13}C\{^1H\}$ spectra of **S8**

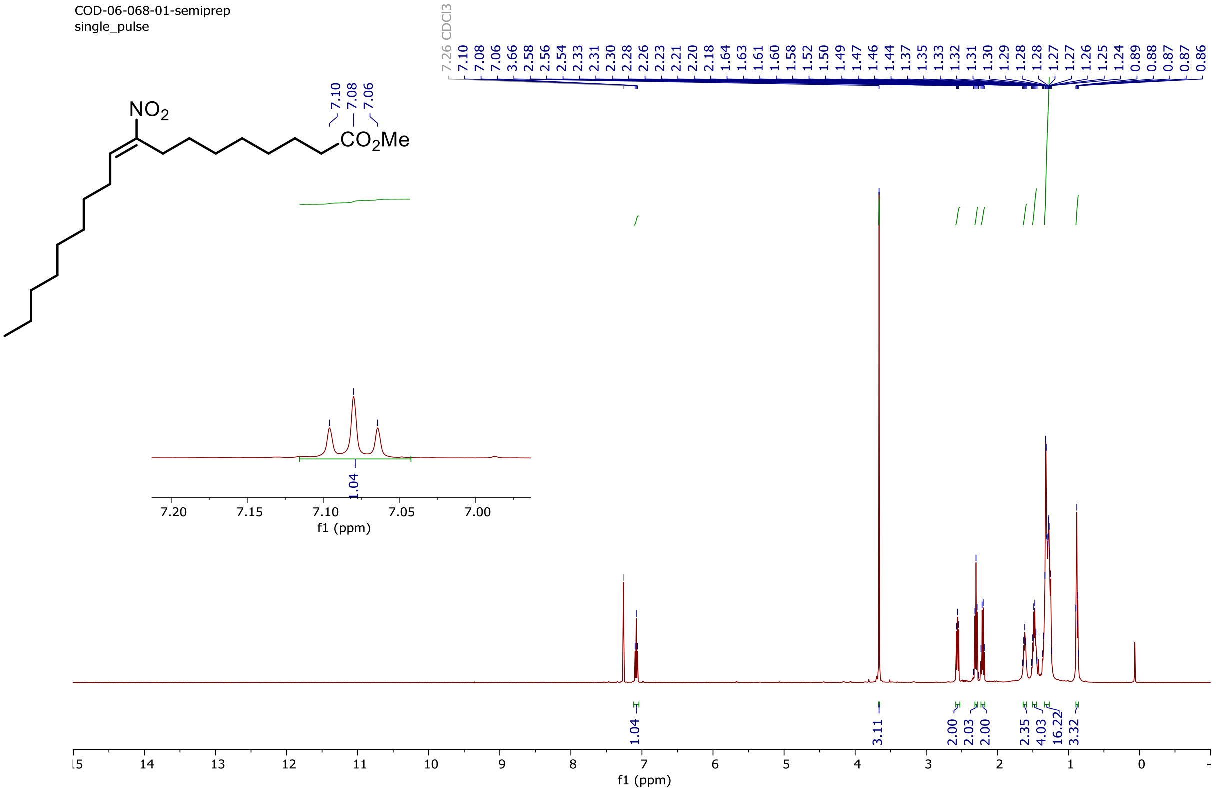

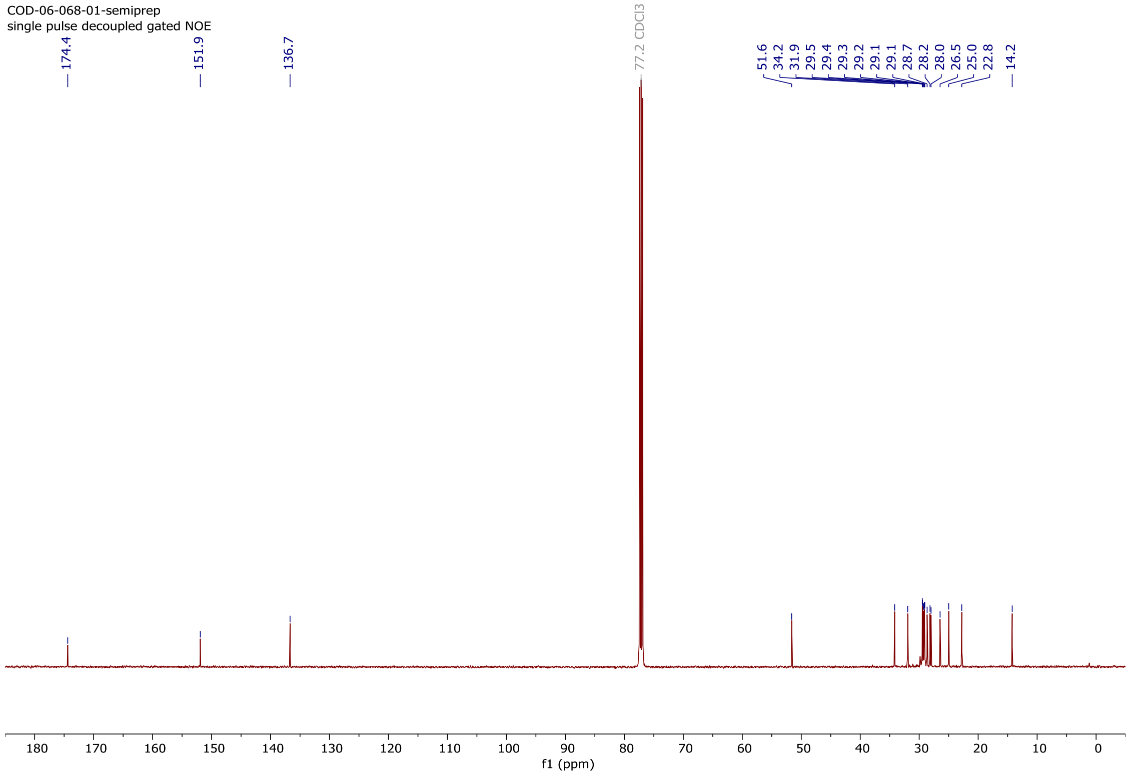

### Copy of [1]H and [13]C{[1]H} spectra of **9-NO₂OA**

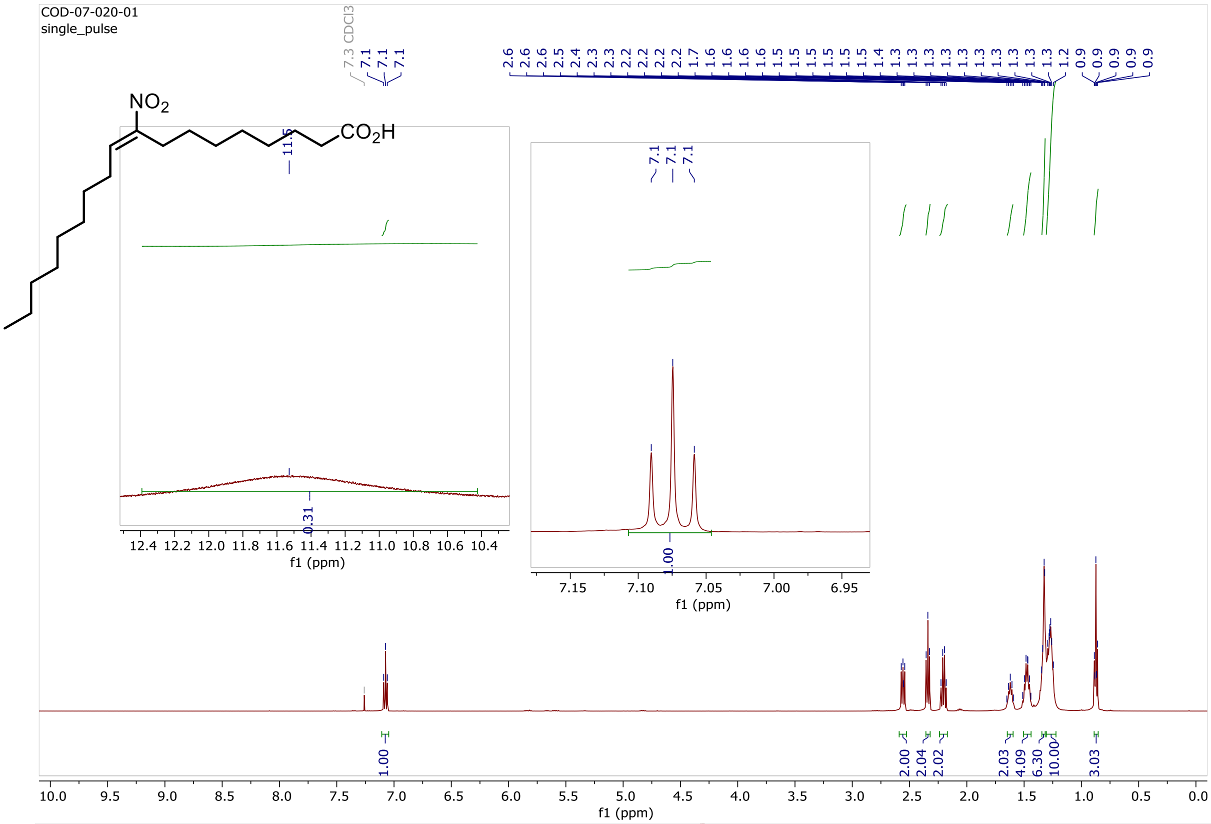

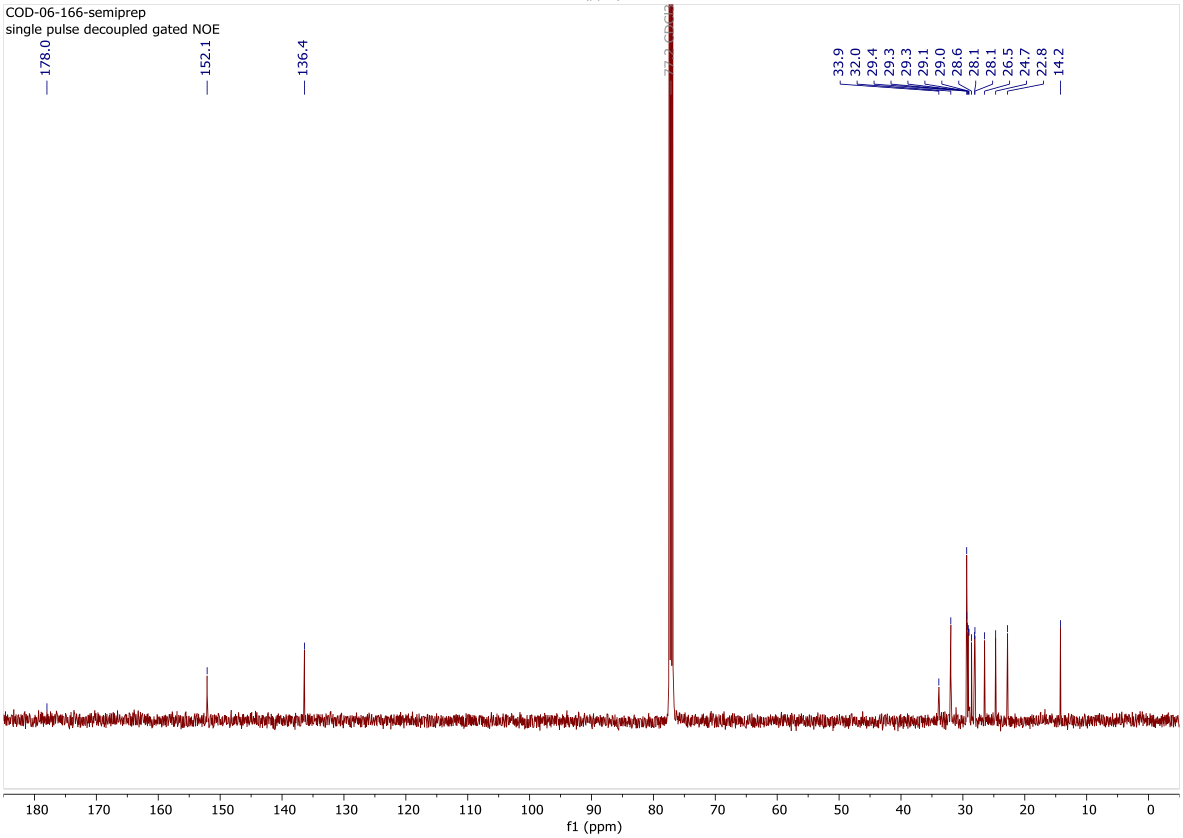

### Copy of $^1$H and $^{13}$C{$^1$H} spectra of **S10**

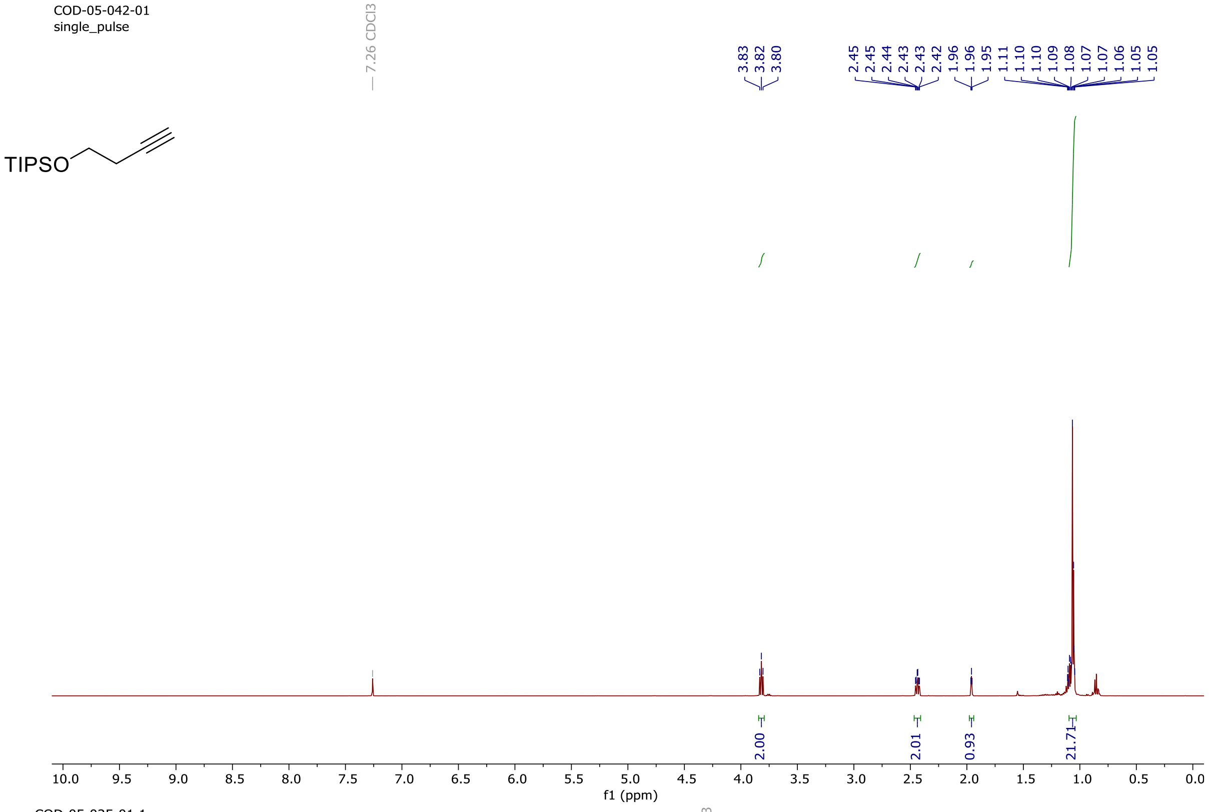

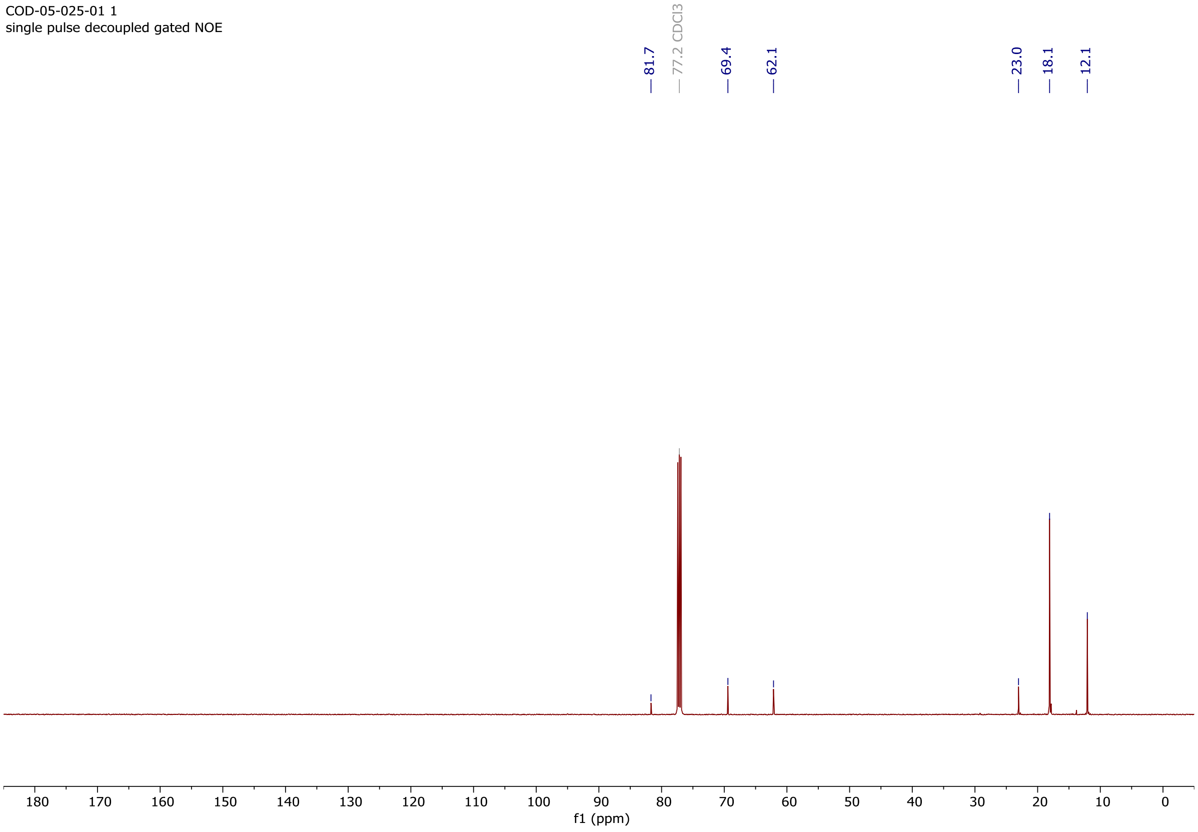

Copy of $^{1}H$ and $^{13}C\{^{1}H\}$ spectra of **S11**

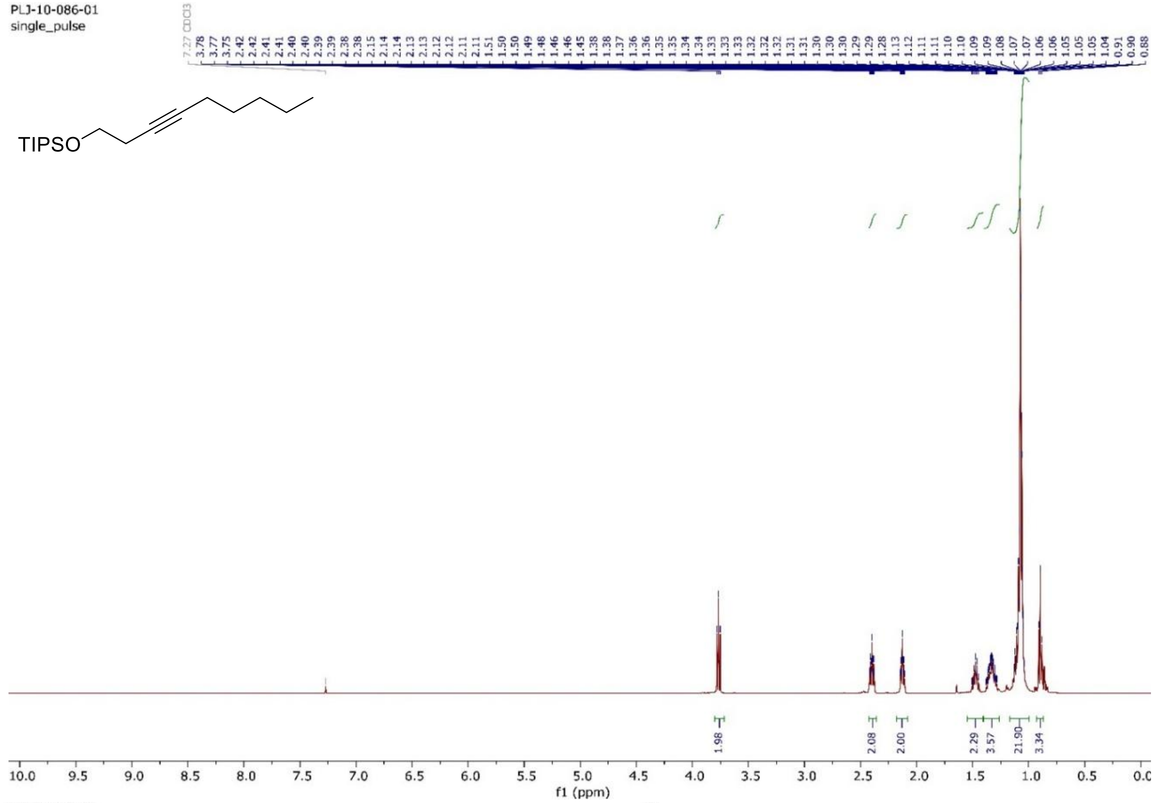

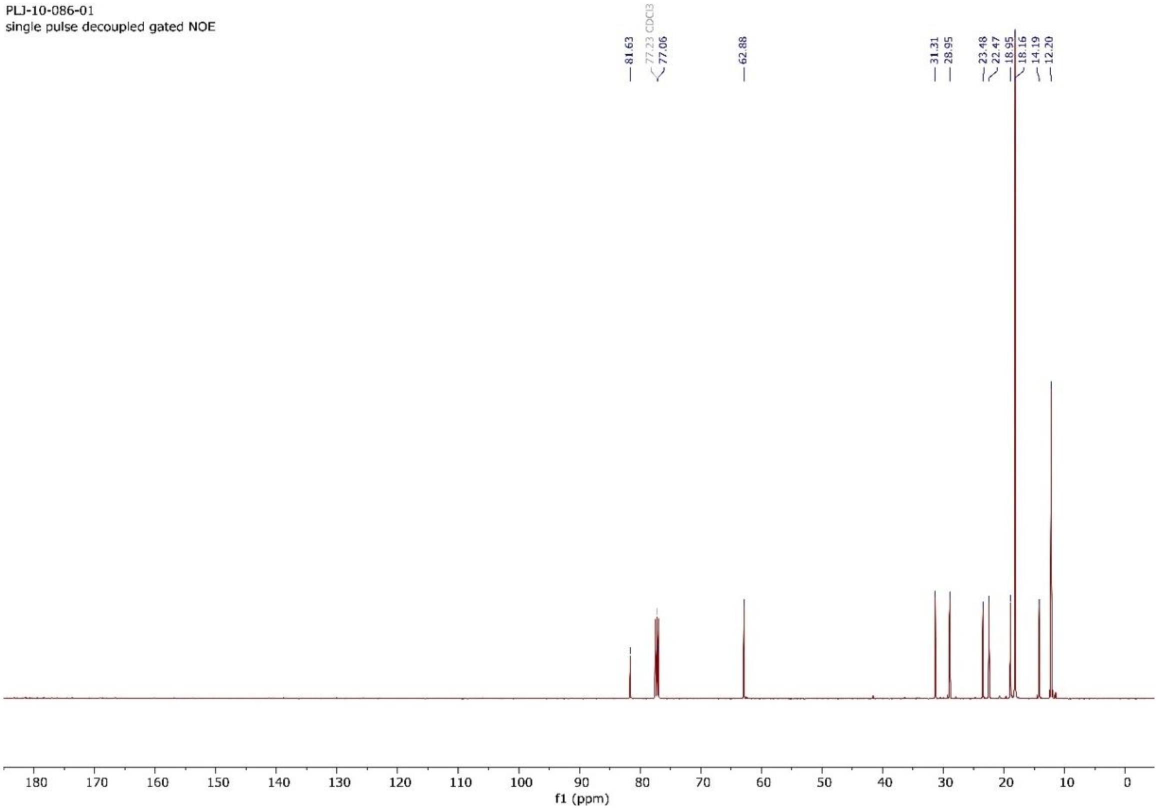

### Copy of $^{1}H$ and $^{13}C\{^{1}H\}$ spectra of **S12**

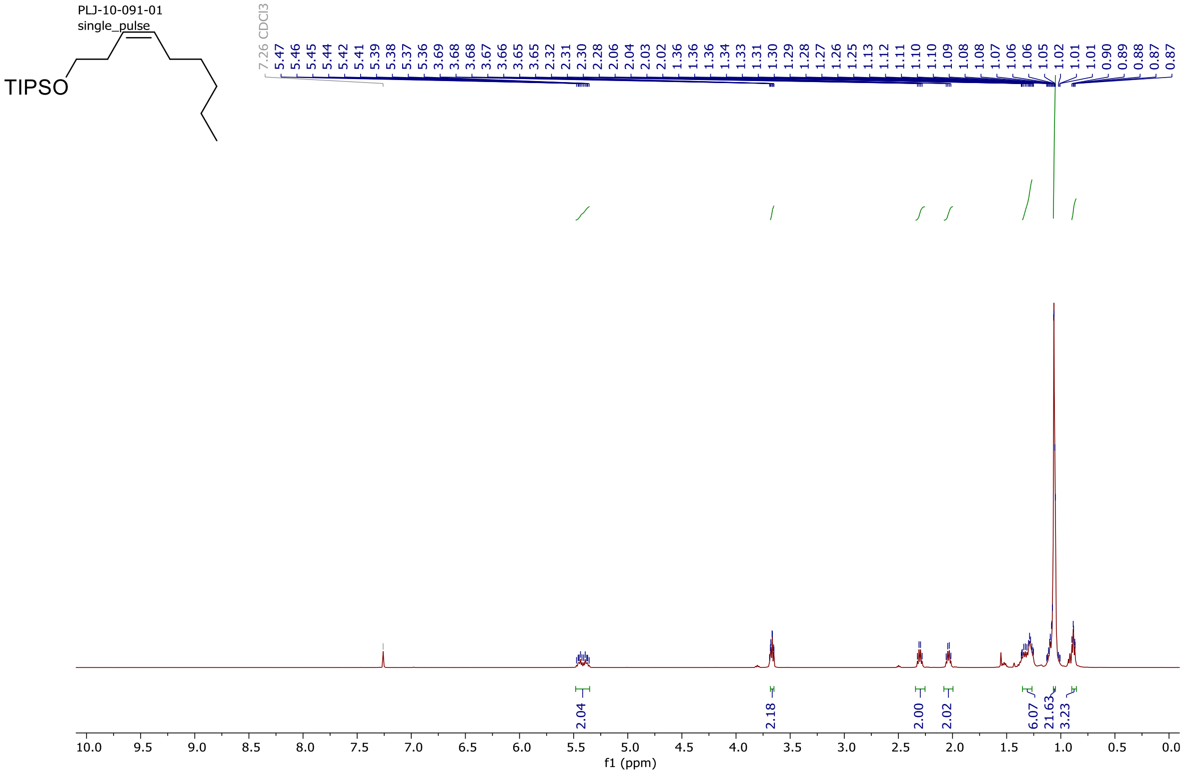

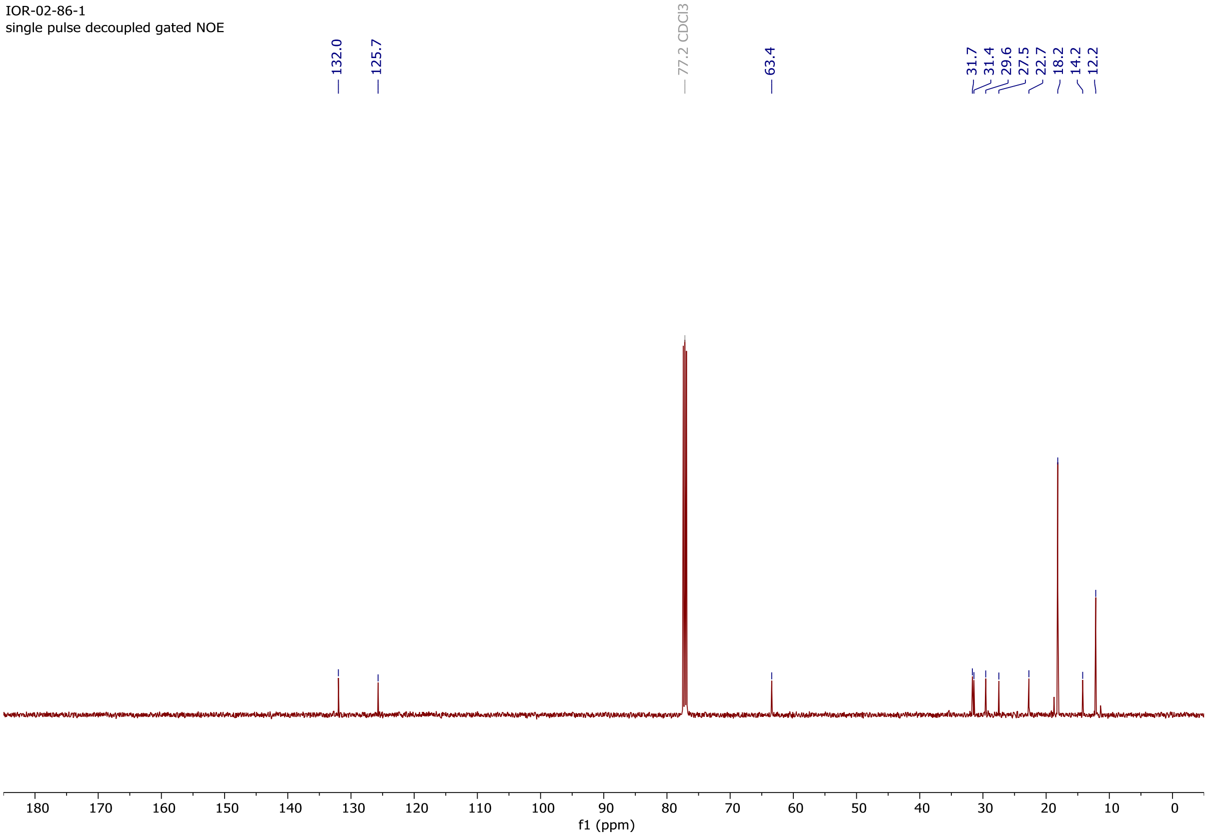

### Copy of $^{1}H$ and $^{13}C\{^{1}H\}$ spectra of **S13**

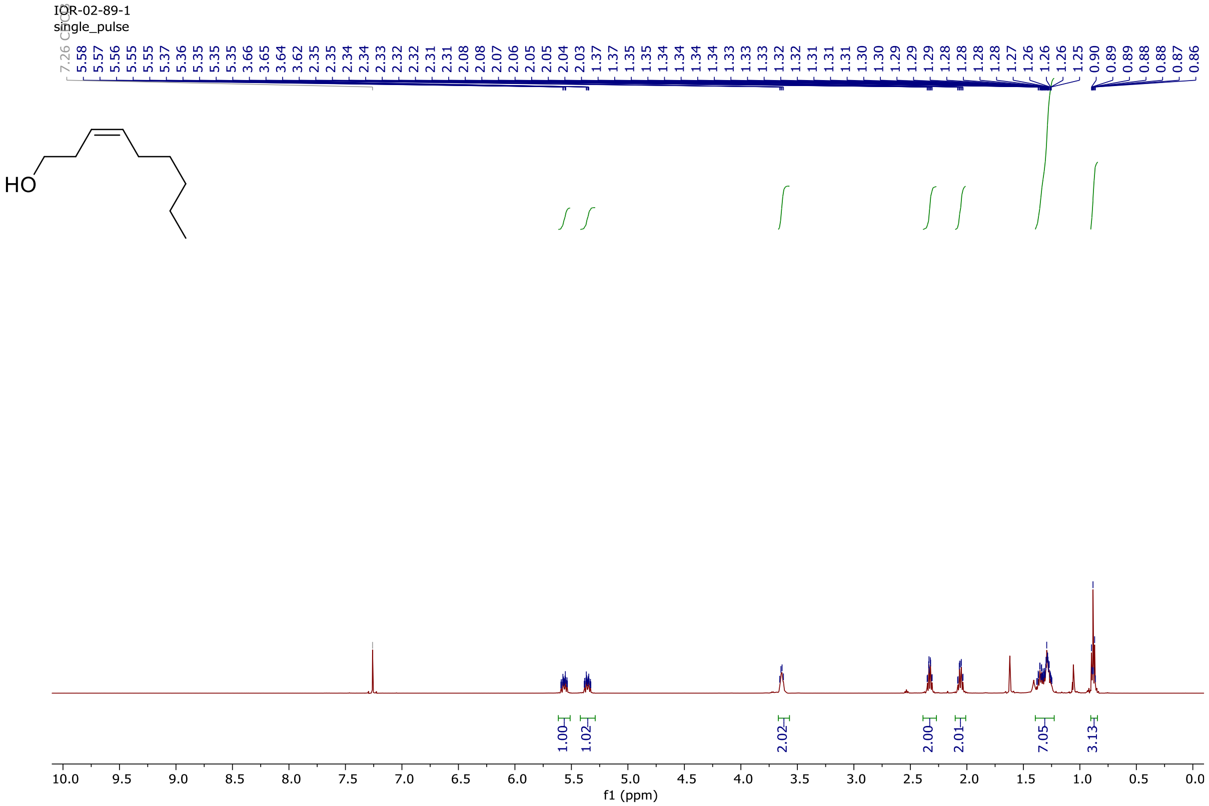

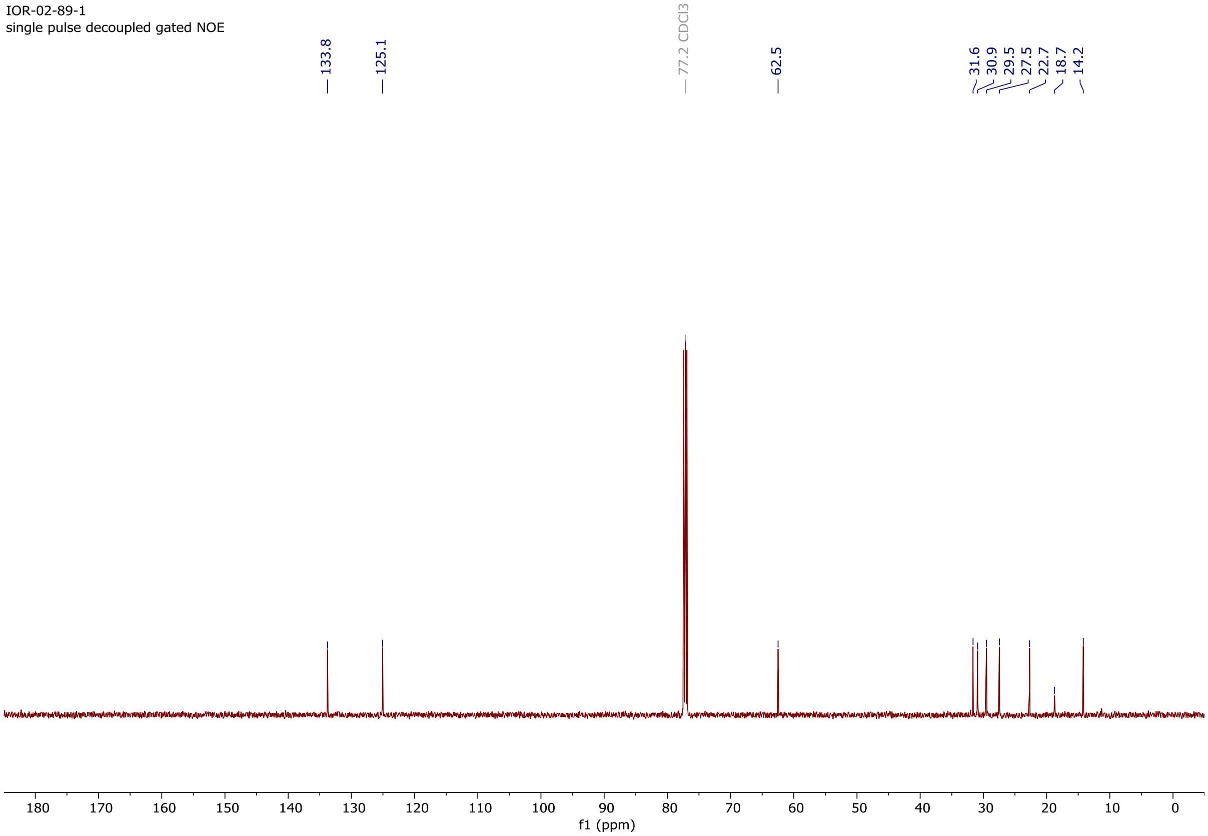

### Copy of $^{1}$H and $^{13}$C{$^{1}$H} spectra of **S14**

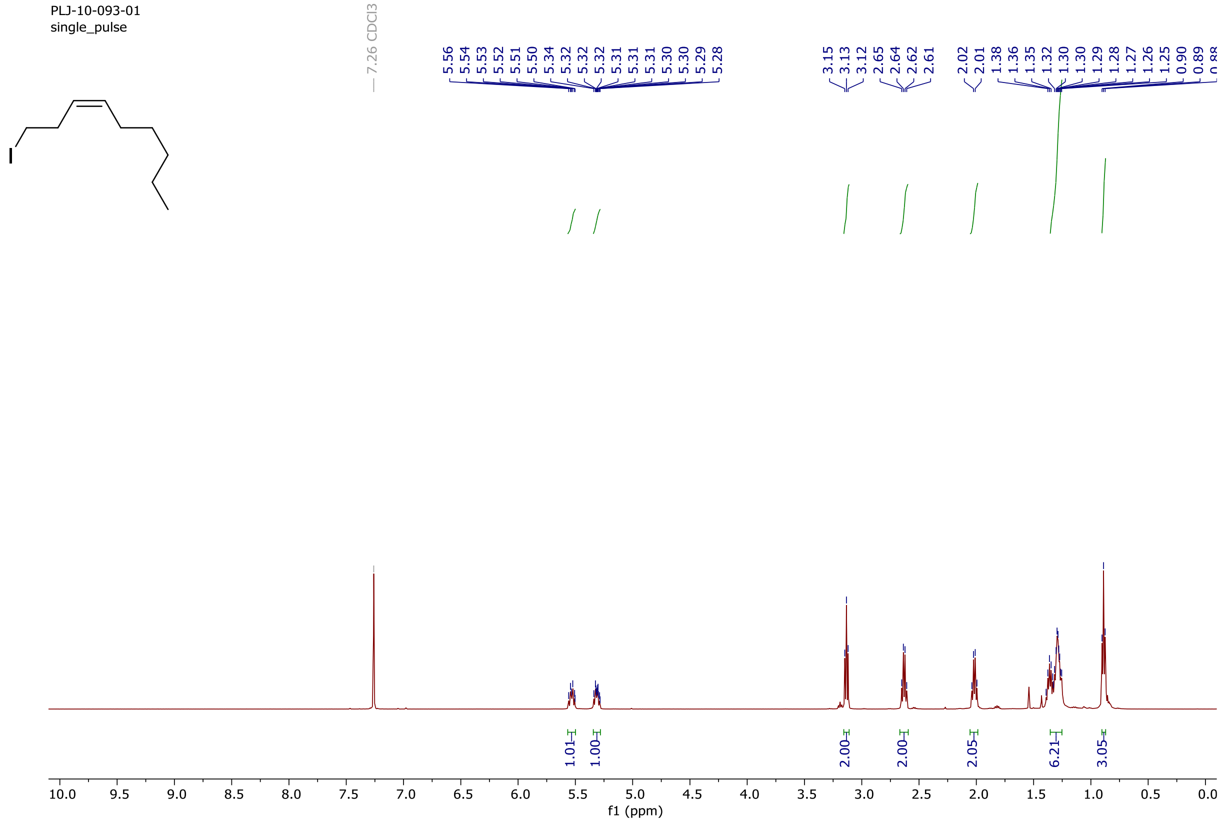

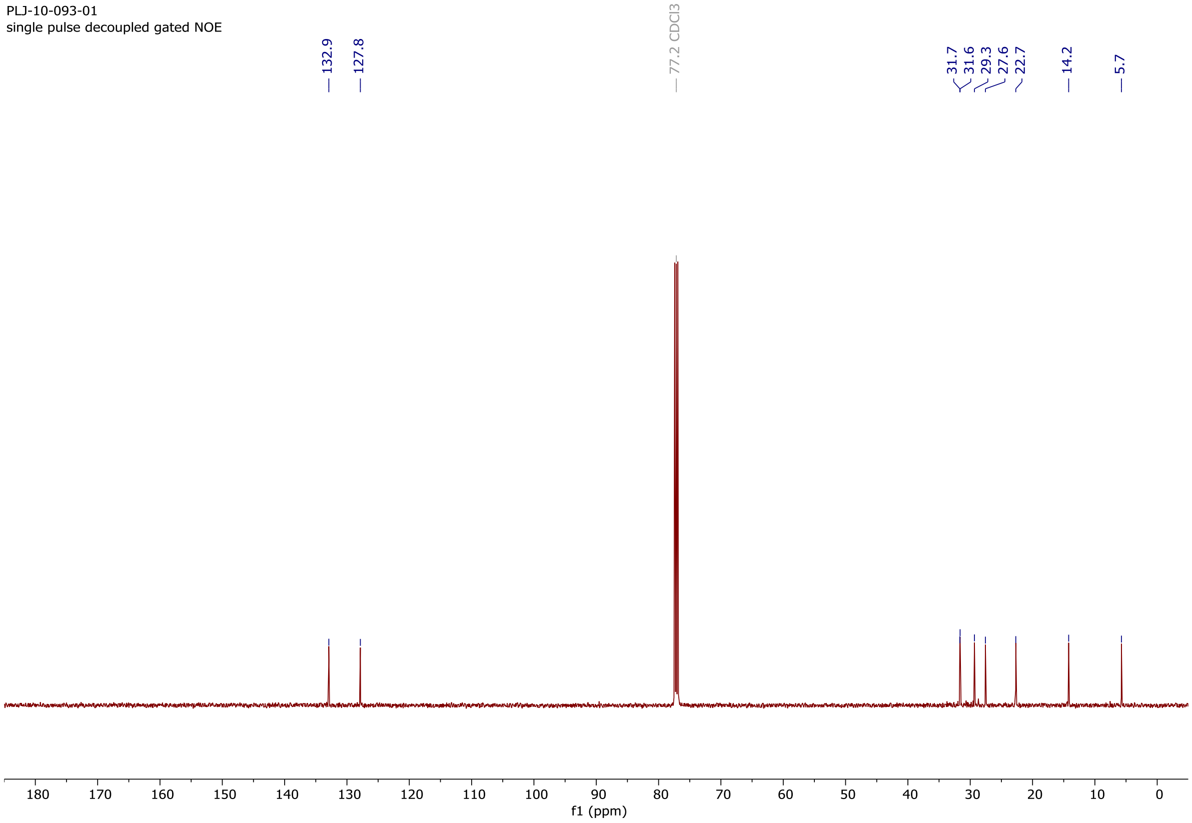

### Copy of $^{1}H$ and $^{13}C\{^{1}H\}$ spectra of **S15**

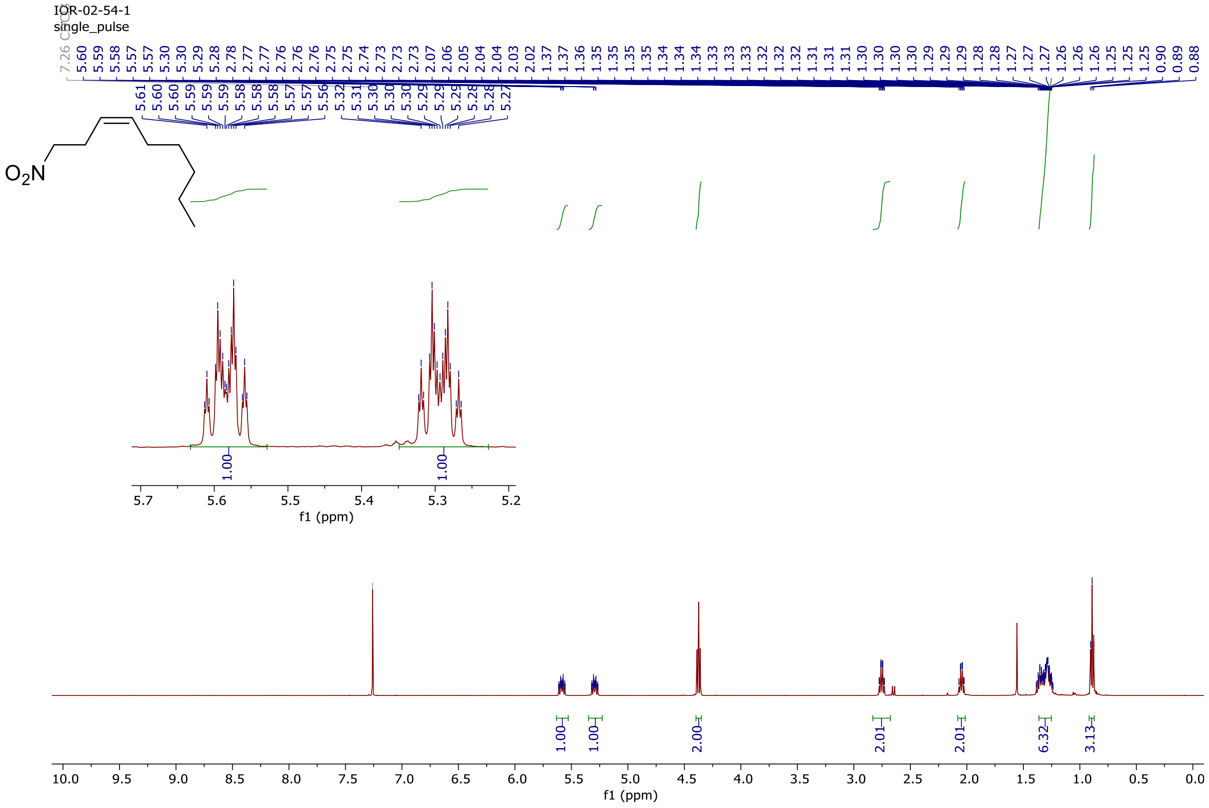

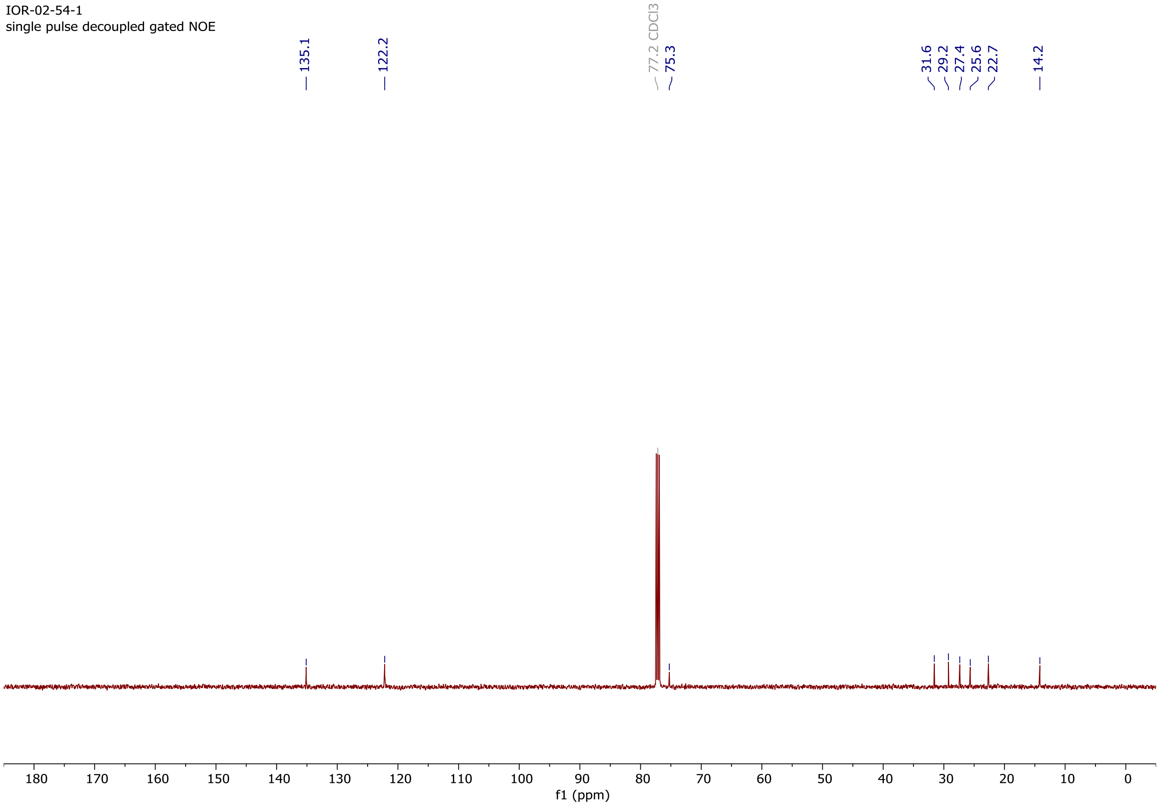

### Copy of $^1$H and $^{13}$C{$^1$H} spectra of **S16**

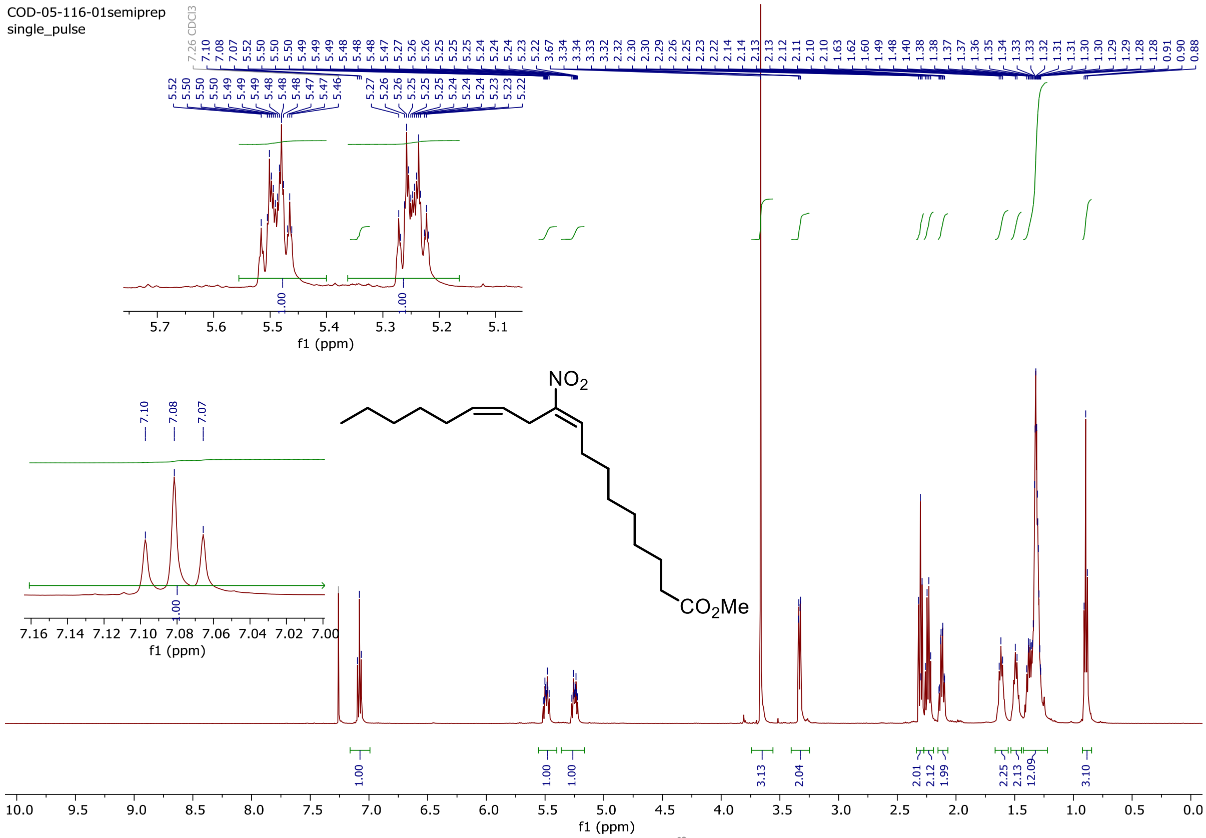

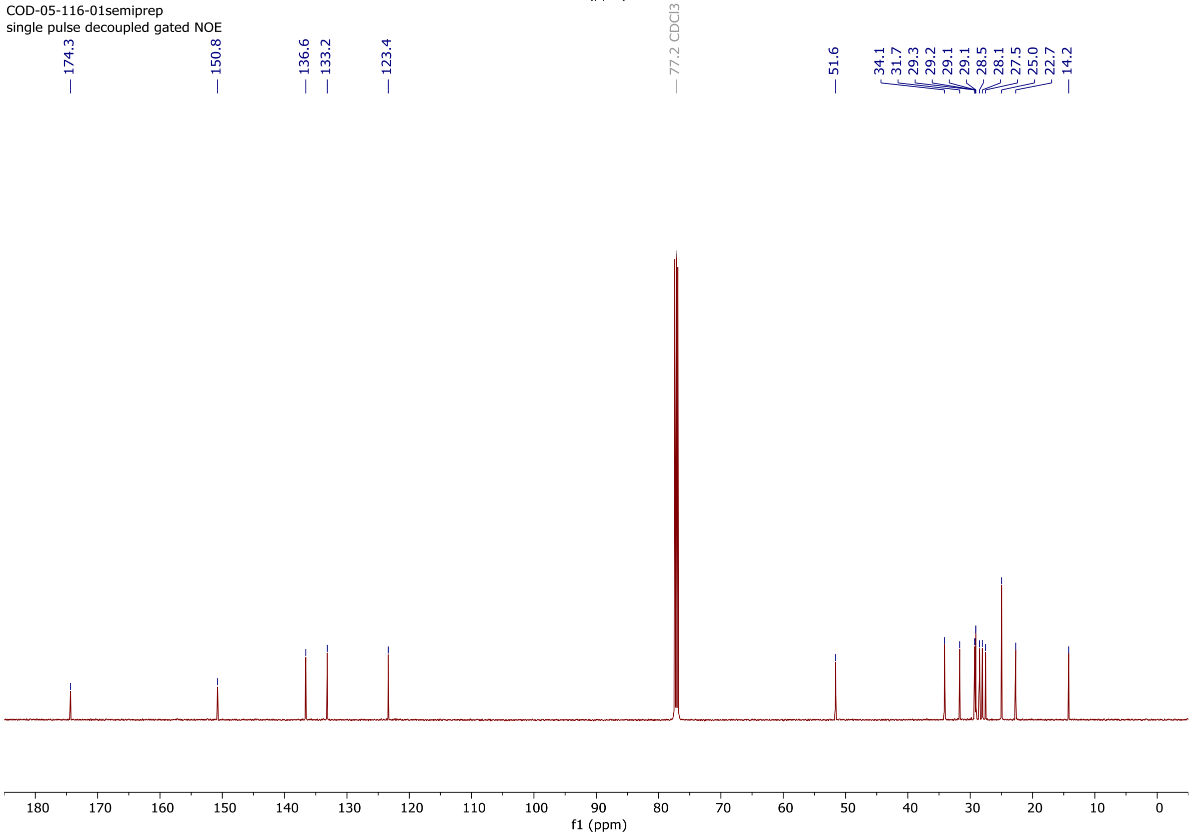

### Copy of $^{1}H$ and $^{13}C\{^{1}H\}$ spectra of **10-$NO_2$LA**

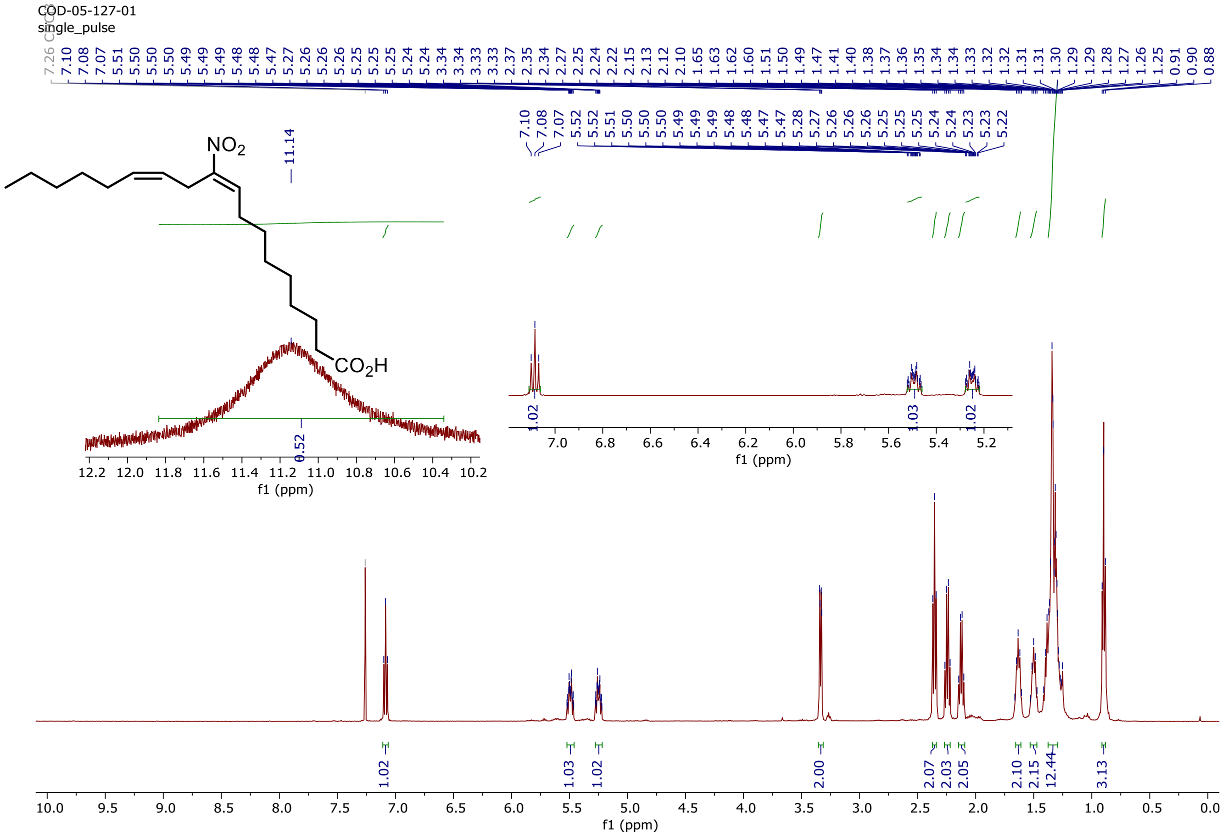

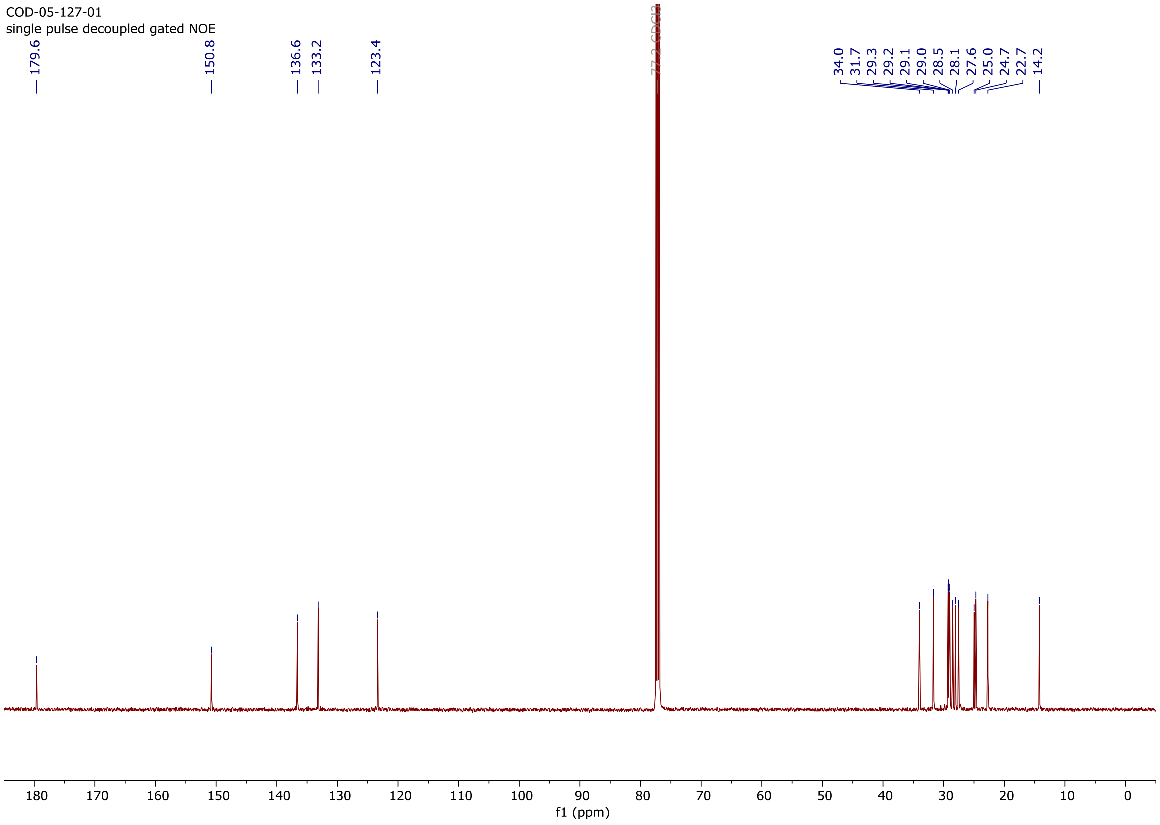

### Copy of $^{1}H$ and $^{13}C\{^{1}H\}$ spectra of **S19**

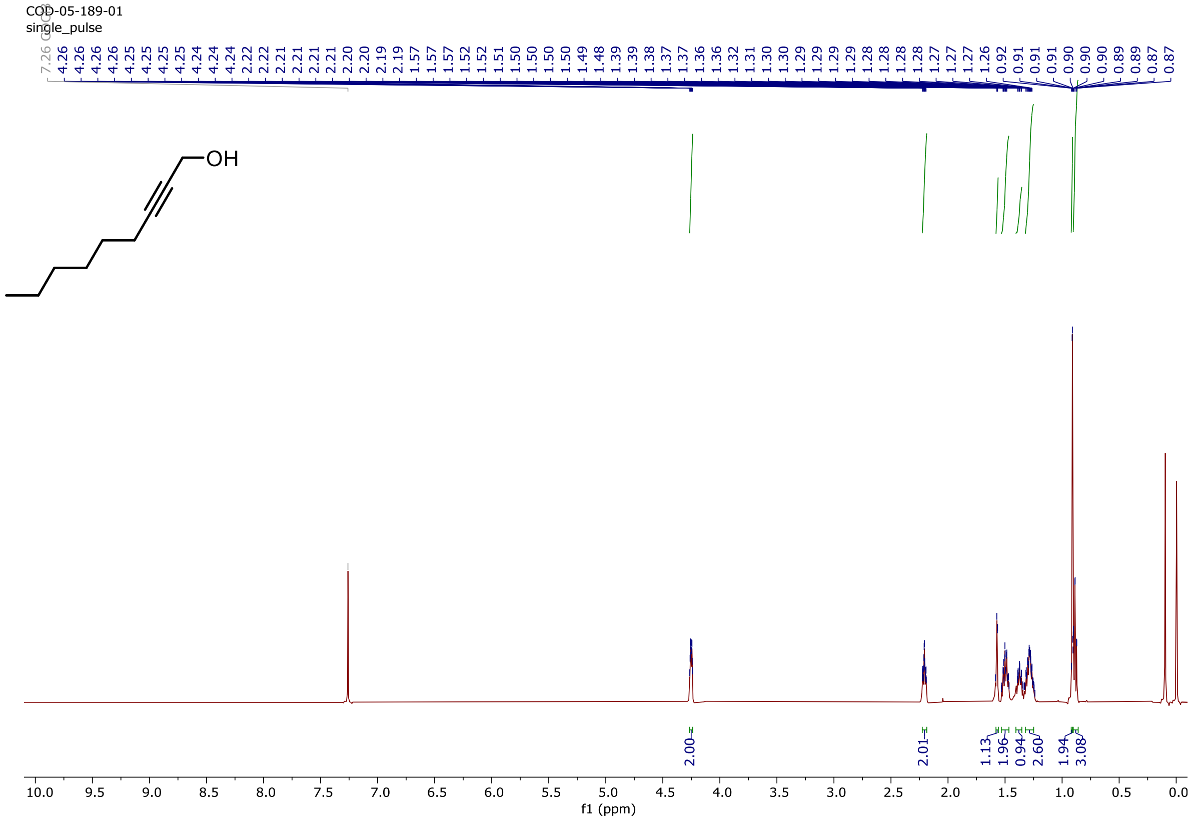

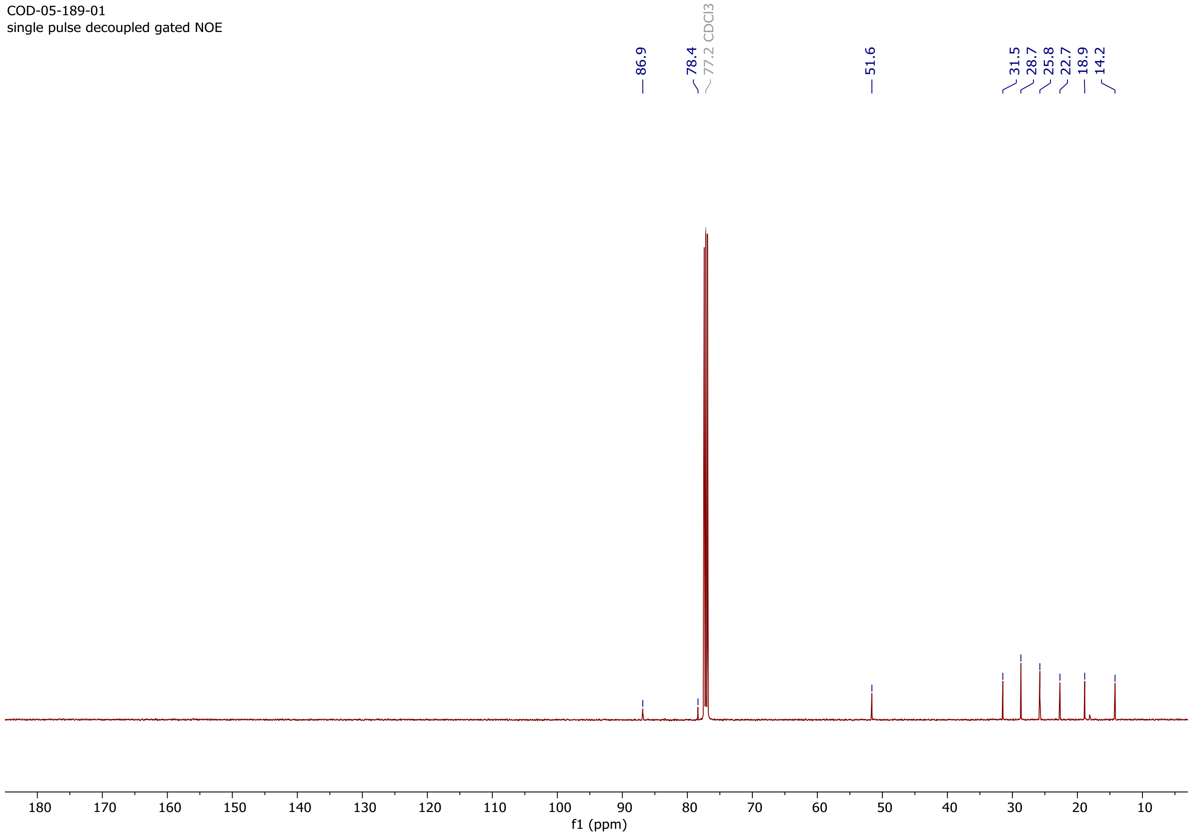

### Copy of $^{1}H$ and $^{13}C\{^{1}H\}$ spectra of **S20**

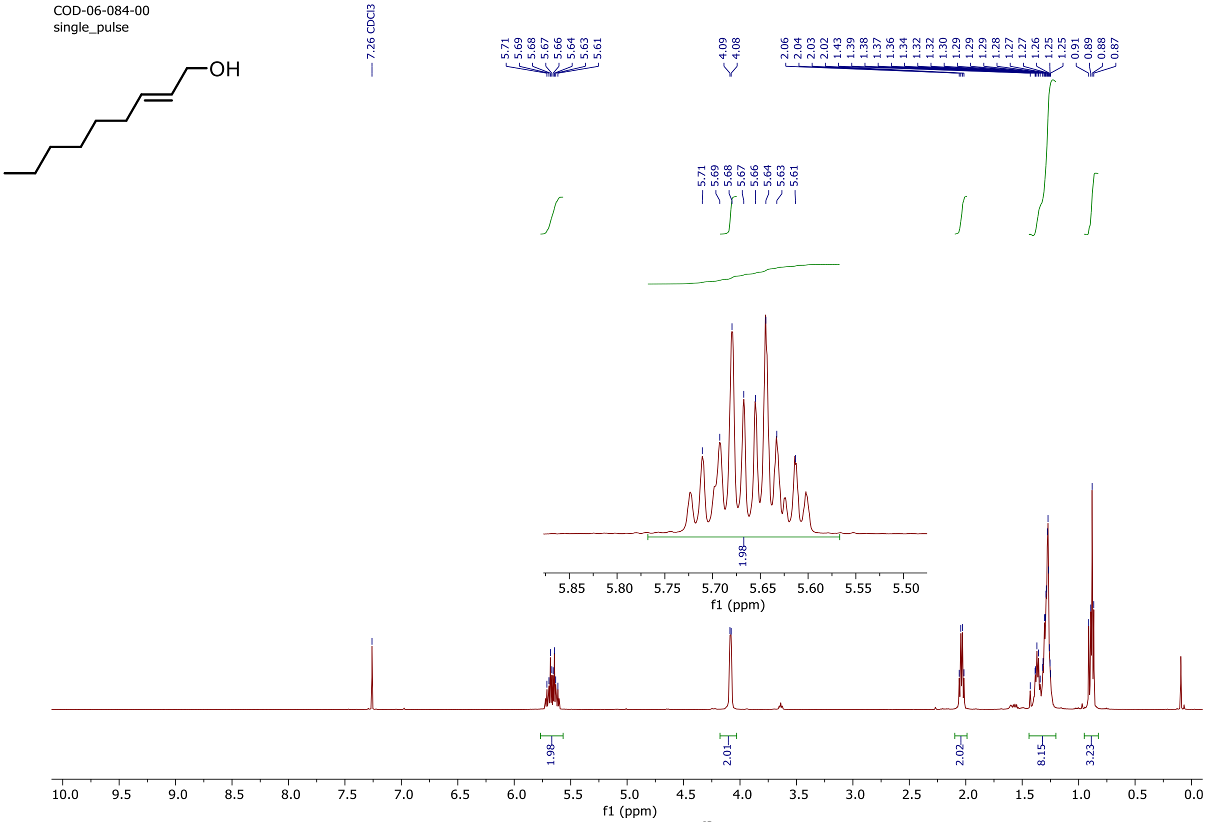

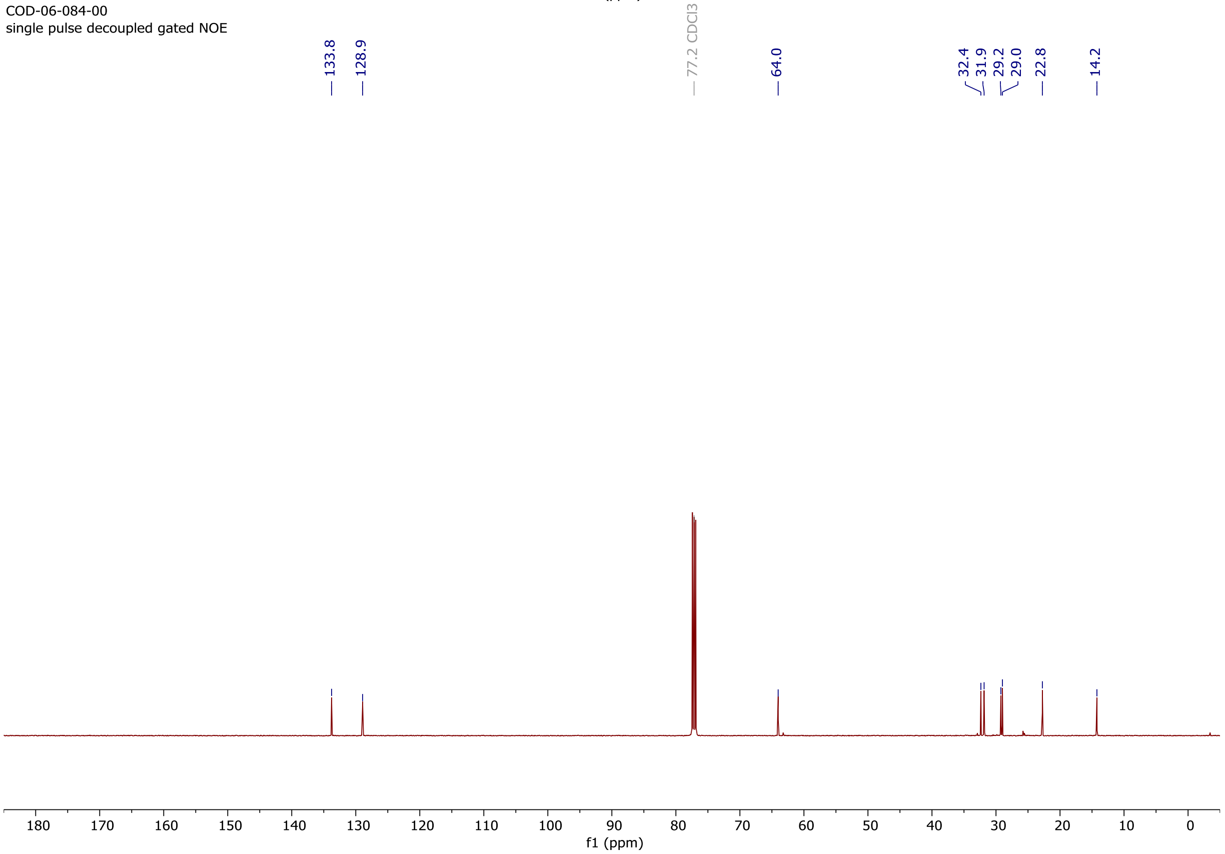

### Copy of $^{1}H$ and $^{13}C\{^{1}H\}$ spectra of **S21**

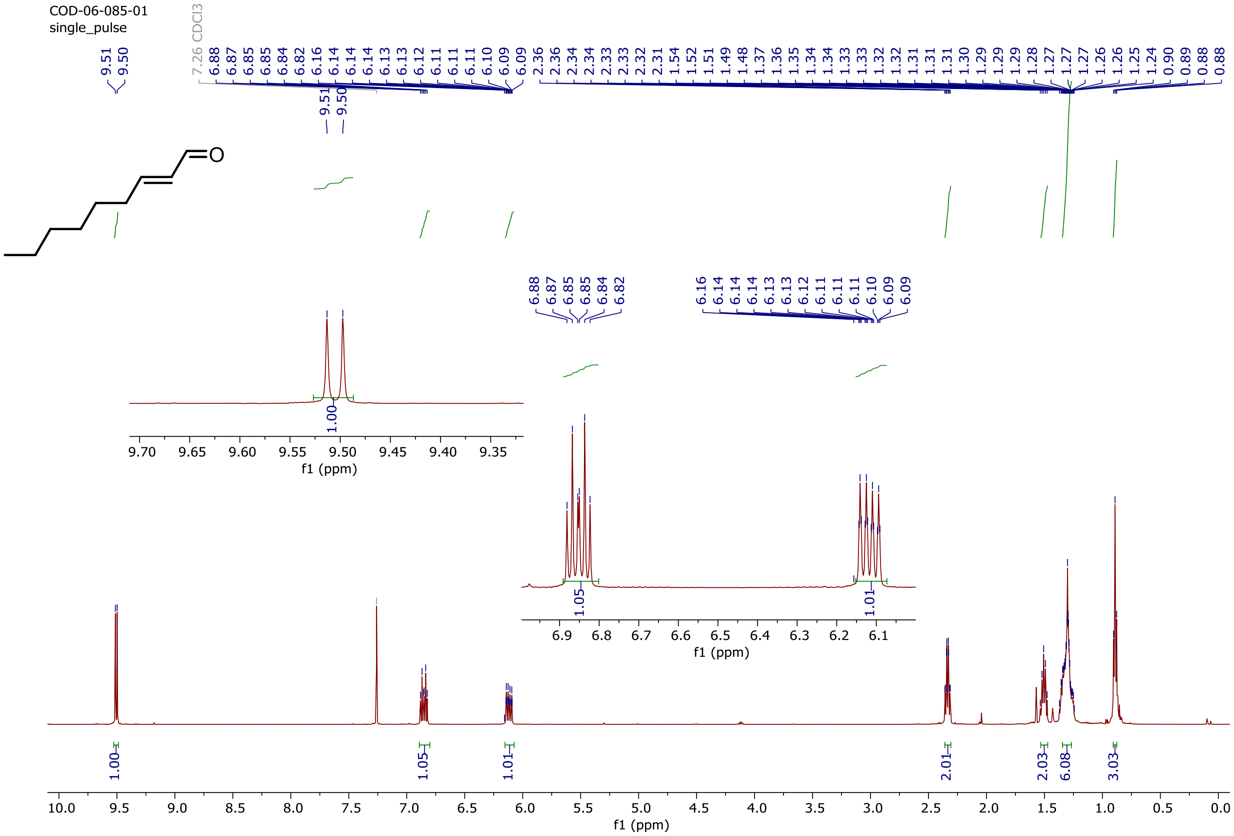

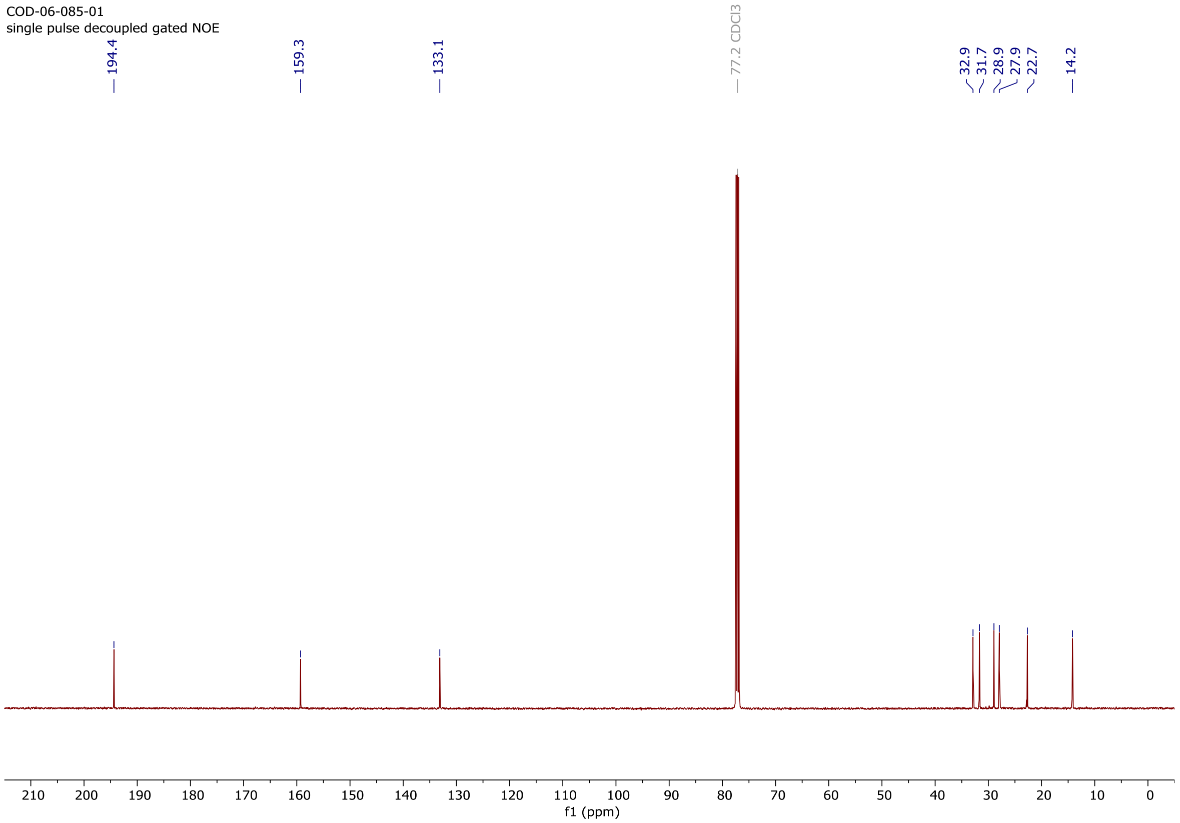

### Copy of $^{1}H$ and $^{13}C\{^{1}H\}$ spectra of **S22**

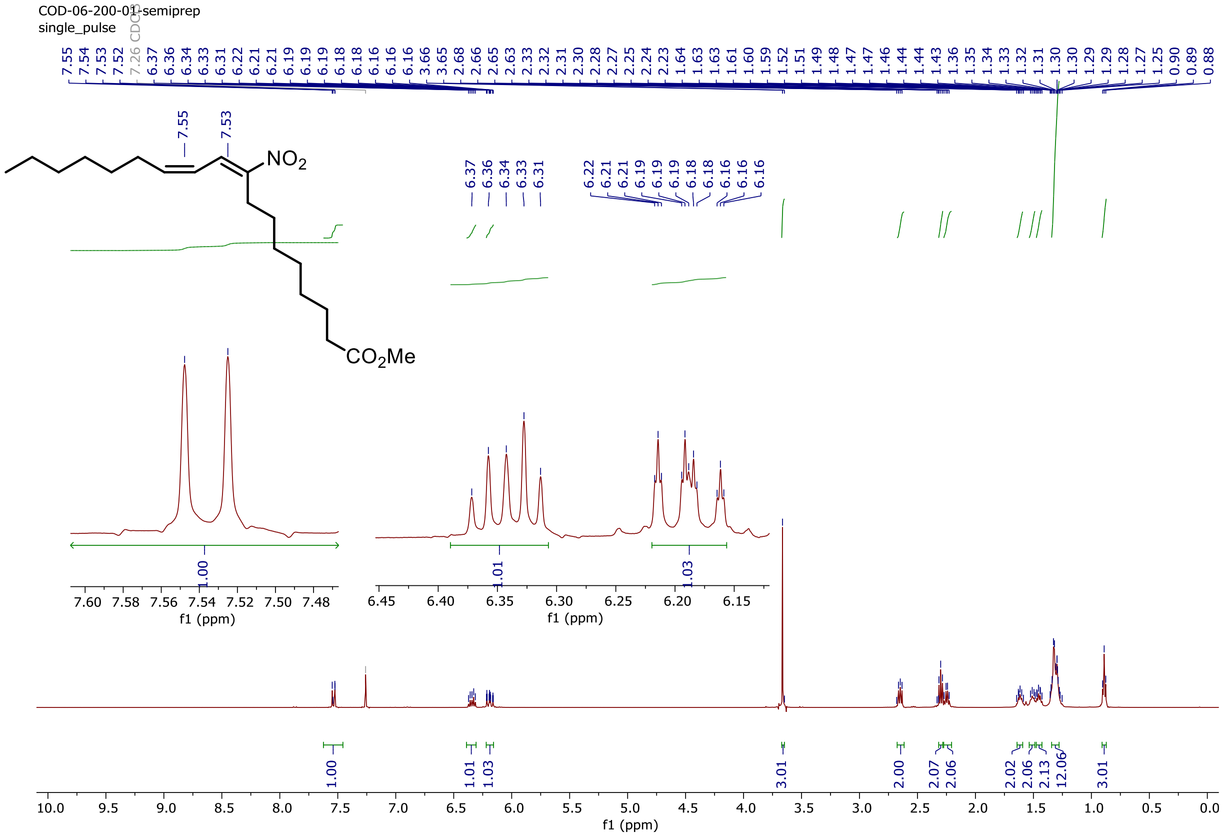

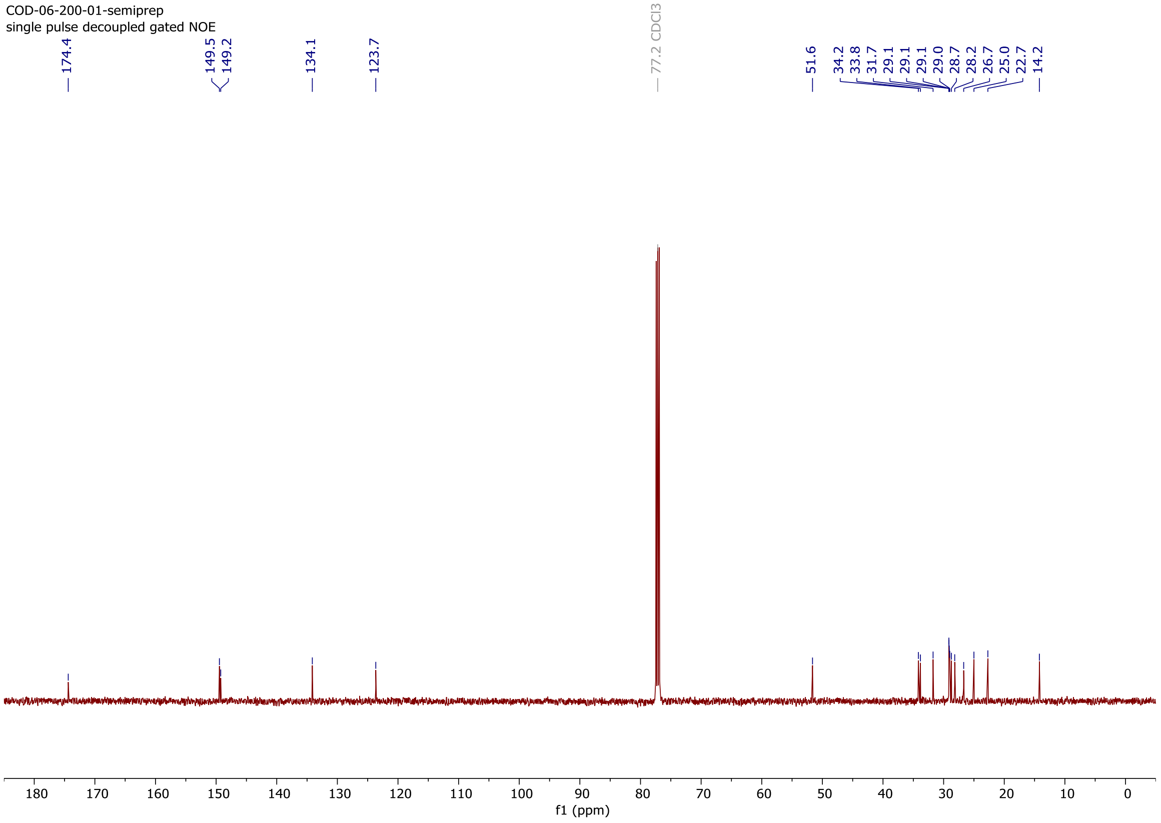

### Copy of $^1$H and $^{13}$C{$^1$H} spectra of **9-NO$_2$cLA**

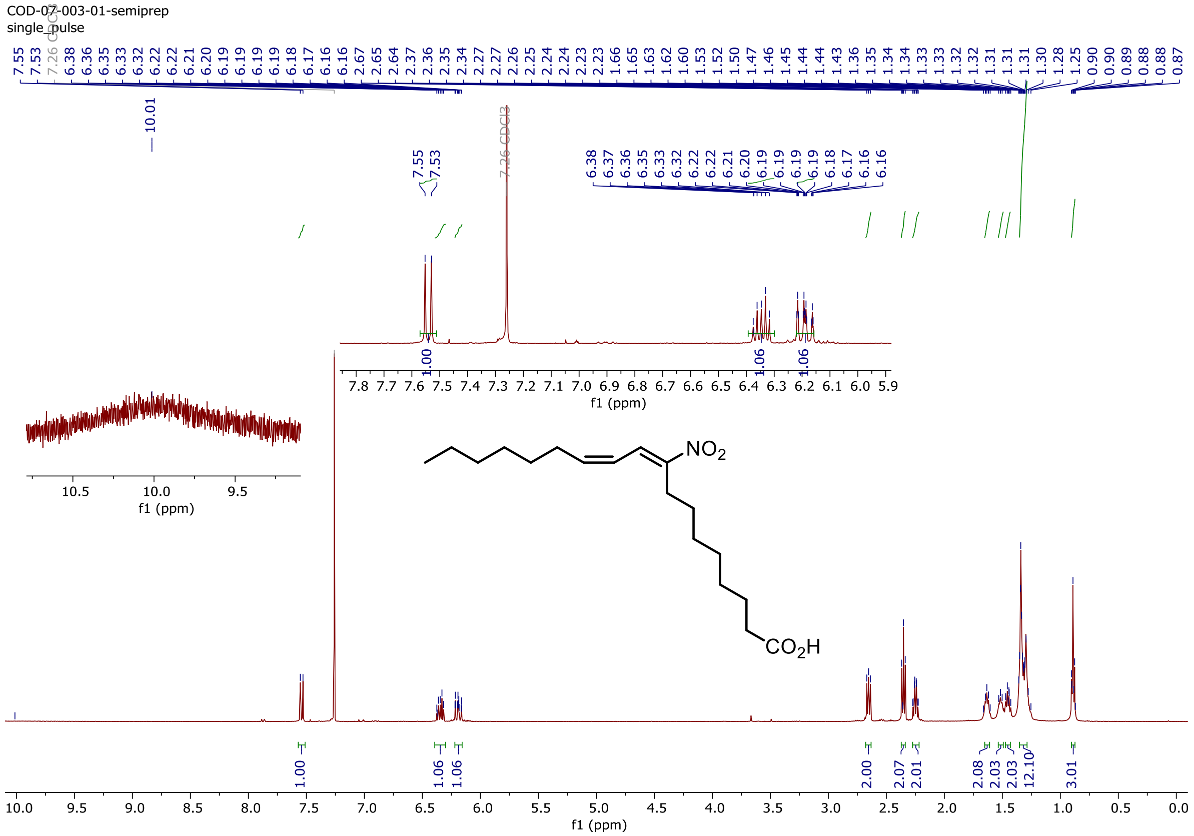

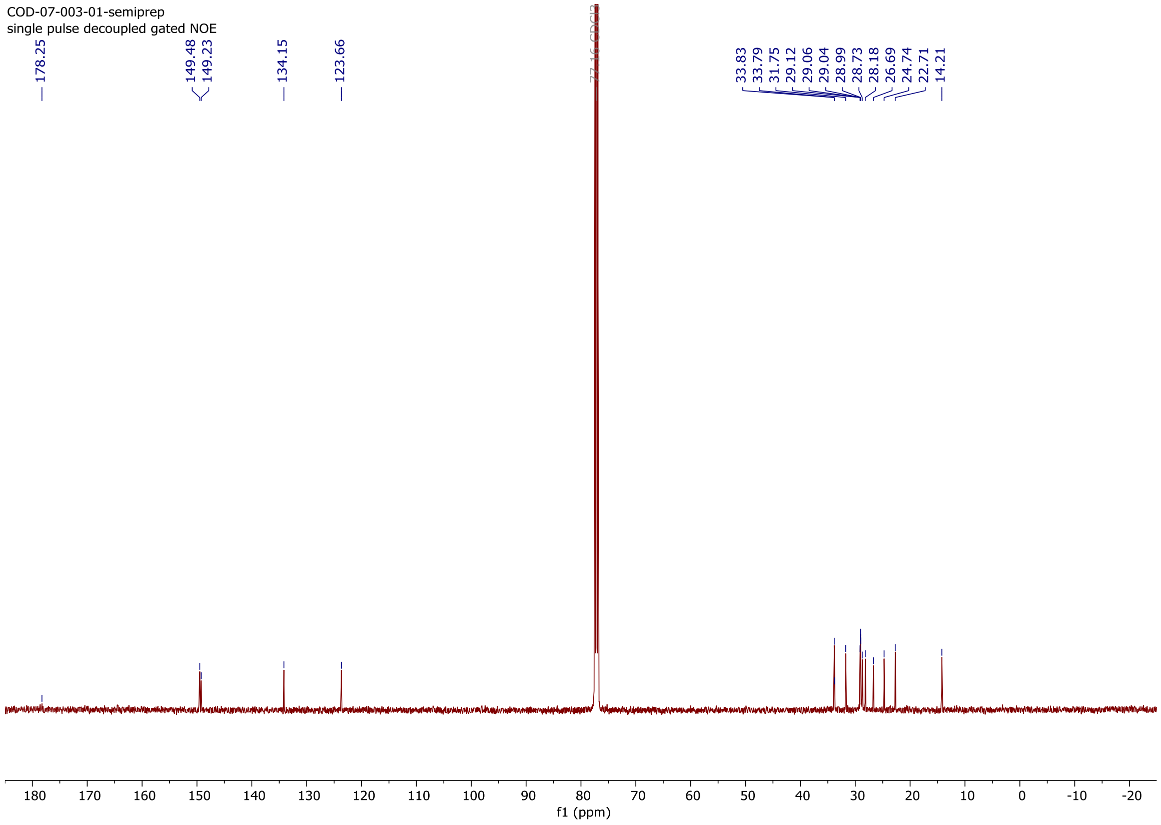

### Copy of $^{1}H$ and $^{13}C\{^{1}H\}$ spectra of **S23**

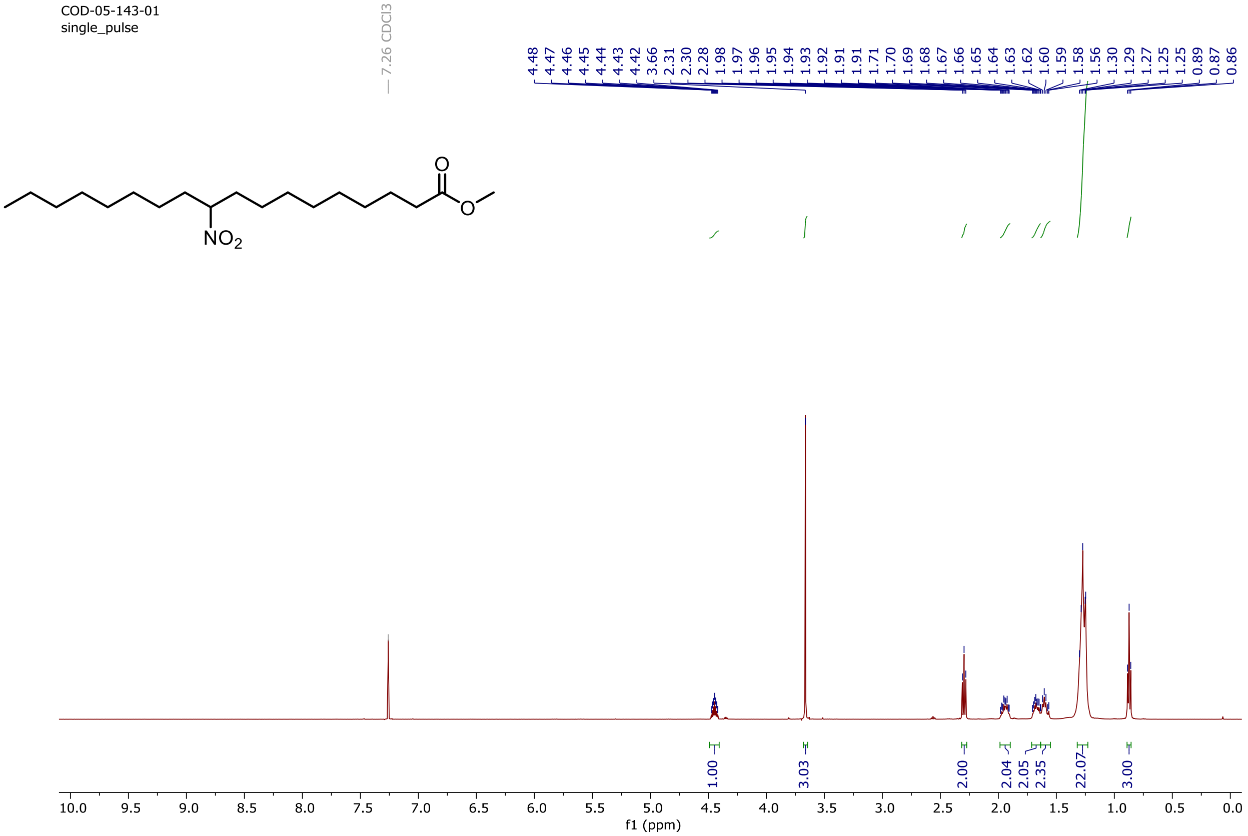

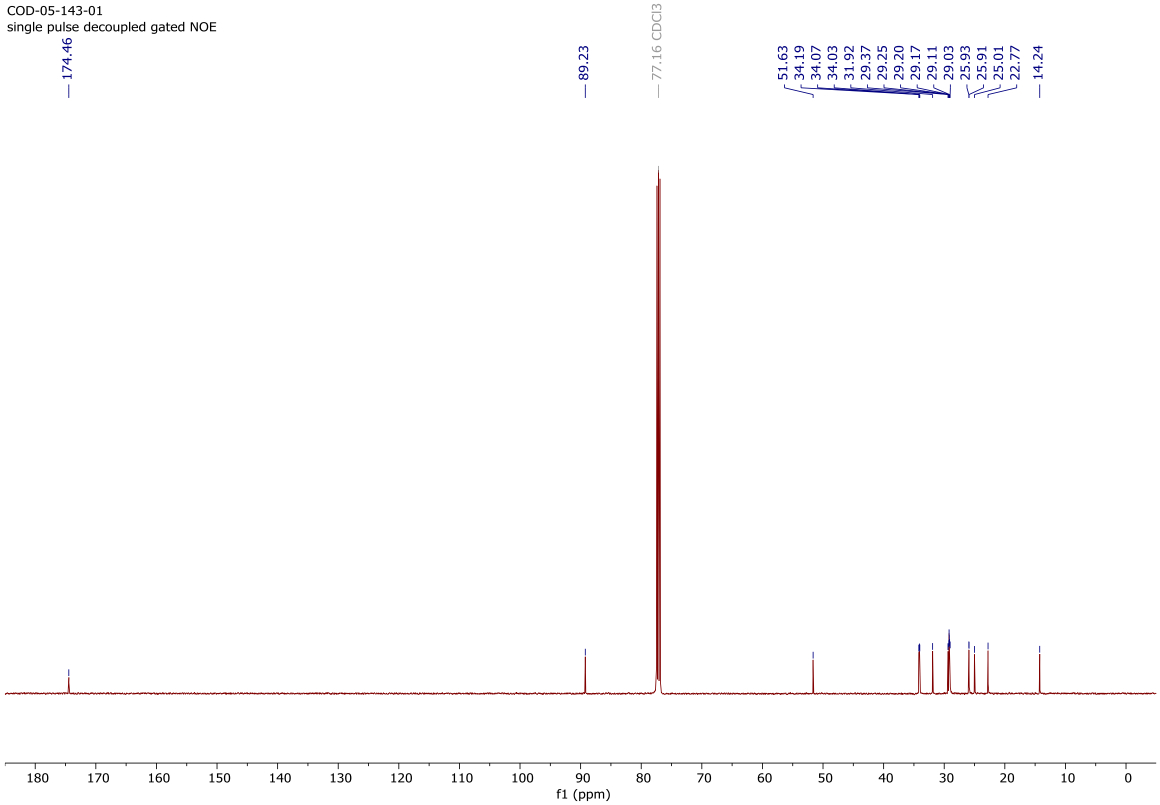

Copy of [1]H and [13]C{[1]H} spectra of **10-nitrostearic acid (10-NO₂SA)**

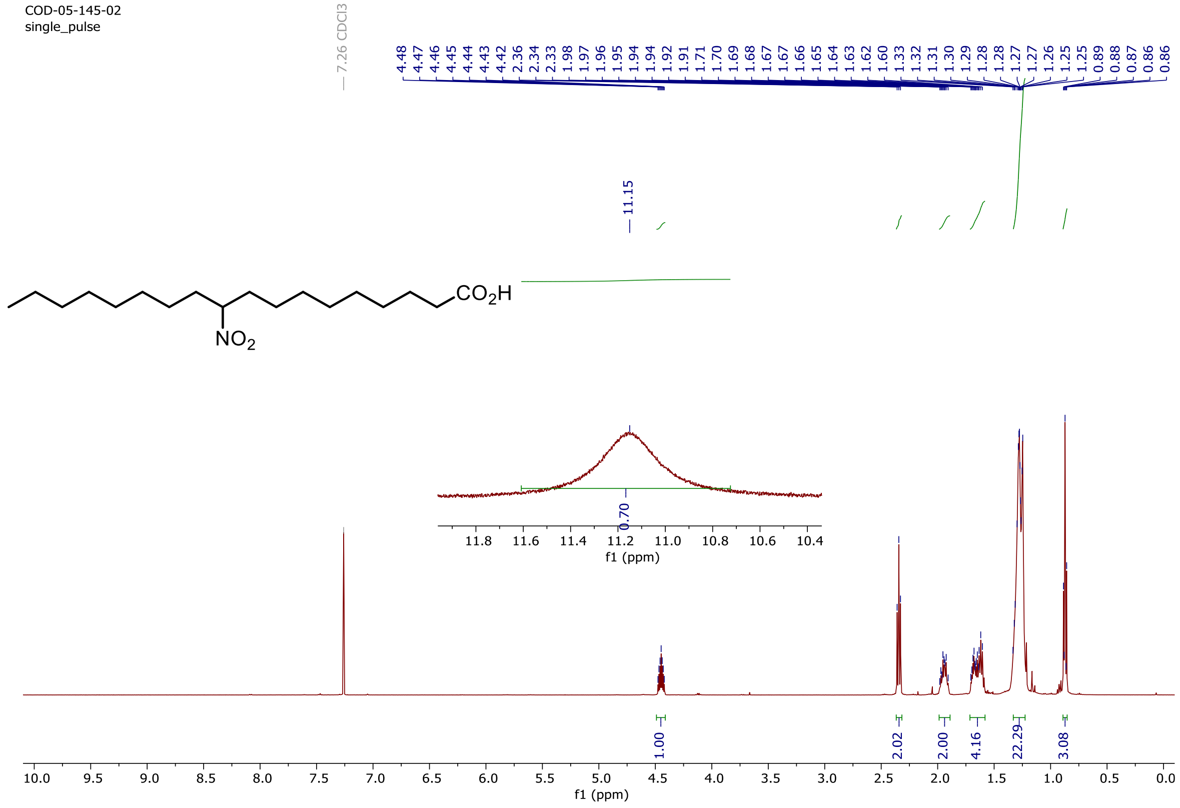

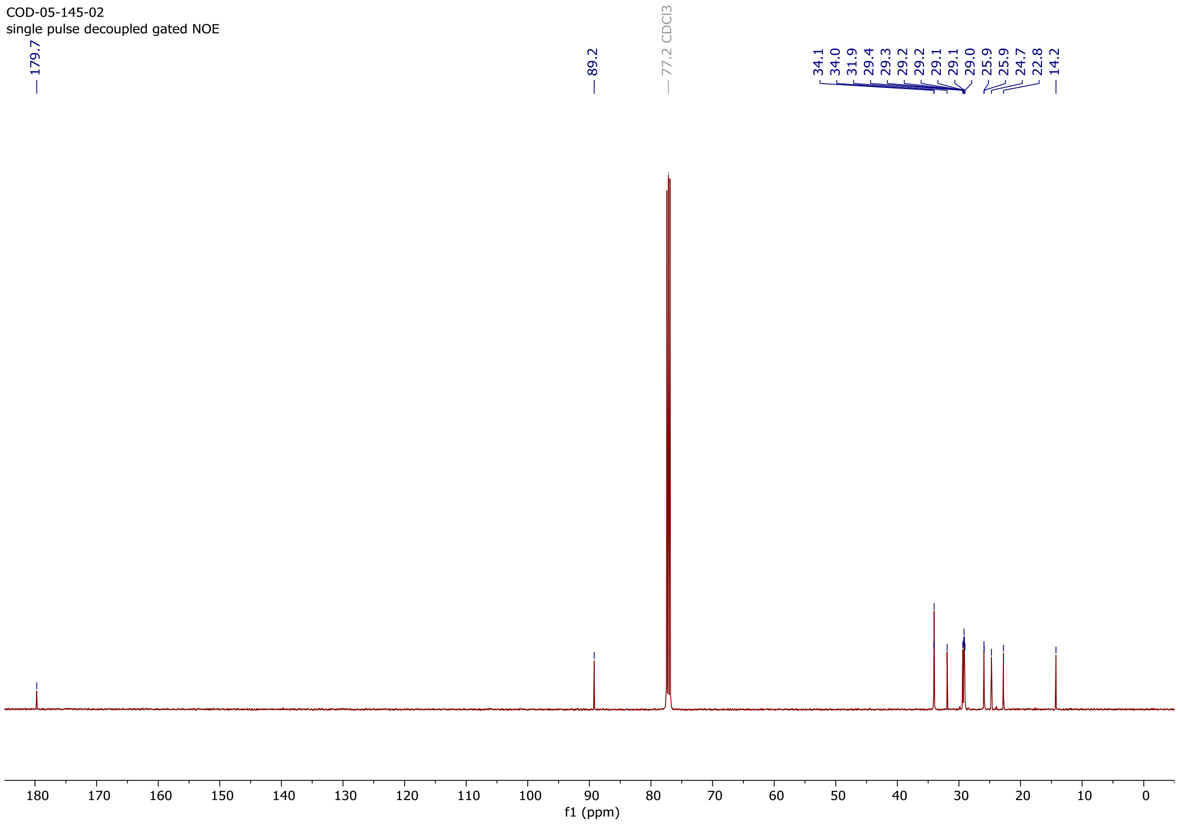
