## Supplementary material for "10-Nitro-oleic acid improves survival following gamma irradiation and modulates hematopoietic and intestinal responses": Publication License Sep-25-2026-graphical abstract

49 Spadina Ave. Suite 200
Toronto ON M5V 2J1 Canada
www.biorender.com

### Confirmation of Publication and Licensing Rights - Open Access

**September 25th, 2026**

| | |
|---|---|
| ***Subscription Type:*** | *Postdoc Plan - Academic* |
| ***Agreement number:*** | *PV2A8XEQXZ* |
| ***Publisher Name:*** | *https://www.biorxiv.org/* |
| ***Figure Title:*** | *Graphical abstract* |
| ***Citation to Use:*** | *Created in BioRender. Pereckova, J. (2026) https://BioRender.com/wxgseko* |

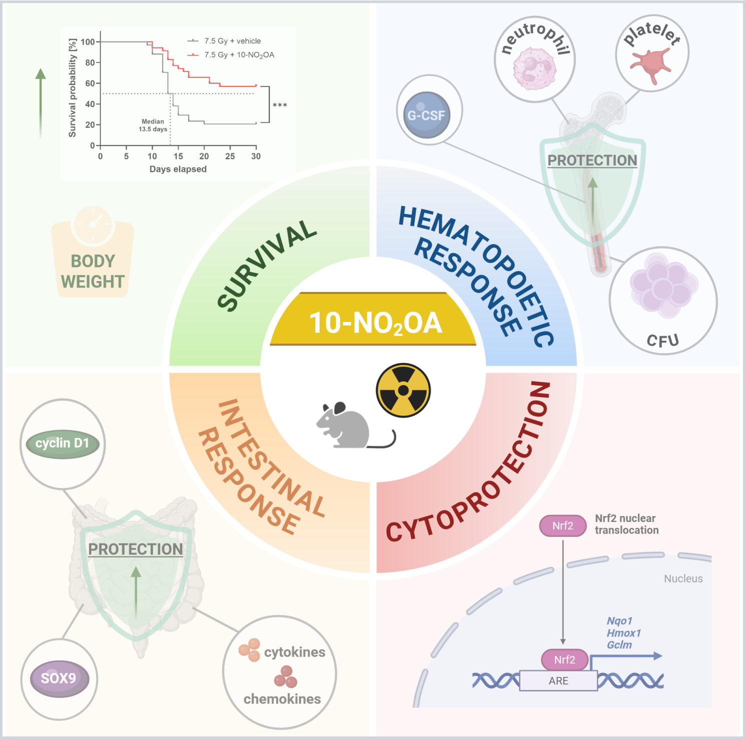


*For any questions regarding this document, or other questions about publishing with BioRender, please refer to our BioRender Publication Guide, or contact BioRender Support at.*
